# Multivariate Random Forests for Cross-Modal Multi-Omics Integration

**DOI:** 10.64898/2026.06.17.732933

**Authors:** Wei Zhang, Lily Wang, Elizabeth J. Franzmann, X. Steven Chen

## Abstract

Multi-omics studies are widely used across many areas of biomedical research. In many diseases, some signals are shared across data types, while others are strongest in a single omics layer. Current multi-omics clustering methods often either merge all data types into a single representation, which can blur biology that is strong in one layer, or rely on linear structure that may miss more complex relationships across data types. We introduce multiRF, a random-forest-based method that handles complex data types and separates shared and modality-specific structure for multi-omics data. multiRF learns sample similarities across omics layers from multivariate random forests, combines them across data types, and uses the resulting weights to estimate the part of each omics layer that is predictable from the others. The remaining residual is treated as modality-specific signal, allowing shared and modality-specific similarities to be clustered separately. In simulations, multiRF recovered shared clusters as well as or better than established integrative methods while more reliably separating modality-specific signal under nonlinear data structures. In TCGA head and neck squamous cell carcinoma, the shared component aligned with the main subtype structure across established reference classifications, while gene- and miRNA-specific components revealed additional immune and developmental biology. In the ADNI cohort with matched blood DNA methylation and structural MRI, the shared cross-modal aging signal was associated with future conversion to mild cognitive impairment or Alzheimer’s disease, and a DNAm-specific residual signal showed exploratory additional information. These results show that multiRF can recover a common disease axis while retaining biologically meaningful signals specific to one data type. multiRF is available as an open-source R package at https://github.com/novawz/multiRF.

## 1 Introduction

Multi-omics studies are now widely used across biomedical research, including cancer, neurodegeneration, immunology, and population health. These studies measure several molecular layers in the same patients, but disease signals are rarely shared equally across data types. Some are consistent across omics layers; others are strongest in a single layer. This split is what makes patient stratification and biomarker discovery from integrated data difficult. Current multi-omics clustering methods tend to fall into two groups. One merges all data types into a single representation, which can blur biology that is strong in one layer. The other relies on linear structure and may miss more complex relationships across data types. Many methods address bulk multi-modal clustering. iCluster [36, 37] and SNF [43] showed that jointly modeling multiple data types can reveal disease subtypes invisible to single-omics analyses. Later latent-variable methods decomposed multi-omics variation into joint and modality-specific components under differing statistical assumptions: AJIVE [13] (common and individual principal-subspace structure), MOFA/MOFA2 [2, 3] (Bayesian group factor analysis), and intNMF [11] (integrative non-negative matrix factorization). DI-ABLO [40] uses sparse partial least squares to identify compact multi-omics signatures across molecular layers. Benchmarks [33, 32] show that these methods perform well on specific cancer-subtyping tasks, though each is limited by data scale, distributional form, or the number of modalities. For single-cell RNA-seq data, weighted nearest neighbors in Seurat v4 [16], deep generative models such as totalVI [14] and MultiVI [4], and graph-linked embedding (GLUE) [10] have advanced multimodal analysis at the cell level, but they depend on single-cell-specific assay structures that do not transfer to typical bulk genomic data.

Graphs are a common way to describe sample relationships in high-dimensional molecular data [42], and the quality of downstream clustering depends directly on how well the graph represents cross-omics similarity. Two recent reviews underline this. Baião et al. [5] classified integration methods by their statistical assumptions, and Wang et al. [44] benchmarked multi-modal clustering algorithms for cancer subtyping, finding that no single method dominates across all scenarios: performance shifts with signal strength, noise level, and the number of modalities. Both reviews argue for flexible methods that make fewer assumptions.

Random forests [9] fit this goal well. Tree splits handle mixed feature types (continuous, binary, count) without requiring a common scale or distributional model across omics layers, where an omics layer is one measured data type such as RNA-seq, miRNA, or DNA methylation. The random subspace procedure [57] provides built-in variable screening in the *p* ≫ *n* regime common to high-throughput data, and the forest-induced proximity (how often two samples share a terminal node) is invariant to monotone feature transformations and robust to outliers and skewed distributions [9, 17]. These properties matter when omics layers follow different underlying distributions. Concretely, each tree partitions samples into terminal nodes, and samples that repeatedly fall into the same node define a local neighborhood. When one omics layer is used to predict another, a multivariate random forest reads cross-omics sample weights off these repeated co-occurrences, and those weights act as a sample-level similarity that records which patients look alike through the lens of cross-omics prediction. Multivariate random forests [35] also accept matrix-valued responses, so one omics layer can predict another and yield a similarity that reflects nonlinear cross-omics structure without specifying a functional form. RF clustering methods continue to develop. Yi et al. [45] proposed an improved affinity estimator with terminal-node reweighting; Bicego [8] introduced dissimilarity-based RF clustering that operates directly on pairwise dissimilarities; Li et al. [27] developed an anchor-graph variant that reduces the cost of proximity construction; and Laska et al. [25] derived closed-form proximity estimators under a treeless RF model. In a biomedical setting, Pfeifer et al. [31] showed that unsupervised RF proximity can be computed in a federated way for privacy-preserving patient stratification. All of these methods, however, operate on a single data matrix and do not address cross-modal fusion.

In prior work [46] we introduced a cross-modal random forest method for biomarker discovery through variable selection, showing that directed cross-modal forests identify biologically relevant features. Here we extend the approach to unsupervised integration and clustering of sample-level bulk multi-modal data. The multiRF (<u>multi</u>-omics integration via <u>R</u>andom <u>F</u>orest) clustering method uses these learned sample weights as the building block for cross-modal similarity. It works differently from latent-factor methods such as AJIVE, MOFA2, or intNMF, which seek a low-dimensional factorization of the data matrices. multiRF instead learns a sample-level cross-omics neighborhood graph, which can capture nonlinear dependence, keep local sample weights, and be split into shared and modality-specific similarities without explicit latent-factor estimation or rank assumptions. The learned cross-omics weights estimate the portion of each omics layer predictable from the others, and the remaining residual is treated as modality-specific signal. A response subsampling step controls the multivariate split criterion for high-dimensional blocks, so the weight matrix is less dominated by high-dimensional noise.

We evaluate multiRF on two simulation benchmarks and two real-data applications. The InterSIM and nonlinear JIVE simulations probe shared-structure recovery, modality-specific signal separation, and stability when the data depart from linear joint-factor assumptions. The first real-data application is cancer subtype discovery in head and neck squamous cell carcinoma (TCGA HNSC), where an unsupervised three-omics integration is compared against three independently derived expert taxonomies (Walter/TCGA, Keck, and the CPTAC-like HPV-negative axes) and the shared and specific components are tested for non-redundant biological and survival signal. The second is conversion to mild cognitive impairment or Alzheimer’s disease in an amyloid- and tau-negative ADNI cohort, where blood DNA methylation and structural MRI are combined to extract a cross-modal biological-aging axis whose association with future conversion is compared with single-modality models and established epigenetic clocks. Together these applications test whether the method recovers meaningful structure across very different biomedical settings.

## 2 Results

Random forests construct an ensemble of decision trees from random subsets of samples and variables. Because each tree recursively partitions samples into terminal nodes, samples that repeatedly co-occur in the same terminal nodes can be treated as similar. Our method builds on this induced similarity structure.

### 2.1 The multiRF clustering method

We developed multiRF, a random-forest-based method for unsupervised integration and clustering of multi-modal omics data (Figure 1). Here, an omics layer means one measured data type such as RNA-seq, miRNA, DNA methylation, or MRI. Given *K* omics layers measured on the same *n* subjects, the method extends the random-forest similarity idea to multi-omics data by fitting multivariate random forests between all pairs of omics data and extracting a sample-level weight matrix from each fit. A response subsampling step controls the split criterion when that response layer is high-dimensional. The learned cross-omics weights are then combined into a global matrix *W_M∗_* and used to estimate the portion of each omics layer that is predictable from the others. The remaining residual is treated as modality-specific signal. This differs from latent-factor methods in the object being learned. Latent-factor methods estimate low-dimensional factors that explain variation in the data matrices. In contrast, multiRF learns a sample-level graph: each edge reflects how strongly one sample helps predict another across omics layers. This graph keeps local, sample-specific relationships and can capture nonlinear dependence without choosing a latent rank. These properties are useful when cross-omics structure is not well described by a small number of linear factors. The resulting shared and modality-specific similarity matrices are then clustered independently. The entire pipeline is implemented in the open-source R package multiRF (https://github.com/novawz/multiRF), which includes a fast implementation of multivariate random forests, the cross-modal variable selection methods from our prior work [46], and the clustering method presented here.

**Figure 1:**
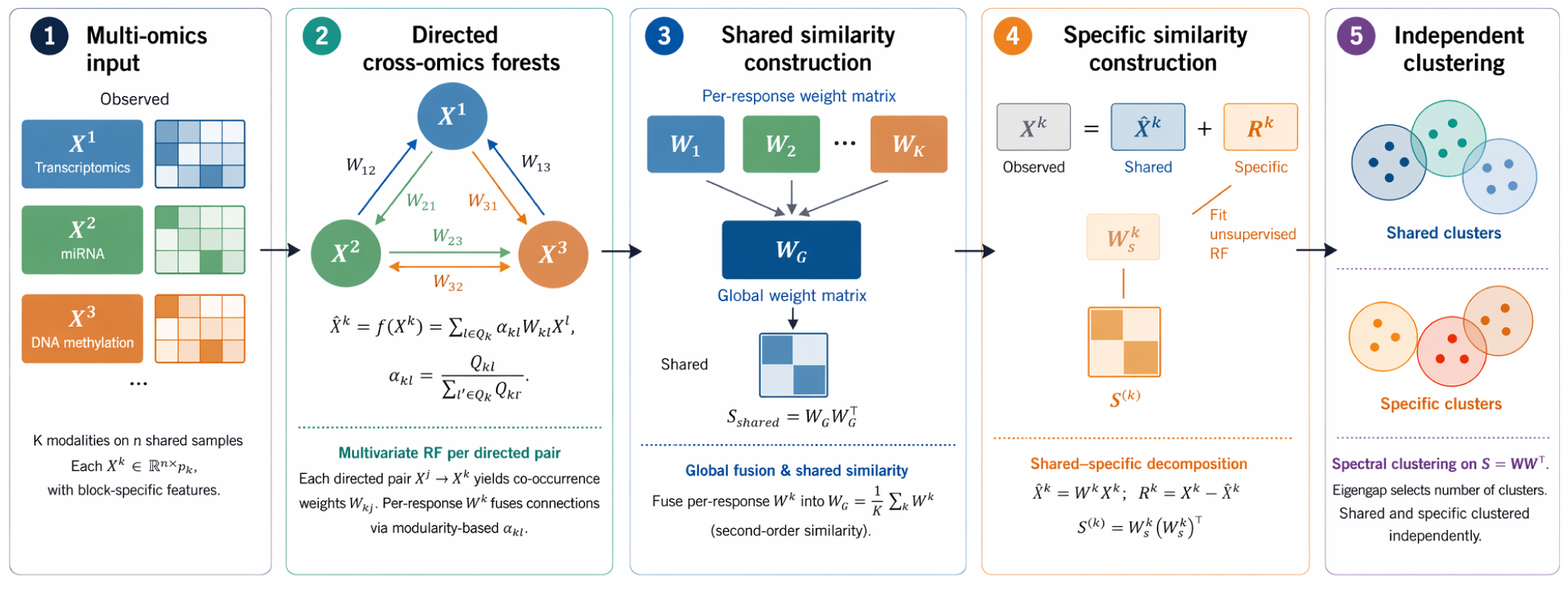
Overview of the multiRF (multivariate random forest) method. For *K* modalities measured on the same *n* subjects, directed cross-omics forests are fit for every predictor–response pair 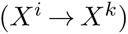, yielding sample co-occurrence weight matrices *W_ki_* Per-response fused weights 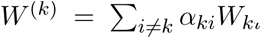 are constructed using modularity-based coefficients 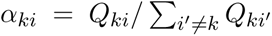, where *Q_ki_* is the Newman modularity of *W_ki_* The global weight matrix 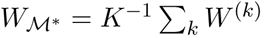 is used to build the shared similarity 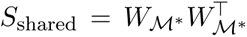, while each *W* ^(*k*)^ drives a cross-omics reconstruction 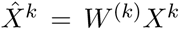 with residual 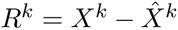 (specific). Spectral clustering is applied independently to the shared and modality-specific similarity matrices.

### 2.2 Simulation Benchmarks

We evaluated multiRF on two simulation studies designed to test shared-structure recovery, shared-specific signal separation, and stability under model mismatch. Each study compares multiRF against baseline integration methods and proximity-based multiRF variants (multiRF-Prox and multiRF-EnhProx; Supplementary Note 3) across multiple signal-to-noise regimes. Full simulation designs are in Sections 4.9.1 and 4.9.2. Data-generating processes, parameter grids, and per-scenario breakdowns are in Supplementary Note 4 and Supplementary Tables 2 and 3.

#### 2.2.1 InterSIM Benchmark

Using InterSIM-generated methylation, gene-expression, and protein profiles [12] with TCGA-derived covariance structures and both shared (*K_Z_*∈ {4, 8}) and modality-specific (*K_U_* = 2) cluster signals, where *K_Z_* denotes the number of shared clusters and *K_U_* the number of modality-specific clusters, the shared signal was present across all three omics layers, while each modality-specific signal was present only in one layer. ARI*_Z_* measured recovery of the shared clusters, whereas ARI*_U_* measured recovery of the modality-specific clusters. For both ARI metrics, values are bounded above by 1, denoting perfect agreement, and values near 0 indicate chance-level agreement. The leakage diagnostic quantified whether a partition intended for one structure also recovered the other structure. It used the same ARI scale, with values near 0 indicating clean separation and larger positive values indicating cross-leakage between shared and modality-specific signals. multiRF achieved near-perfect shared clustering across all scenarios (mean ARI*_Z_* = 0.989), second only to SNF (0.999) and well ahead of block.sPLS (0.976), MOFA2 (0.931), and RGCCA (0.786, Figure 2a, Supplementary Fig. 1, Supplementary Table 3). SNF was strong for shared-structure recovery, but because it returns a fused network rather than an explicit shared-specific decomposition, we did not interpret it as recovering modality-specific clusters in the same way as multiRF. Under weak signal (*δ* = 0.5), multiRF maintained ARI*_Z_* = 0.978, whereas AJIVE dropped to 0.517 and intNMF to 0.427. In the separate response-subsampling sweep, appending 50% pure-noise features produced little loss at the default *q*_try_ = ⌈*q_y_/*3⌉, consistent with the noise-control benefit of response subsampling (Supplementary Note 1; sweep across subsampling fractions in Supplementary Fig. 3). The main advantage of multiRF appeared in specific-structure recovery. It was the only method to reliably detect the injected modality-specific clusters (mean ARI*_U_* = 0.785, Figure 2b). The next-best methods were multiRF-EnhProx (0.344), multiRF-Prox (0.290), and AJIVE (0.200). All remaining baselines produced ARI*_U_* near zero, indicating that their single-partition output mixes shared and specific signals. The leakage diagnostic supported this interpretation, as mean leakage was near zero for multiRF across the 16 InterSIM scenarios, whereas methods that reached high ARI*_Z_* but did not return a separate specific partition showed substantial cross-leakage (mean leakage 0.115 for SNF, 0.135 for RGCCA, 0.112 for mixKernel), indicating that their fused output absorbed shared-cluster structure into what should have been a modality-specific assignment (Figure 2c; Supplementary Table 3 and full results in Supplementary Note 4).

**Figure 2:**
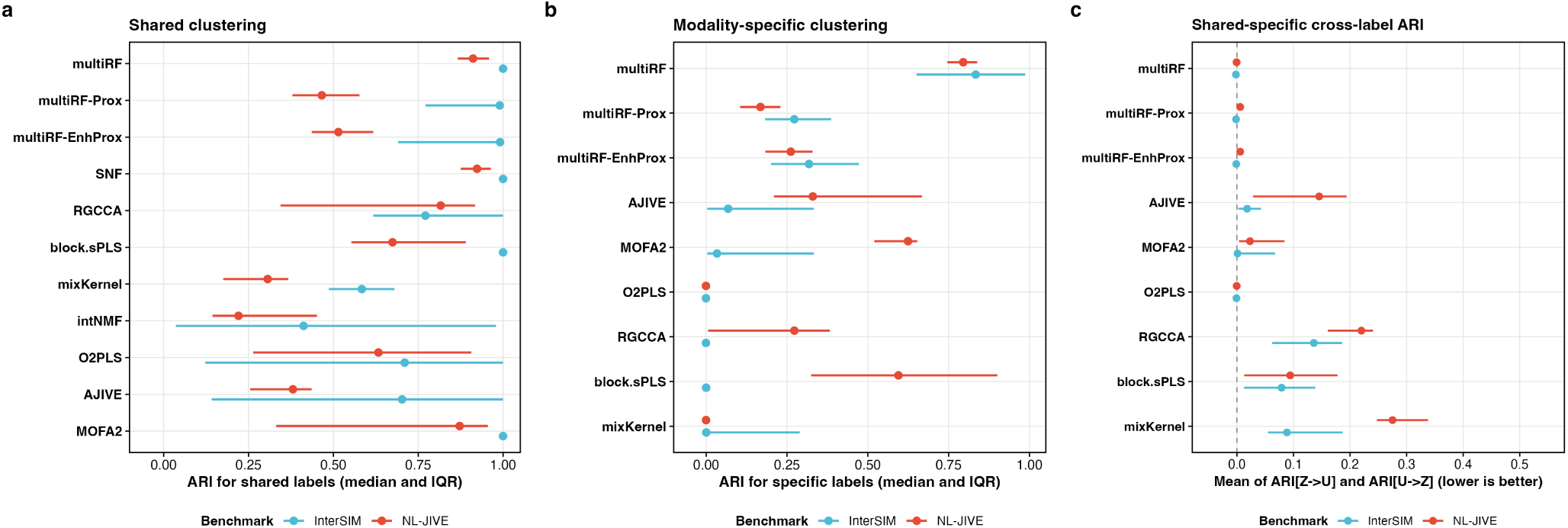
Simulation benchmark results. **a** Shared clustering performance across all benchmarked methods, measured by ARI*_Z_*. **b** Modality-specific clustering performance among methods that return explicit shared and specific components, measured by ARI*_U_*. **c** Cross-label ARI between shared and modality-specific assignments, summarizing ARI*_Z_*_→_*_U_* and ARI*_U_*_→_*_Z_* as defined in Methods. Lower values indicate cleaner separation of the two signal layers. Points show medians across simulation replicates and scenarios, and horizontal bars show interquartile ranges.

#### 2.2.2 Nonlinear JIVE Benchmark

The NL-JIVE simulation applied element-wise nonlinear transformations (*f_J_*∈ {identity, mixed}) to the shared signal in a latent-factor data-generating model, testing stability when the data no longer follow the linear assumptions underlying most factor-based methods. The identity setting is the case in which the data follow the linear structure assumed by factor models. The mixed setting is the nonlinear case: some shared features remain linear, while others are squared before the omics layers are generated. Under this setting (*f_J_*= identity), all factor-based methods performed well, with MOFA2, RGCCA, O2PLS, and multiRF all achieving mean ARI*_Z_* near 0.87 (Figure 2a; per-method per-scenario breakdown in Supplementary Fig. 2). Under the mixed nonlinear setting, the linear methods collapsed. RGCCA fell from 0.869 to 0.248, O2PLS from 0.876 to 0.208, MOFA2 from 0.870 to 0.421, and intNMF from 0.814 to 0.019. multiRF lost only 0.014 (0.869 → 0.855), and SNF showed a similarly modest drop (0.876 → 0.872). This stability is consistent with the ability of tree-based methods to adapt feature by feature without requiring one global linearity assumption. Specific-structure recovery followed the same pattern among methods with explicit shared-specific outputs. multiRF achieved the highest ARI*_U_* in this group. SNF remained strong for shared structure, but its fused network does not provide an explicit residual component, so its per-modality affinities were treated as a comparator rather than as a direct modality-specific cluster recovery result. The same qualitative pattern held in the heterogeneous NL-JIVE setting, where the view-specific signals differed in strength across views: multiRF retained high shared recovery (mean ARI*_Z_* = 0.958), high specific recovery (mean ARI*_U_* = 0.837), and near-zero cross-evaluation leakage, whereas several baselines either lost specific recovery or showed higher leakage (full results in Supplementary Table 3 and Supplementary Note 4).

#### 2.2.3 Runtime

Runtime on both simulation benchmarks was similar to the fastest latent-variable baseline and clearly ahead of graph- and factorization-based alternatives. On the InterSIM setting with *n* = 500 samples across three blocks, multiRF ran in roughly the same time as MOFA2 and in under half the time required by SNF or intNMF. On the larger NL-JIVE setting with *n* = 1,000 and up to 500 features per view, multiRF was about twice as fast as MOFA2 (Supplementary Table 3). These runtime results indicate that the shared-specific decomposition does not impose a major runtime cost. multiRF delivers the accuracy gains reported above at a runtime comparable to the strongest integrative baseline we tested.

### 2.3 TCGA HNSC: Cancer Subtype Discovery

Head and neck squamous cell carcinoma (HNSCC) is a demanding test of shared–specific decomposition because its main subtype structure is reproducible across cohorts, whereas important biology is stronger in some data types than in others. Molecular taxonomy in HNSCC has progressed from early expression-defined subtypes [47, 58], through exome-scale genomic characterization of recurrent driver alterations [48, 49], to the multi-platform TCGA atlas [50] and later integrative refinements [59, 60]. These reference systems describe a similar disease pattern from different data types. By contrast, immune infiltration is most visible in RNA expression, whereas imprinted and fetal-associated miRNA variation is most visible in miRNA. HNSCC therefore lets us ask two questions at once: whether the shared component aligns with established reference classifications, and whether the modality-specific components add information beyond that shared subtype pattern.

We applied multiRF to 517 HNSCC tumors from TCGA with matched RNA-seq, miRNA-seq, and Illumina 450K methylation profiles (5,000 most variable genes and CpGs, all 525 miRNAs passing quality control filters), linking all six directed cross-omics connections without any subtype label. The real-data cohorts and modality dimensions are summarized in Table 1. The method jointly estimates a *shared* partition that reflects the main cross-modal structure and *modality-specific* residual components that reflect signal unique to each data type. Walter/TCGA centroid calls were available for 279 patients (36 HPV-positive, 243 HPV-negative), and nine recurrently altered drivers (*TP53*, *CDKN2A*, *CASP8*, *NOTCH1*, *HRAS*, *PIK3CA*, *NFE2L2*, *KEAP1*, *CUL3*), copy-number burden, anatomic subsite, AJCC stage, and overall survival provide the biological and clinical readouts used below. We organize the HNSCC results around four questions: shared subtype alignment, HPV-negative CPTAC-like axes, gene-specific immune microenvironment, and miRNA-specific developmental biology (Figures 3, 4, and 5). Survival models are then used as a secondary test of whether these layers associate with survival information.

**Figure 3:**
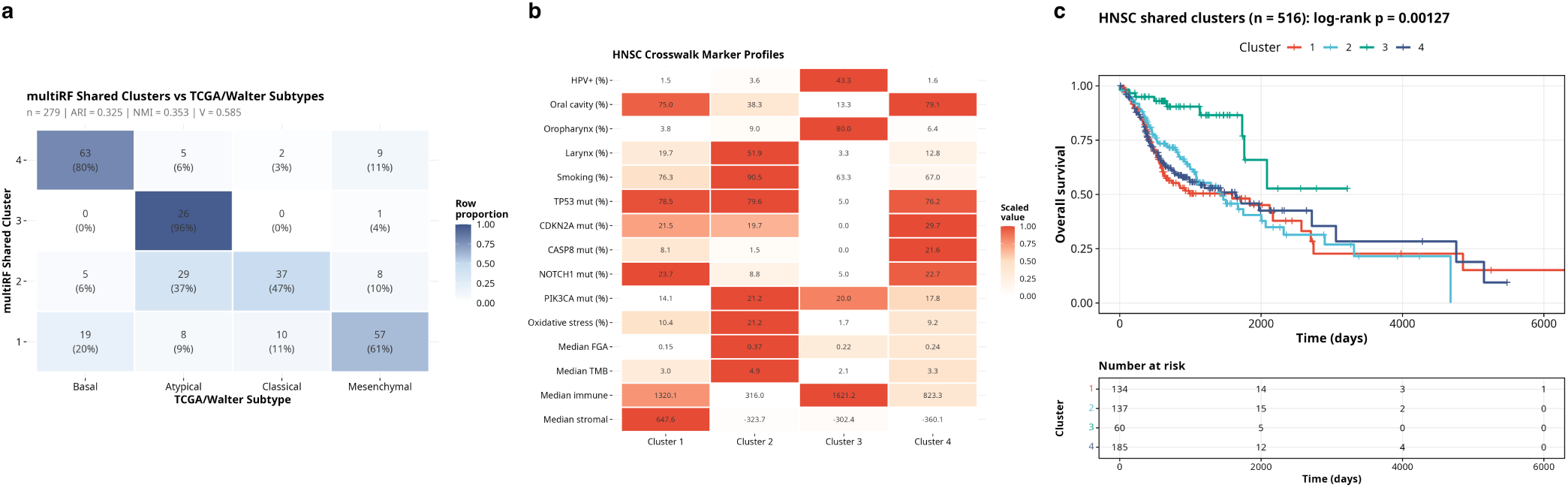
TCGA HNSC shared-cluster discovery and clinical association (*n* = 517 after QC; *n* = 279 with curated Walter/TCGA labels). **a** Contingency heatmap of the four multiRF shared clusters against centroid-based Walter/TCGA subtypes, annotated with row proportions and cell counts (ARI = 0.325, NMI = 0.353, Cramér’s *V* = 0.585, *χ*^2^ *p* = 1.9 × 10^−56^). **b** Crosswalk marker profile of each shared cluster across HPV status, anatomic subsite (oral cavity / oropharynx / larynx), smoking, canonical HNSCC drivers (TP53, CDKN2A, CASP8, NOTCH1, PIK3CA), NFE2L2–KEAP1–CUL3 oxidative-stress alterations, chromosomal instability (median FGA / TMB), and ESTIMATE immune / stromal scores. **c** Kaplan–Meier curves for overall survival across the four shared clusters (log-rank *p* = 1.3 × 10^−3^), with per-cluster risk tables.

**Table 1:** Summary of real-data applications.

|  | TCGA HNSC | ADNI |
| --- | --- | --- |
| Subjects ($n$) | 517 | 87 |
| Modalities ($K$) | 3 | 2 |
| Modality 1 | mRNA (5,000 genes) | Structural MRI (85 features) |
| Modality 2 | DNAm (5,000 CpGs) | Blood DNAm (4,332 CpGs) |
| Modality 3 | miRNA (full panel) | — |
| Outcome | Overall survival, PFI | MCI/AD conversion |
| Evaluation | Log-rank, $C$-index | Cox HR, $C$-index |
| Feature selection | Top 5,000 by MAD (mRNA, DNAm); all miRNAs | Brain-blood + AD-prioritized CpGs |
| Baseline status | Tumor at diagnosis | Cognitively normal (A–T–) |

#### 2.3.1 Shared Clustering Aligns with Walter/TCGA Subtypes

We selected four clusters using the shared similarity matrix on 517 HNSCC tumors derived from multiRF, and unsupervised assignment aligned with the four canonical Walter/TCGA subtypes without seeing any subtype label (Figure 3a, Table 2, per-pair tests and agreement statistics in Supplementary Table 4). Each cluster was dominated by one TCGA subtype, and the subtype agreement was within the range typically used to support reproducible HNSC subtype recovery. Cluster-level marker and phenotype profiling then showed that this shared partition corresponds to the known HNSCC disease axis rather than to an arbitrary clustering of mixed features (Figure 3b, per-cluster tests and full marker heatmap in Supplementary Fig. 4 and Supplementary Table 5). The Atypical-like cluster was the HPV-driven, oropharyngeal, *TP53* -wildtype group. The Classical-like cluster captured the heavy-smoker, laryngeal, copy-number-high phenotype with *NFE2L2* /*KEAP1* /*CUL3* oxidative-stress hits. The Basal-like cluster concentrated *CASP8*, *HRAS*, and *CDKN2A* lesions with a strong EGFR-ligand and keratinization program. The Mesenchymal-like cluster combined the highest stromal/immune infiltration with the strongest EMT signature at near-zero HPV positivity. The same partition was concordant with the Keck five-subtype classification and, for the three mostly HPV-negative clusters, with CPTAC-like axes (Table 2, signature scores and per-cluster axis profiles in Supplementary Fig. 5 and Supplementary Table 6). Cluster 3 corresponded to HPV-Immune in the Keck classification, cluster 1 to Mesenchymal, cluster 2 to Classical, and cluster 4 to Basal. Because the CPTAC reference classification was derived exclusively from HPV-negative tumors [60], only the three mostly HPV-negative clusters have CPTAC-like counterparts. Cluster 2 aligned with the CIN-like axis, Cluster 4 with Basal-like, and Cluster 1 with Immune-like. The HPV-positive Cluster 3 has no direct CPTAC counterpart. The shared component is therefore the main HNSCC subtype axis. It aligns with the Walter/TCGA four-way expression classification, remains interpretable under the Keck system, and, after restricting to HPV-negative disease, continues to align with CPTAC-like proteogenomic axes, reflecting the common structure these independently derived reference systems were each built to describe.

**Table 2:**
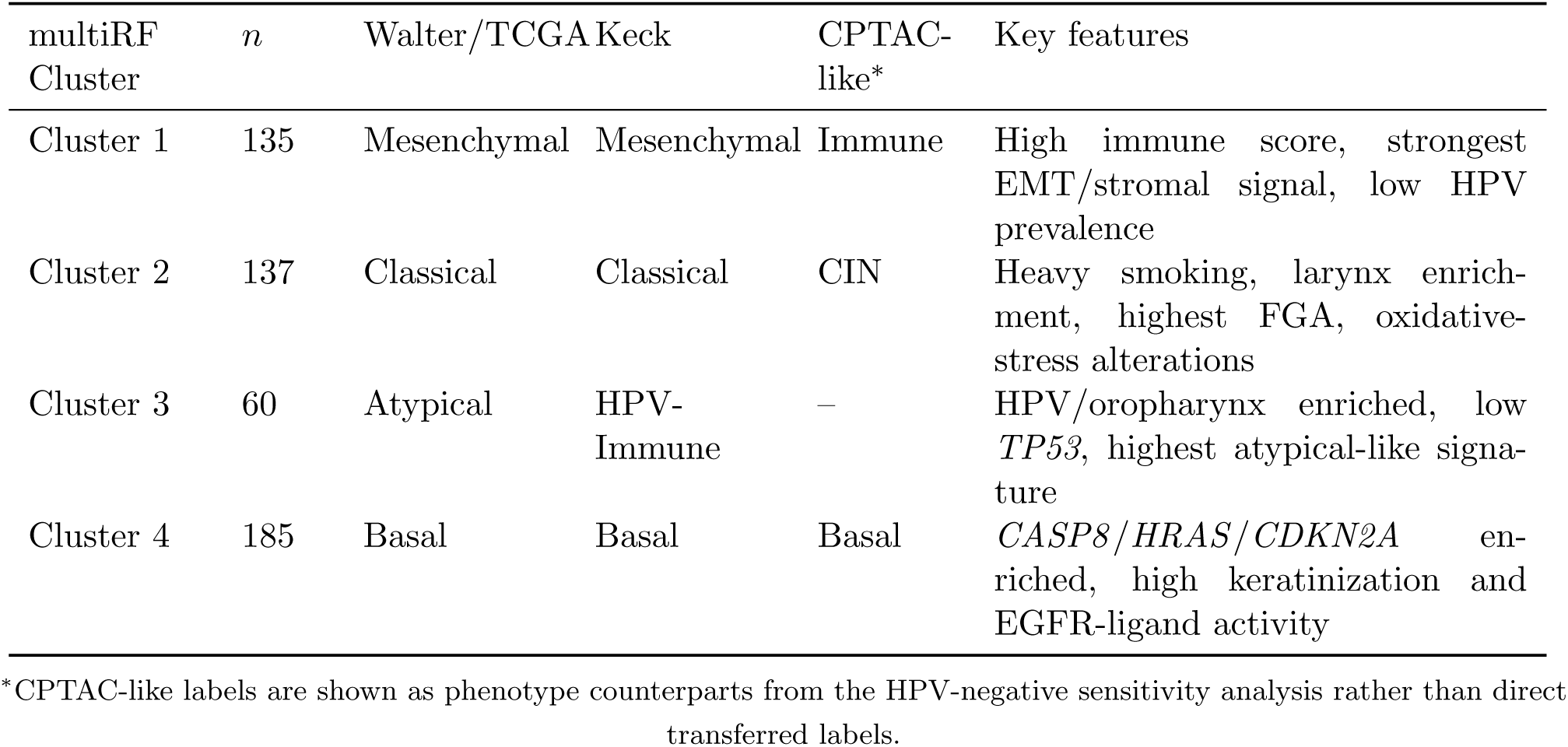
HNSC alignment of multiRF shared clusters to established subtype systems. Each row summarizes one shared cluster using the main centroid-based Walter/TCGA label, the most likely Keck assignment, the corresponding HPV-negative CPTAC-like counterpart, and the canonical biological features supporting that mapping.

Beyond alignment with the shared subtype structure, the four shared clusters separated overall survival among patients with available OS follow-up (log-rank *p* = 1.3 × 10^−3^, *n* = 516, Figure 3c). Bootstrap re-sampling and random-seed reinitialization further showed that the shared partition was highly reproducible (Supplementary Fig. 6). Benchmarking against four alternative integrative methods on the identical feature panel (Table 3) showed that multiRF had the best overall balance across subtype agreement, survival separation, and survival discrimination, with full method-by-method comparisons deferred to the Supplementary Results.

**Table 3:** HNSC comparison across integrative clustering methods. The table distinguishes survival separation (log-rank *p*, *C*-index) from agreement with the curated Walter/TCGA subtype geometry (ARI). iClusterPlus has the highest ARI (0.332), with multiRF a close second (0.325, a difference of 0.007); mul-tiRF combines this near-best subtype agreement with the strongest survival separation among the methods compared (log-rank *p* = 1.3 × 10^−3^, tied with SNF, and *C* = 0.574, 0.007 below SNF’s 0.581). SNF is therefore slightly stronger on survival ranking but markedly weaker on ARI (0.267), and iClusterPlus is slightly stronger on ARI but substantially weaker on survival (*p* = 2.3 × 10^−2^, *C* = 0.552). multiRF therefore offers the most balanced profile across the two evaluation axes.

| Method | Log-rank $p$ | Concordance Index ($C$) | ARI vs centroid subtype |
| --- | --- | --- | --- |
| multiRF shared | $1.3 \times 10^{-3}$ | 0.574 | 0.325 |
| SNF fused | $1.3 \times 10^{-3}$ | 0.581 | 0.267 |
| MOFA2 shared | $5.7 \times 10^{-3}$ | 0.571 | 0.237 |
| iClusterPlus | $2.3 \times 10^{-2}$ | 0.552 | 0.332 |
| AJIVE joint | $1.4 \times 10^{-2}$ | 0.568 | 0.301 |

#### 2.3.2 HPV-Negative Analysis and CPTAC-Like Axes

Because HPV status reshapes HNSC driver biology, immune microenvironment, and prognosis, and because the CPTAC proteogenomic subtypes were explicitly defined on HPV-negative tumors [60], we repeated the analysis in the HPV-negative subset as a biological sensitivity analysis (Section 4.10.3). This restricted analysis is a stricter test than the full-cohort version. With the large HPV-positive group removed, the shared component can no longer rely on the HPV/non-HPV contrast, so any structure it still identifies must reflect biology beyond HPV status. Clustering analysis supported two primary modes but placed the second-largest gap at *k* = 3, which we adopted so that the partition can be interpreted against the three CPTAC-like HPV-negative axes.

The three HPV-negative clusters separated along the CPTAC-like signature axes (Figure 5a,b). One cluster was copy-number high, laryngeal-enriched, and heavy smoker-dominated, mapping to CIN-like. A second loaded on EGFR-ligand and basal-differentiation markers with oral-cavity predominance, mapping to Basal-like. A third was immune-infiltrated and checkpoint-high, mapping to Immune-like. These three states are not a trivial relabeling of the original four clusters. Rather, they show that after removing the HPV-driven Atypical group, the remaining shared structure remains consistent with the basal, chromosomal-instability, and immune-related axes reported by CPTAC from an independent proteogenomic analysis.

#### 2.3.3 Gene-Specific Component Captures Immune Microenvironment

Once the shared subtype axis had been removed, the leading gene-specific axis tracked immune infiltration more strongly than any shared-component dimension (Figure 4a,b, full statistics in Supplementary Table 7). The eigenvector was strongly positive with ESTIMATE immune and stromal scores and strongly negative with tumor purity, and cell-type deconvolution mapped this signal primarily to immune infiltration, with contributions from dendritic cells, B cells, M1 macrophages, and CD8+ T cells. Signature-level analysis sharpened this: the axis aligned with T-cell/cytotoxic, interferon-*γ*, and antigen-presentation programs, but not with keratinization or EGFR-ligand activity. These targeted bulk signatures overlap with programs emphasized by single-cell HNSCC studies, including partial EMT/stromal, epithelial-differentiation, stress/hypoxia, cell-cycle, and tumor-microenvironment programs [61]. The gene-specific component is thus a second layer of HNSCC variation that sits on top of the shared subtype structure rather than replacing it: the shared component organizes the major etiologic and epithelial-state axis, while the gene-specific component reflects a modality-specific immune-microenvironment signal that the shared clustering does not isolate.

**Figure 4:**
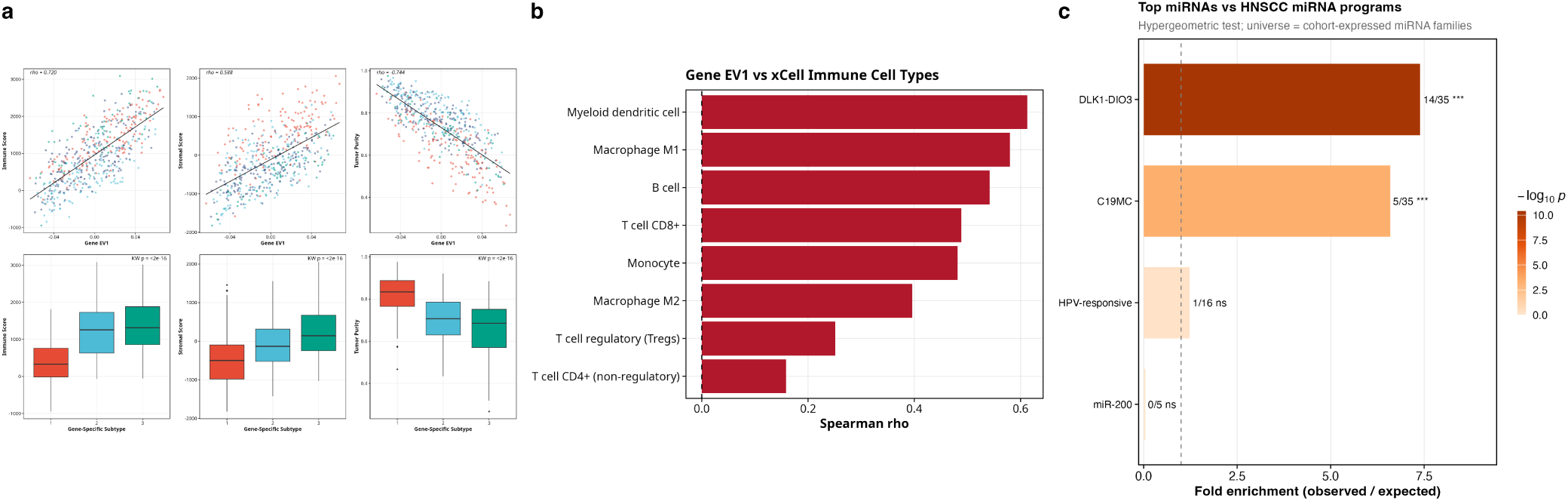
Gene and miRNA residual biology revealed after removing the shared cross-omic structure. **a** Gene-specific EV1 versus ESTIMATE immune score, ESTIMATE stromal score, and tumor purity, color-coded by shared cluster; lower panels show the distribution of each score across the four shared clusters. Gene EV1 tracks immune infiltration (*ρ* = 0.72), stromal content (*ρ* = 0.59), and lower tumor purity (*ρ* = −0.74). **b** Spearman correlations of gene EV1 with xCell deconvolution estimates for eight immune cell populations; myeloid dendritic cells (*ρ* = 0.61) and M1 macrophages (*ρ* = 0.58) load most strongly. **c** Hypergeometric fold-enrichment of top miRNA-specific features against known HNSCC miRNA programs (DLK1–DIO3 imprinted cluster, C19MC, HPV-responsive and miR-200 programs), with BH-corrected *q*-values annotated.

These associations held after spectral re-clustering of the gene-specific similarity matrix and in sensitivity analyses restricted to HPV-negative tumors or adjusted for stage, subsite, and purity (Supplementary Fig. 7). With the shared-cluster results, this supports a layered view of HNSCC: the shared partition reflects the main disease axis, while the gene-specific axis adds immune variation that the shared subtype structure does not resolve. This immune axis also carried a modest overall-survival association in the full clinical Cox model (gene EV1 HR = 0.82, *p* = 0.03; Table 4). However, in the full clinical sensitivity models this association was attenuated after adding tumor purity, ESTIMATE immune score, or ESTIMATE stromal score (HR = 0.85–0.89, all *p >* 0.28; Supplementary Table 14), indicating that the gene-specific survival signal is largely tied to microenvironmental composition.

**Table 4:**
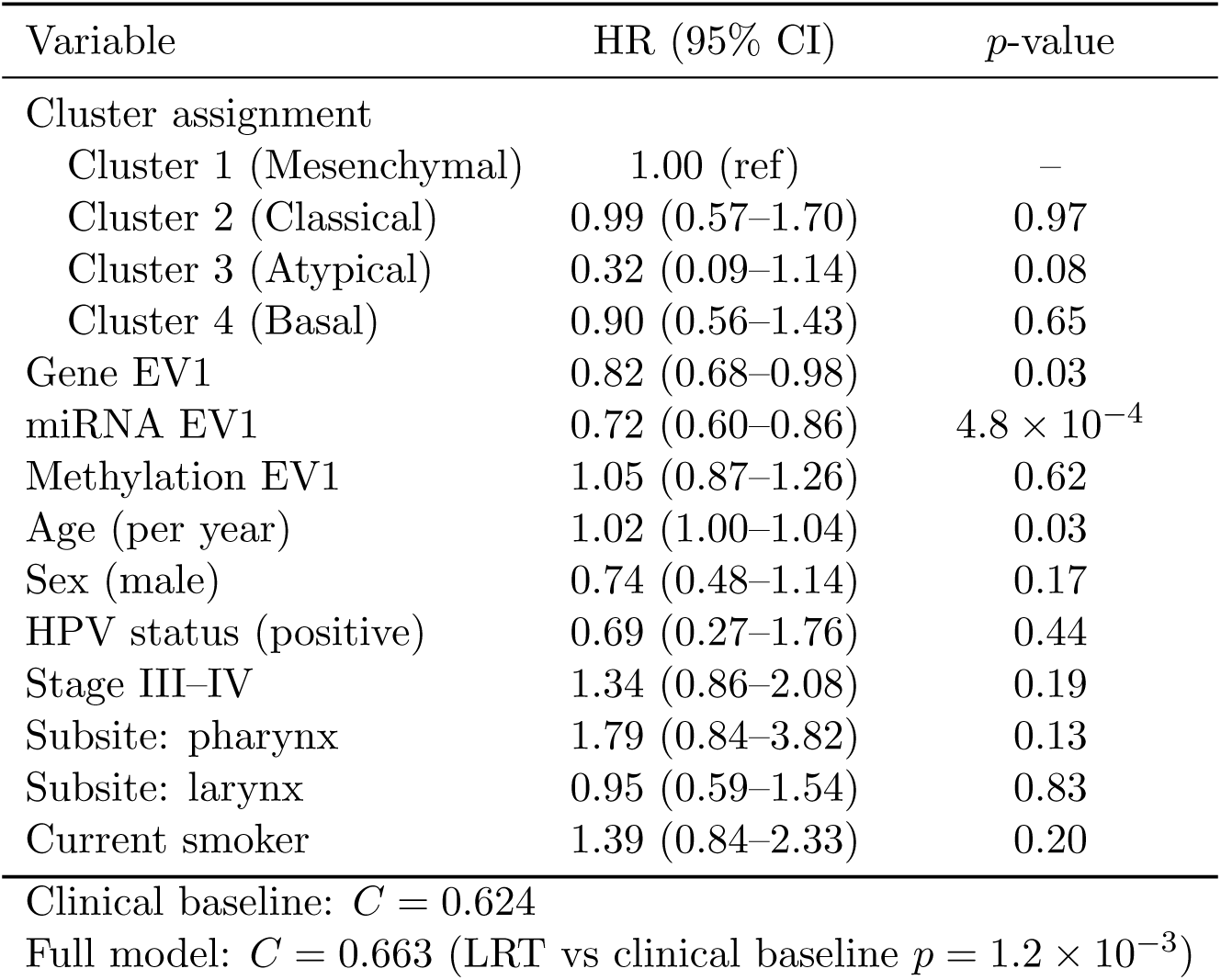
HNSC multivariable Cox model for overall survival. Adjusted hazard ratios (HR) with 95% confidence intervals from the full clinical nested model including cluster assignment, three modality-specific eigenvectors (gene EV1, miRNA EV1, methylation EV1), age, sex, HPV status, stage, anatomic subsite, and smoking status (*n* = 272 complete cases of the 516 patients with overall-survival follow-up; the remainder lacked at least one of HPV status, stage, subsite, or smoking history).

#### 2.3.4 miRNA-Specific Component Encodes an Imprinted and Fetal-Associated Signal

While the gene-specific residual captured the immune microenvironment, the miRNA-specific component showed a different layer of biology. The leading eigenvector loaded negatively on the EMT/stromal program and positively on keratinization, with only a weak interferon-*γ* association (Supplementary Fig. 9). The biology of this axis became clear from its driver miRNAs. More than half of the top-weighted contributors mapped to the chromosome 14q32 DLK1–DIO3 imprinted cluster, with additional loading on the primate-specific C19MC trophoblast cluster, while HPV-responsive and miR-200 reference sets showed no enrichment (Figure 4c, full hypergeometric statistics and target-program assignments in Supplementary Table 8, Supplementary Fig. 9, and Supplementary Tables 9 and 10). Both clusters are normally associated with developmental or placental contexts rather than adult squamous epithelium, so this axis is best read as an imprinted and fetal-associated miRNA signal. The miRNA-specific component is therefore a separate post-transcriptional layer that the shared partition does not distinguish.

This axis also carried modest but detectable survival information. Spectral re-clustering on the miRNA-specific similarity separated EMT-high from keratinization-high tumors more sharply than the shared partition did (Supplementary Fig. 9), and adding the miRNA-specific cluster assignment to a Cox model already containing shared cluster, age, sex, HPV status, stage, subsite, and smoking improved the C-index from *C* = 0.624 to *C* = 0.652 (likelihood-ratio *p* = 4.9 × 10^−3^) (Supplementary Table 11). When the three leading modality-specific eigenvectors were modeled together with shared cluster and the same full clinical covariates, the C-index increased from *C* = 0.624 for the clinical baseline to *C* = 0.663 (*n* = 272 complete cases; Table 4, Supplementary Fig. 10, and Supplementary Table 13; PFI estimates in Supplementary Table 15). In this model, miRNA EV1 remained associated with overall survival (HR = 0.72, *p* = 4.8 × 10^−4^), whereas the methylation-specific eigenvector did not. Unlike gene EV1, the miRNA EV1 association was stable after adding tumor purity, ESTIMATE immune score, or ESTIMATE stromal score to the full clinical model (HR = 0.71–0.72, all *p <* 5×10^−4^; Supplementary Table 14). Together with the gene-specific immune axis, the miRNA-specific component adds another modality-specific layer that carries biological and secondary survival information beyond the discrete shared partition.

Full per-cluster canonical-marker tables, the Keck five-subtype crosswalk, the gene-specific axis with sensitivity-stratified Cox estimates, miRNA-specific program-overlap and putative-target analyses, the methylation-specific component, the multi-method clustering benchmark with parallel progression-free interval analyses, the nested Cox model decomposition, and bootstrap stability of the shared partition are reported in Supplementary Note 5 (HNSC Supplementary Results).

### 2.4 ADNI: Cross-Modal Biological Aging and Dementia Progression Analysis

Predicting MCI/AD conversion in cognitively normal, amyloid- and tau-negative (A−T−) older adults is one of the hardest problems in preclinical AD research [24]. Direct supervised prediction of conversion is difficult even in large cohorts, because conversion from A−T− to MCI/AD is a rare event and classifiers trained on imbalanced, low-event-rate outcomes are prone to overfitting [34]. The problem is compounded with multi-modal data, as the intersection of modalities often reduces the usable sample to a fraction of the full cohort. Because advancing age is the strongest established risk factor for dementia [1], biological age acceleration, the gap between biological age and chronological age, offers a more robust alternative. Rather than learning a conversion classifier directly, age acceleration reframes the question as an unsupervised one. We learn an aging axis from all available subjects regardless of outcome, then test whether individuals who age faster along that axis are also the ones who later progress. Because the aging axis is estimated without using conversion labels, it is less susceptible to overfitting and provides an interpretable, biologically grounded intermediate phenotype. Large-cohort epigenetic studies have associated accelerated biological aging with incident dementia risk [51], and MRI-based brain-age models show that a positive brain-age gap is elevated in MCI and AD [52, 53].

However, existing age-acceleration tools largely fail in the A−T− setting. Structural MRI-derived brain age lags behind molecular changes [23], while traditional epigenetic clocks [18, 15, 26] track chronological aging rather than incipient dementia risk [39]. The common limitation is that these approaches each operate within a single modality, yet the transition from normal aging to neurodegeneration involves coordinated changes across both the epigenome and brain structure [39]. To capture this cross-modal aging signal, we need an integration method that can separate the shared aging component from modality-specific variation. The multiRF shared-specific decomposition addresses this by separating the aging signal coupled across the epigenome and the brain from residual modality-specific variation.

We applied this framework to 87 cognitively normal (A−T−) participants from the Alzheimer’s Disease Neuroimaging Initiative (ADNI) [22], integrating baseline blood DNAm and structural MRI (cohort details, preprocessing, and the evaluation framework are in Methods, Section 4.11). The primary ADNI DNAm analysis used a prespecified 4,332-CpG union panel, formed from 3,503 AD-prioritized CpGs identified in an independent prefrontal-cortex EWAS and 831 CpGs with reported brain–blood methylation correlation, with two overlapping CpGs counted once (Methods, Section 4.11). This targeted panel was intended to enrich the blood methylation layer for AD biology and peripheral-to-brain epigenetic signal. We benchmarked multiRF against single-modality predictors, established integration methods (SNF [43], AJIVE [13], MOFA2 [3]), and four established DNA methylation aging measures (Horvath [18], Hannum [15], PhenoAge [26], DunedinPACE [7]) over a median 4.8-year follow-up (24 conversion events). Our evaluation first assessed the chronological age representation (in-sample *R*^2^) to establish a valid aging axis, then tested whether the resulting biological age acceleration was associated with MCI/AD conversion. The primary associations used Cox models adjusted for age, sex, education, MMSE, and APOE *ε*4, with downstream leave-one-out cross-validation of the age-regression and Cox steps after fixed representation learning. We applied variable selection and pathway enrichment analysis to map these statistical signals back to interpretable AD biology.

#### 2.4.1 Biological Age Estimation

Before interpreting age acceleration, we first verified that each method’s low-dimensional representation contains a clear aging signal. Table 5 reports in-sample age *R*^2^ for all methods. The supervised elastic net brain age model achieved *R*^2^ = 0.706, providing an upper bound for what a model trained directly on chronological age can extract. Among unsupervised multi-modal methods, SNF fused explained the most age variance (*R*^2^ = 0.595), closely followed by multiRF shared (0.567), with MOFA2 shared (0.237) and AJIVE joint (0.188) trailing further behind. For single-modality baselines, MRI PCA alone reached *R*^2^ = 0.570, while DNAm PCA, applied to the same 4,332 CpGs used for multiRF, captured only 0.273. That the multiRF shared signal (0.567) was higher than DNAm PCA (0.273) and close to MRI PCA (0.570) indicates that cross-modal integration recovers age-related variance that blood methylation alone cannot explain, consistent with evidence that aging involves coordinated molecular and structural changes [39]. The lower *R*^2^ of MOFA2 and AJIVE suggests that these factor-based methods capture less of the nonlinear age signal in this small, two-modality setting (Figure 6a).

**Table 5:** ADNI biological age and association with AD conversion. HR is the adjusted hazard ratio per one-year increase in age acceleration for age-in-years biological-age estimates from a Cox model with covariates (age, sex, education, MMSE, APOE *ε*4). DunedinPACE is a pace-of-aging score rather than an age-in-years estimate, so its HR and CV HR are reported per one-SD increase in the pace residual. CV columns report downstream leave-one-out cross-validation of the age-regression and Cox steps after fixed representation learning.

| Category | Method | Age $R^2$ | HR (adj) | $p$ (adj) | $C$ | CV $R^2$ | CV HR | CV $p$ |
| --- | --- | --- | --- | --- | --- | --- | --- | --- |
| multiRF | Shared | 0.567 | 1.194 | 0.010 | 0.653 | 0.501 | 1.169 | 0.011 |
|  | Specific (DNAm) | 0.103 | 1.398 | 0.062 | 0.635 | -0.004 | 1.164 | 0.225 |
|  | Specific (MRI) | 0.010 | 2.085 | 0.628 | 0.570 | -0.030 | 1.165 | 0.806 |
| Multi-modal | SNF fused | 0.595 | 1.128 | 0.083 | 0.615 | 0.551 | 1.114 | 0.093 |
|  | MOFA2 shared | 0.237 | 1.155 | 0.172 | 0.593 | -0.078 | 1.035 | 0.514 |
|  | AJIVE joint | 0.188 | 1.248 | 0.046 | 0.629 | 0.067 | 1.188 | 0.109 |
| Single-modality | MRI PCA | 0.570 | 1.161 | 0.023 | 0.627 | 0.479 | 1.140 | 0.016 |
|  | DNAm PCA | 0.273 | 1.004 | 0.967 | 0.551 | 0.132 | 0.964 | 0.628 |
|  | Concatenated PCA | 0.293 | 1.009 | 0.921 | 0.554 | 0.069 | 0.968 | 0.677 |
| Supervised | Elastic net (MRI) | 0.706 | 1.172 | 0.013 | 0.640 | 0.692 | 1.164 | 0.015 |
|  | Elastic net (DNAm) | 0.980 | 1.048 | 0.814 | 0.548 | 0.979 | 1.048 | 0.814 |
| Epigenetic clocks | Horvath | 0.453 | 0.954 | 0.538 | 0.562 | 0.393 | 0.957 | 0.533 |
|  | Hannum | 0.575 | 1.016 | 0.795 | 0.555 | 0.532 | 1.016 | 0.783 |
|  | PhenoAge | 0.509 | 0.921 | 0.249 | 0.572 | 0.468 | 0.926 | 0.249 |
|  | DunedinPACE | — | 0.796 | 0.339 | 0.559 | — | 0.780 | 0.303 |

#### 2.4.2 AD Progression Analysis

We fit Cox proportional hazards models adjusted for age, sex, education, MMSE, and APOE *ε*4 carrier status (Table 5). multiRF shared had the strongest observed conversion association among all unsupervised methods (HR = 1.194 per year of age acceleration, *p* = 0.010, *C* = 0.653, Figure 6c,d). Each one-year increase in cross-modal age acceleration was associated with an approximately 19% higher hazard of conversion, after adjusting for the prespecified clinical covariates. Kaplan–Meier curves grouped by age-acceleration tertile showed the same pattern. The accelerated tertile (Q3) showed worse conversion-free survival than the decelerated tertile (Q1, Figure 6b). Within the same 87 matched DNAm–MRI participants, the alternative unsupervised integrations did not reach significance after covariate adjustment, and neither did DNAm PCA, concatenated PCA, elastic-net DNAm, or the standard blood-based epigenetic aging measures examined here. MRI-only baselines retained some conversion association, but they remained weaker than the shared cross-modal signal in this matched comparison. Additional direct-coordinate Cox benchmarks, which add each learned representation directly to the clinical Cox model rather than first reducing it to scalar age acceleration, are reported in Supplementary Results and Supplementary Table 18. Compared with the supervised MRI model, multiRF shared achieved a comparable effect size (HR = 1.194 per year) without using conversion labels during representation learning, and with a higher *C*-index (0.653 vs. 0.640). This is consistent with the interpretation that jointly modeling blood DNAm and brain MRI recovers an aging component that single-modality approaches only partly recover. The CpG-panel ablation in Supplementary Results showed that the shared age-conversion signal was directionally consistent when the DNAm feature space was restricted to either component of the biologically motivated union panel (Supplementary Table 19), supporting the use of the union panel as the primary AD- and brain-informed feature space for this preclinical analysis.

The shared-specific decomposition also produced an important result from the DNAm residual. The DNAm-specific component explained very little chronological age variance (*R*^2^ = 0.103), yet it showed a borderline association with AD conversion (HR = 1.398 per year, *p* = 0.062, *C* = 0.635). The difference from DNAm-only approaches is that the multiRF decomposition first removes the shared aging component, isolating epigenetic variation distinct from the main cross-modal aging axis. This residual, not the raw DNAm aging signal, carries the disease-relevant trend. This does not mean that the shared aging axis is unrelated to disease. The shared signal was itself associated with conversion (HR = 1.194 per year, *p* = 0.010), consistent with the interpretation that cross-modal coordinated aging carries AD risk. The raw DNAm data therefore appears to hold at least two survival-related layers: a cross-modal aging component (captured by the shared signal) and a DNAm-specific component (captured by the specific residual). DNAm PCA and elastic net DNAm combine both layers into a single axis dominated by normal-aging CpG changes, diluting the smaller disease-related component below the detection threshold. The multiRF decomposition separates these two layers, allowing each to be tested independently. This explains why three methods applied to the same CpG data produce different results. Additional ADNI analyses, including direct Cox benchmarks, CpG-panel ablation, all-DNAm sensitivity, modality-specific quadrant analysis, per-eigenvector annotations, discrete subtype Cox modeling, and embedding-dimension robustness, are reported in Supplementary Note 6.

## 3 Discussion

We present multiRF, a nonlinear method for cross-modal multi-omics integration that returns a shared representation reflecting variation across data types together with modality-specific residuals that carry data-type-specific signal. Three features set it apart from existing approaches. First, the cross-modal random forest base learner captures nonlinear relationships between data types, whereas AJIVE [13], O2PLS [41], and MOFA2 [3] rely on linear decompositions that can miss complex relationships among molecular layers. Second, the shared–specific separation falls out of a two-step estimation: the cross-omics reconstruction extracts the shared component, and an unsupervised forest on the residual extracts the specific component, with no orthogonality constraints or rank choices. Third, the mixture-model variable selection returns per-feature importance scores for both components, which supports direct biological interpretation.

The HNSC application makes this separation concrete. The shared partition aligned with the four canonical Walter/TCGA expression subtypes without access to any subtype label, and the same partition stayed interpretable under the Keck five-subtype system and the CPTAC-like HPV-negative axes. That breadth of agreement matters: it shows the shared component reflects a main HNSCC disease axis rather than overfitting to one reference classifier. The gene-specific axis then added a second, non-redundant signal, aligning with ESTIMATE immune and stromal scores, tumor purity, and immune cell-type deconvolution, and remaining associated with survival in the full clinical Cox model that contained shared cluster, age, sex, HPV status, stage, anatomic subsite, and smoking status. Methods that output only a single fused graph (SNF [43], intNMF [11]) cannot separate these two signals, because they collapse the etiologic and immune axes into one partition; the linear factor methods (AJIVE [13], MOFA2 [3]) did not recover the four-way subtype structure at comparable resolution in our benchmarks. The miRNA-specific axis put the decomposition to a different use. Its top loadings concentrated in the chromosome 14q32 DLK1–DIO3 imprinted cluster and the primate-specific C19MC trophoblast cluster, pointing to imprinted and fetal-associated miRNA variation distinct from the shared etiologic grouping, and its leading eigenvector also remained associated with overall survival after adjustment for shared cluster and the full clinical covariate set. Read together, the HNSCC results describe a layered disease: the shared component captures the main subtype structure, the gene-specific component adds immune variation beyond it, and the miRNA-specific component adds a post-transcriptional layer enriched for imprinted and fetal-associated miRNA clusters that also tracks overall survival. This picture fits single-cell HNSCC work showing that malignant programs such as partial EMT, epithelial differentiation, stress/hypoxia, and cell-cycle activity coexist with stromal and immune programs in the tumor ecosystem [61]. In the present bulk setting these programs are best read as contextual signatures for interpreting the shared and gene-specific axes, not as direct estimates of single-cell states.

The ADNI application illustrates the same principle in a very different setting. A small, amyloid- and tau-negative cohort (*n* = 87) is ordinarily a hard case for unsupervised integration, and the shared–specific decomposition is central to the analysis. The cross-modal shared signal reflects a coordinated DNAm–MRI aging axis associated with MCI/AD conversion (HR = 1.194 per year, *p* = 0.010, *C* = 0.653) within the matched DNAm–MRI subset used for all multi-modal comparisons. The single-modality DNAm and clock baselines in this matched subset were included for a like-for-like comparison, not to claim that multiRF was benchmarked against every available DNAm-only subject. To address that sample-selection issue, we repeated the DNAm-only baselines in all available cognitively normal participants with DNAm and followup diagnosis; those sensitivity analyses revealed no stronger adjusted conversion associations for DNAm PCA or the established clocks. The DNAm-specific residual in the matched analysis, despite explaining little chronological age variance, showed a weaker exploratory conversion association (*p* = 0.062), which suggests that residual epigenetic variation may hold information the main cross-modal aging axis does not.

We chose the random forest as the default base learner for several practical reasons. Each split reduces variance in the response block, so the resulting sample weights reflect cross-modal dependence rather than within-modality geometry alone. The random subspace mechanism gives built-in variable screening when *p_k_*≫ *n*, and the same multivariate forest structure serves both the supervised cross-modal stage and the unsupervised specific stage by treating the predictor matrix itself as a multivariate pseudo-response and column-resampling at each node [21, 19]. In principle any supervised learner that produces sample-by-sample weight matrices could serve as the base learner, so it can be swapped without changing the rest of the method.

The method has limitations. When a modality-specific subgroup is strongly correlated with the shared phenotype, part of that signal can be absorbed into the shared reconstruction. The sample-by-sample weight matrix requires *O*(*n*^2^) storage, which is manageable for cohorts of several thousand samples but would need sparse versions for much larger datasets. Connection selection is also static: connections are ranked once and not revisited after fusion, so an informative connection that receives a low initial score is permanently excluded.

For the ADNI application specifically, the sample size (*n* = 87) limits statistical power, and the quadrant analysis and the DNAm-specific signal should be read as hypothesis-generating rather than confirmatory; validation in a larger, independent cohort is needed before any clinical conclusion. The primary CpG panel combined two biologically motivated sources, AD-prioritized CpGs for disease relevance and brain–blood correlated CpGs for interpretability of blood DNAm as a proxy for brain methylation, so the union panel is a biologically guided feature space emphasizing both AD biology and peripheral-to-brain epigenetic signal rather than an unbiased genome-wide discovery screen. Repeating DNAm PCA and established DNAm aging measures in all available baseline CN participants with DNAm data (213 subjects, 64 conversion events) again yielded no significant adjusted associations, which eases the concern that the matched-subject restriction alone explains the DNAm-only baseline results (Supplementary Results). The comparison with established DNA methylation aging measures is a focused benchmark against commonly used clocks and pace-of-aging measures, not a definitive evaluation of all available epigenetic age models.

Several directions follow from this work. The shared and specific reconstructions can serve as denoised input features for downstream supervised models and may improve outcome prediction in small-sample settings where raw high-dimensional features overfit. Recent CODEX-based HNSCC work shows that spatial tumor immune architecture, including discrete cellular neighborhoods and tertiary lymphoid structures, can be prognostic [62]; our bulk TCGA analysis does not infer such neighborhoods directly, but the gene-specific immune residual suggests that bulk multi-omics profiles retain information about microenvironment composition, so extending multiRF to infer cellular neighborhoods or spatial ecosystem states from single-cell or spatial multi-omics data is a natural next step. An iterative connection selection that re-evaluates connection quality after fusion could improve stability when some connections carry moderate but useful signal. The decomposition also offers a simple route to cross-modal imputation: the shared reconstruction recovers values predictable from cross-modal neighborhood structure while the specific reconstruction learns within-modality patterns, together borrowing strength across modalities while preserving modality-specific variation. Finally, the base learner need not be a random forest; other tree-based models could offer different trade-offs in flexibility, interpretability, and computational cost.

## 4 Methods

### 4.1 Overview

The multiRF method constructs a sample-level similarity matrix from multi-modal data using multivariate random forests fitted across directed inter-omics connections. A multivariate forest trained to predict one omics layer from another implicitly learns a sample-level weight matrix whose rows encode cross-omics sample neighborhoods. For each block *k*, a per-response weight matrix *W* ^(*k*)^ (the modularity-weighted average of connections where *k* is the response) is a weighted average across biologically similar samples: *X̂*^(^*^k^*^)^ = *W* ^(*k*)^*X*^(*k*)^ is designed to capture the cross-omics consensus, and the residual *R*^(*k*)^ = *X*^(*k*)^ − *X̂*^(^*^k^*^)^ is designed to focus on modality-specific variation. A global weight matrix 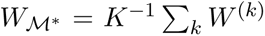 averages the per-response matrices to capture shared structure across all modalities. Shared and specific signals are then clustered independently without explicit latent-factor estimation or rank assumptions. The pipeline proceeds as follows. First, for each directed connection (*a* ← *b*) between omics blocks, a multivariate random forest is trained with block *b* as the predictor and block *a* as the response, yielding a per-connection forest weight matrix *W_m_* (Section 4.3). The response subsampling step (*q*_try_) regularizes the split criterion when the response block is high-dimensional, reducing the influence of high-dimensional noise on the learned weight matrix. Each connection is scored by the modularity of its weight matrix (Section 4.4), and the per-connection weights are fused into per-response matrices *W* ^(*k*)^ and then into a global matrix *W_M∗_* (Section 4.5). Each omics layer is then decomposed into a shared reconstruction *X̂*^(^*^k^*^)^ = *W* ^(*k*)^*X*^(*k*)^ (using the per-response weight matrix) and a residual *R*^(*k*)^ (Section 4.6), and shared and omics-specific similarity matrices derived from the weight matrices are clustered independently (Section 4.7). Full algorithm for the pipeline is provided in Supplementary Note 2.3.

### 4.2 Notation and Setup

Let 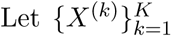 be a multi-modal dataset comprising *K* modalities (for example, gene expression, DNA methylation, miRNA, or imaging) measured on a common set of *n* samples 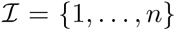. Each matrix 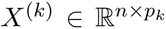 contains *p_k_* features for the same *n* samples. We refer to each matrix as an omics layer, meaning one measured data type. We write 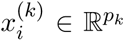 for the *i*-th row and treat each layer as column-standardized prior to forest fitting so that splitting rules are invariant to the feature-wise scale of each omics layer. Sample alignment across modalities is assumed upstream, which matches the typical setting in cancer genomics and imaging-genetics studies where multiple assays are collected on the same biospecimen or subject visit.

We define a set of directed connections

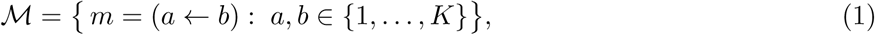

where connection *m* = (*a* ← *b*) specifies block *b* as predictor and block *a* as response. ℳ may include cross-omics (a ≠ *b*) and within-omics (*a* = *b*) connections. Given the multi-modal data and ℳ, our goal is to construct a sample-level similarity matrix *S* ∈ ℝ*^n^*^×^*^n^* that reflects shared predictive neighborhood structure across modalities, suitable for downstream clustering into *G* groups 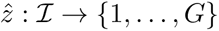.

### 4.3 Directed Cross-Omics Forests

#### 4.3.1 Multivariate splitting rules

To model each cross-modal connection 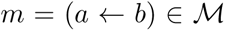, we fit a multivariate random forest with *X*^(*b*)^ as the predictor matrix and *X*^(*a*)^ as the multivariate response. Unlike a univariate regression forest, each tree is grown by a composite splitting rule that considers multiple response dimensions jointly [35, 46]. At each candidate split in node *t*, a random subset 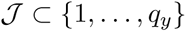 of size *q*_try_ is drawn from the *q_y_* response columns (analogous to the predictor-side mtry but applied to responses), and the selected responses are standardized to unit within-node variance. The split score is

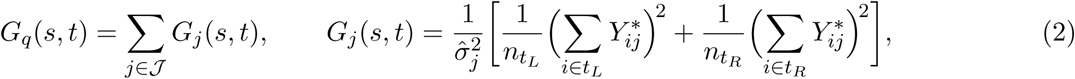

where 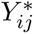 is the centered value of response variable *j* for sample 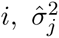 is the within-node variance of column *j*, and *t_L_, t_R_* denote the left and right daughter nodes with sizes 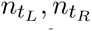. The default *q*_try_ = ⌈*q_y_/*3⌉ balances signal inclusion against noise so that each split evaluates a random one-third of the response columns. Different splits within the same tree, and different trees within the forest, draw different subsets 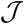, so the averaged weight matrix combines different views of the joint response structure.

When *q_y_*≫ *n*, evaluating all *q_y_* dimensions at every split allows high-dimensional noise to dominate the composite criterion, producing terminal nodes that reflect spurious response variation rather than true cross-modal structure. Restricting the split evaluation to *q*_try_ ≪ *q_y_* random response columns regularizes the criterion, reducing its effective variance while preserving the multivariate character of the tree. Across the forest, different trees evaluate different random subsets, so the averaged weight matrix *W_m_* aggregates information from all response dimensions despite each individual split seeing only a subset. This mechanism reduces the influence of high-dimensional noise on the weight matrix rows. With an appropriate *q*_try_, signal-carrying response dimensions have greater influence on the split criterion, and the resulting weight vectors *w_i_* place more mass on informative neighbors. An empirical study of this choice across signal-sparsity settings is given in Supplementary Note 1.

#### 4.3.2 Forest weights matrix

The forest consists of *B* trees. Each tree *t* partitions the *n* samples into terminal nodes. Let *ℓ_t_*(*i*) denote the terminal node to which sample *i* is assigned in tree *t*. The forest weight matrix for the cross-modal connection *m* is defined as:

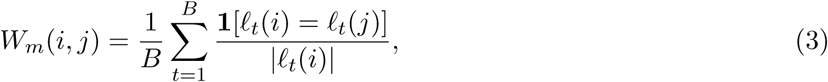

where |*ℓ_t_*(*i*)| is the number of samples in the terminal node containing sample *i* in tree *t*. By construction, *W_m_*(*i, j*) ≥ 0 and Σ*_j_ W_m_*(*i, j*) = 1 for all *i*, so the forest prediction 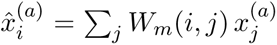 is a weighted average of observed values [28]. *W_m_* is a prediction operator and a data-driven, sample-level similarity measure. Samples that frequently co-occur in small terminal nodes receive large weights, indicating a tight predictive neighborhood along the direction *a* ← *b*.

Because sample *i* always belongs to its own terminal node, the diagonal entry *W_m_*(*i, i*) is strictly positive. If used directly, the reconstruction includes the observed value itself, shrinking the residual toward zero and attenuating omics-specific signal. We remove this self-contribution by setting the diagonal to zero and redistributing the weight proportionally:

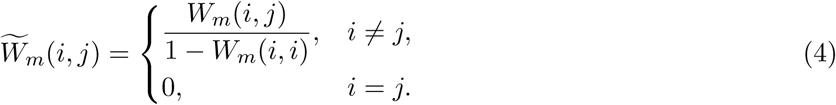

For each row, only the *v* largest entries are retained, the rest set to zero, and the row renormalized to sum to one. This restricts each sample’s neighborhood to its *v* most similar peers, suppressing weak co-occurrences from chance terminal-node overlap. The truncation level *v* is selected by an *entropy-elbow* criterion. We sweep a grid of candidate *v* values and compute the mean row entropy 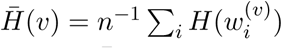, where 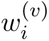 is the *i*-th row truncated to its top-*v* entries and renormalized. As *v* increases, *H̅* (*v*) rises (more neighbors retained) until additional neighbors contribute little entropy. The elbow (the first *v* at which the smoothed marginal gain falls below a fraction of the peak gain) is selected as *v*^∗^. When the forest weights are already highly concentrated and the selected *v*^∗^ ≥ 0.8 *n*, truncation is skipped. For notational convenience, we write *W_m_* for the adjusted and truncated matrix in the following text. A detailed derivation of this adjustment is given in Supplementary Note 2.1.

### 4.4 Connection Scoring

Different directed connections carry different amounts of cross-modal signal. Rather than selecting a subset of connections and discarding the rest, we score every connection by the modularity of its forest weight matrix and use these scores as fusion weights, so that informative connections contribute more to the fused matrix while weak connections are down-weighted automatically. The modularity scores defined below feed directly into the per-response fusion coefficients *α_km_* of Eq. (6) in Section 4.5. The current subsection records how those scores are computed from each connection’s forest weight matrix.

For each connection 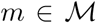, the raw forest weight matrix is symmetrized as 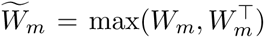 and treated as a weighted adjacency matrix. Weighted Louvain community detection [6] is applied to the resulting graph, and Newman’s modularity [30]

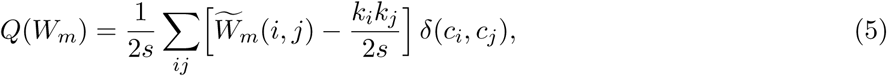

is recorded, where 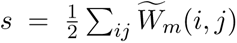 is the total edge weight, 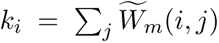 is the strength of node *i*, *c_i_* is the community assignment, and *δ* is the Kronecker delta. High modularity indicates that the neighborhood graph has clear block structure compatible with downstream clustering. Low modularity signals a diffuse graph whose rows are nearly uniform. All connections are retained for fusion and the modularity scores 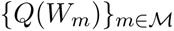 serve solely as quality-based fusion weights (Section 4.5).

### 4.5 Cross-Connection Fusion

Fusion aggregates the connection-specific weight matrices 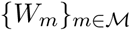 into a single sample-level weight matrix 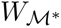 in two stages. An inner per-response stage first combines connections sharing a common response block, and an outer global stage then averages across response blocks. This two-stage design prevents any single modality from dominating the fused similarity simply because it was chosen as the response of many connections, and it allows the modularity-based quality weights computed in Section 4.4 to enter the fusion at the stage where they are most informative (within a response block, where the modularity scores are directly comparable).

For each block *k*, define 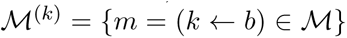 as the subset of connections in which block *k* is the response. Within this subset, each connection is weighted by its modularity:

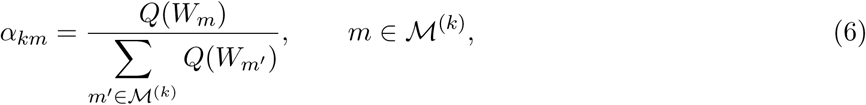

and the per-response weight matrix is

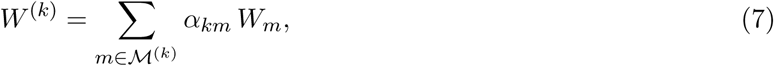

followed by row-renormalization. Because the normalization in Eq. (6) is restricted to 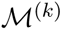, the per-response matrix *W* ^(*k*)^ is built exclusively from forests trained to predict block *k*. Their terminal-node structure is therefore optimized for reconstructing block *k*.

The global weight matrix is the uniform average of the per-response matrices:

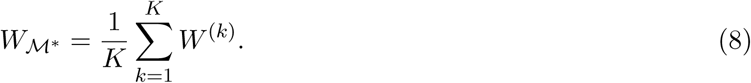

When all connections have comparable modularity and each response block has the same number of predictor connections, this reduces to the uniform average over all *W_m_*. Because the modularity scores act as soft quality weights, weak connections contribute proportionally less without being discarded entirely, preserving cross-modal information from every directed connection. Optional top-*v* truncation can be applied to 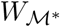 if further sparsification is needed.

### 4.6 Shared-Specific Decomposition

We assume each omics layer decomposes as

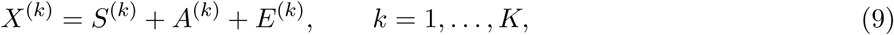

where *S*^(*k*)^ denotes the shared component driven by cross-modal structure common to all modalities, *A*^(*k*)^ is a modality-specific component present only in block *k*, and *E*^(*k*)^ is noise. Concretely, samples sharing the same shared cluster assignment *Z_i_* = *Z_j_* have similar *S*^(*k*)^ profiles, while samples in the same modality-specific subgroup 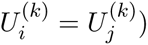 have similar *A*^(*k*)^ profiles. This is a *generative* decomposition. multiRF does not estimate *Z* or *U* ^(*k*)^ as latent factors but instead separates *S*^(*k*)^ from *A*^(*k*)^ directly at the data level through cross-modal smoothing and residual extraction, without explicit dimensionality reduction or factor estimation. AJIVE [13] and O2PLS [41] estimate analogous shared and specific components under explicit rank or variance-decomposition assumptions. SNF [43] and intNMF [11] produce a single fused representation, mixing shared with specific signal.

#### Per-response reconstruction and residual

Using the per-response weight matrix *W* ^(*k*)^ defined in Eq. (7), the marginal reconstruction is

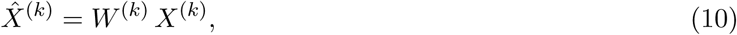

and the residual is

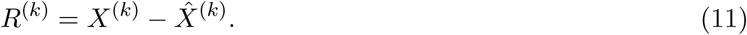

Using the per-response *W* ^(*k*)^ rather than the global 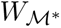 for reconstruction is important for the quality of the decomposition. Because *W* ^(*k*)^ is built exclusively from forests trained to predict block *k* (Section 4.5), the reconstruction is optimized for block *k*. If the global 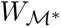 were used instead, connections where block *k* served as the *predictor* (not the response) would contribute, causing the reconstruction to under-fit and shared signal to leak into the residual *R*^(*k*)^. The global 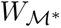 is reserved for the shared-clustering step (Section 4.7), where a single unified similarity across all modalities is needed.

#### Specific weight matrix

To extract remaining structure from the residuals, we fit an unsupervised random forest [21, 19] on each *R*^(*k*)^ independently, yielding a block-specific weight matrix 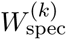 defined analogously to Eq. (3) (full construction via column-wise pseudo-responses in Supplementary Note 2.2). The specific reconstruction

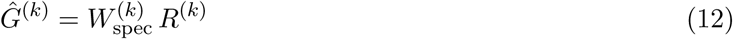

estimates the modality-specific component *A*^(*k*)^ by local averaging in the geometry of the residual. Because 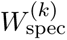 is estimated from *R*^(*k*)^ alone, it is designed to focus on residual structure after the cross-omics reconstruction step. This gives a practical diagnostic for modality-specific signal and leakage without requiring explicit orthogonality constraints [13].

### 4.7 Clustering

From the shared weight matrix 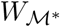 and the omics-specific weight matrices 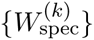 we derive sample-level similarity matrices and cluster them with spectral clustering, producing one shared partition that reflects cross-modal structure and *K* specific partitions that reflect modality-specific structure. Treating the shared and specific components symmetrically, rather than collapsing them into a single affinity, preserves the interpretational distinction between disease-defining axes that align across omics and signals that are only resolvable within a single data type.

#### Shared clustering

The second-order similarity matrix is defined as

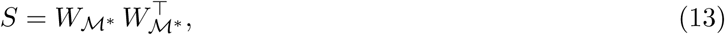

with the diagonal set to zero. *S*(*i, j*) = ⟨*w_i_, w_j_*⟩ measures overlap between the neighborhood weight vectors of samples *i* and *j*: two samples are similar when they draw on the same set of neighbors with comparable weights. Because each row of 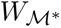 sums to one and is nonnegative, *S*(*i, j*) ∈ [0, 1] and the matrix is symmetric. Spectral clustering is then applied to *S*. The number of clusters is selected by the eigengap criterion.

#### Specific clustering

The same procedure is applied to each omics-specific weight matrix 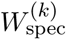, yielding per-block partitions 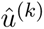. Because the residual signal is typically weaker than the shared signal (the main cross-omics structure has been removed), specific clustering is the harder task.

#### Alternative clustering strategies

The multiRF package also includes proximity-based clustering variants that aggregate terminal-node co-occurrence counts instead of operating on the weight matrix. In simulation studies these variants did not improve upon the second-order similarity approach described above (Supplementary Note 3), and we therefore use the similarity-based strategy throughout the main experiments.

### 4.8 Competing Methods

We compare multiRF against nine established cross-modal integration methods. Unless stated otherwise, the same method set and evaluation protocol are used across all simulation studies in this section. Supplementary Table 1 summarizes all methods evaluated in the experiments. The multiRF method is implemented as a standalone R package (https://github.com/novawz/multiRF) featuring a fast implementation of multivariate random forests [35], the cross-modal variable selection methods introduced in our prior work [46], and the clustering pipeline presented here. The package also supports randomForest-SRC [20, 19] as an alternative back-end. All baseline methods are run with default settings except that dimension-related hyperparameters (number of components or latent rank) are set to *K*−1 to match the rank of the shared signal, giving each method a fair chance to separate all clusters. O2PLS is inherently a two-view method and therefore uses only the first two omics layers. For all methods, the true number of clusters *K* is provided as input to the downstream clustering step. To ensure fair comparison across multiRF variants, the top-*v* tuning performed by the first multiRF variant is cached and reused by all remaining variants on the same dataset.

### 4.9 Simulation Studies

We validate multiRF through two simulation studies. Clustering quality is assessed via the Adjusted Rand Index (ARI*_Z_*) for shared-structure recovery, ARI*_U_* for omics-specific recovery when a method returns block-specific partitions, and a leakage diagnostic that summarizes how cleanly the shared and modality-specific partitions are separated. Here *Z* denotes the true shared label and *U* the modality-specific label. The leakage diagnostic is the mean of ARI*_Z_*_→_*_U_* (ARI of the method’s shared partition against the true modality-specific labels) and ARI*_U_*_→_*_Z_* (ARI of the method’s modality-specific partition against the true shared label); a value near zero indicates a well-separated decomposition where neither partition recovers the other layer’s signal. The number of shared clusters is denoted by *K_Z_*, and the number of modality-specific clusters is denoted by *K_U_* (details in Supplementary Note 4).

#### 4.9.1 Simulation 1: InterSIM Benchmark

Using the InterSIM package [12], we generated correlated three-block profiles (methylation, gene expression, protein) with TCGA-derived covariance structures, injecting both shared (*K_Z_*∈ {4, 8}) and modality-specific (*K_U_* = 2) cluster signals. Here, TCGA-derived covariance structures refer to the empirical within- and between-omics covariance patterns included in InterSIM from TCGA ovarian cancer methylation, gene-expression, and protein data; using these structures makes the simulated features correlated in a way that resembles real multi-omics data rather than independent Gaussian noise. The shared signal was generated by assigning each sample to one of *K_Z_* common clusters and shifting a subset of features in each omics layer according to that common label. To add omics-specific signal, we then assigned each omics layer its own independent two-class label (*K_U_* = 2), independent of the shared label, and shifted a separate subset of features only within that layer according to its layer-specific label. We evaluated 16 factorial scenarios (*n* ∈ {500, 1000}, *δ* ∈ {0.5, 2.0}, *ρ*_noise_ ∈ {0, 0.2}) with 30 replications each. Full data-generating process, equations, and parameter tables are provided in Supplementary Note 4.

#### 4.9.2 Simulation 2: Nonlinear JIVE Benchmark

To test stability under model mismatch, we adopted a multi-view latent factor model inspired by Joint and Individual Variation Explained (JIVE) [29] with increasing nonlinear distortions (*f_J_* ∈ {id, mixed}) applied to the joint component, where id leaves each shared feature unchanged, *x*^2^ squares every shared feature, and the mixed regime assigns each shared feature independently to either id or *x*^2^ with probability 0.5. The identity setting is the case in which the data follow the linear structure assumed by factor models. The mixed setting keeps some shared features linear and squares others, creating a nonlinear shared signal that challenges linear decomposition methods. Details are in Supplementary Note 4.

### 4.10 Real-Data Application: TCGA HNSC

#### 4.10.1 Cohort and Data

We applied multiRF to 517 head and neck squamous cell carcinoma (HNSCC) patients from The Cancer Genome Atlas [50], integrating three omics modalities: gene expression (RNA-seq, 5,000 most variable genes), miRNA expression (miRNA-seq, all 525 miRNAs passing quality filters), and DNA methylation (Illumina 450K array, 5,000 most variable CpG sites), all measured on the same tumor specimens. For gene expression and DNA methylation, feature selection to the top 5,000 per modality was performed by marginal variance ranking after log-transformation and quantile normalization, matching the filtering used in prior TCGA pan-cancer clustering studies [63]. Because the miRNA modality contained fewer than 5,000 features after low-expression and zero-variance filtering, all surviving miRNAs were retained without further selection.

Clinical annotations included HPV status (36 positive, 243 negative among the 279 patients with available status), tobacco smoking history, anatomic subsite (oral cavity, oropharynx, larynx, hypopharynx), AJCC pathologic stage, and overall survival. Somatic mutation calls for nine genes recurrently altered in HNSCC (*TP53*, *CDKN2A*, *CASP8*, *NOTCH1*, *HRAS*, *PIK3CA*, *NFE2L2*, *KEAP1*, *CUL3*) and copy-number burden (fraction genome altered, FGA) were obtained from the same TCGA publication.

All six directed cross-omics connections among the three blocks were specified for multiRF. Gene ↔ methylation, gene ↔ miRNA, and methylation ↔ miRNA (each bidirectional pair expanded into two directed edges, giving six edges in total). This fully connected configuration lets the forest learn predictive neighborhoods in both directions of every pair, consistent with known regulatory links (promoter methylation regulates transcription, miRNAs post-transcriptionally regulate mRNA, methylation shapes and is shaped by miRNA-mediated regulation) and avoiding a priori restriction to a single regulatory chain. Connection quality was assessed by modularity. All six directed connections were retained and fused with modularity-based weights. The number of shared clusters was determined by the eigengap criterion applied to the shared similarity matrix. The eigengap peaked at *k* = 4, consistent with the four canonical expression subtypes identified by Walter et al. [58] and the TCGA [50]: Atypical, Classical, Basal, and Mesenchymal. For omics-specific clustering, spectral eigengap selected *k* = 3 for gene and miRNA and *k* = 4 for methylation.

Of the 517 tumors, 516 had overall-survival follow-up used for the log-rank tests in Figure 3c, 279 had curated centroid-based subtype labels from Walter et al. [58]/TCGA [50] used for agreement metrics, 272 had complete clinical covariates (age, sex, HPV status, stage, subsite, smoking) used for the multivariable Cox model in Table 4, and 243 were HPV-negative and entered the HPV-negative refit (Figure 5). Per-analysis sample sizes are reported in each figure and table caption.

**Figure 5:**
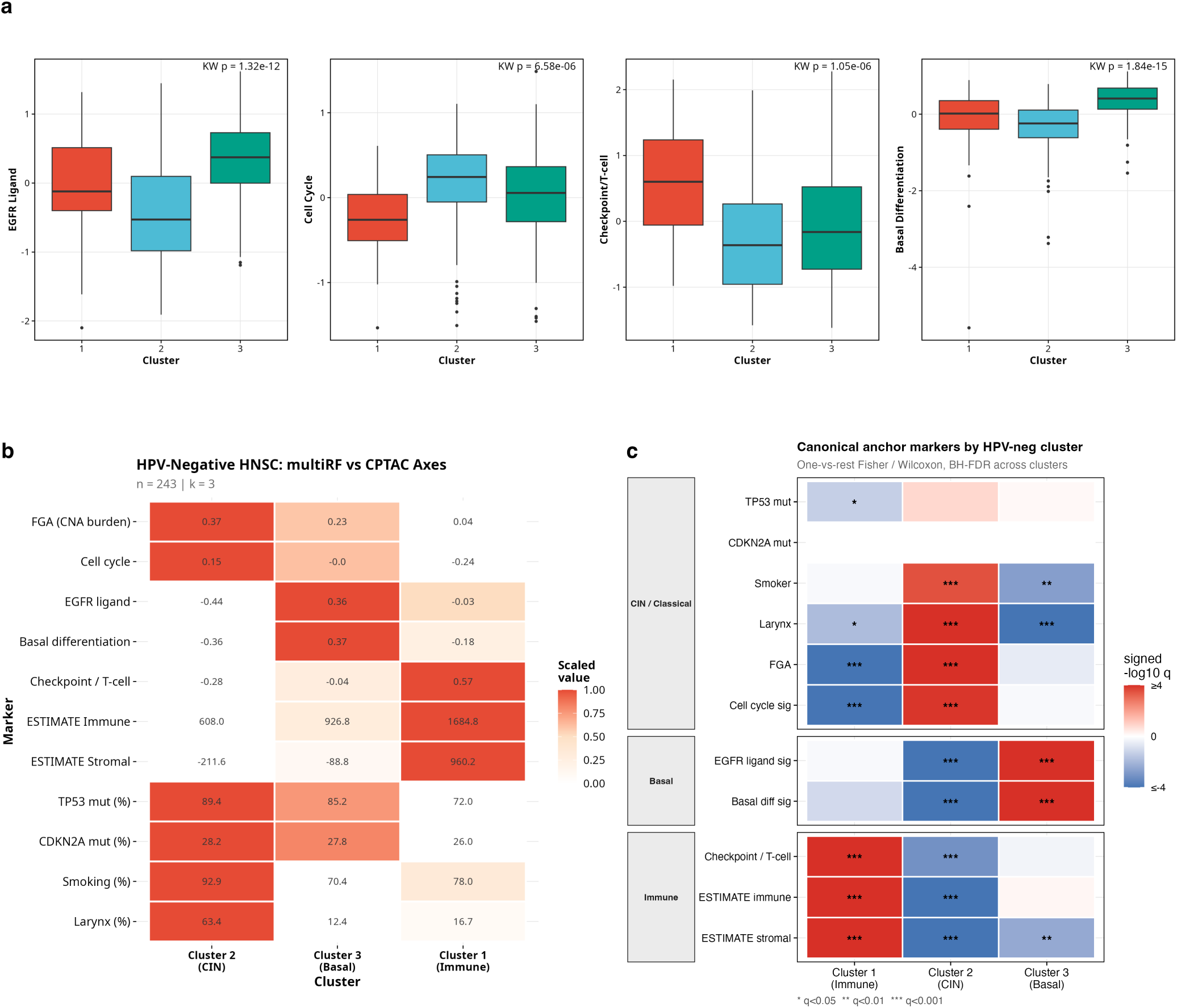
HPV-negative refit of multiRF resembles CPTAC-like CIN / Basal / Immune proteogenomic axes (*n* = 243, *k* = 3). Cluster numbers in this HPV-negative refit are local to the restricted analysis and are independent of the full-cohort cluster numbering. **a** Distribution of four hallmark signatures (EGFR ligand, cell-cycle, checkpoint / T-cell, basal differentiation) across the three HPV-negative clusters, with Kruskal–Wallis *p*-values annotated. **b** Scaled cluster-level profile over CPTAC-like features (FGA, cell cycle, EGFR ligand, basal differentiation, checkpoint, ESTIMATE immune / stromal, TP53 and CDKN2A mutation rates, smoking, larynx), supporting the assignment of Cluster 2 to CIN-like, Cluster 3 to Basal-like, and Cluster 1 to Immune-like. **c** One-vs-rest Fisher / Wilcoxon tests of canonical anchor markers, grouped by expected biology (CIN-like / Classical-like: TP53, CDKN2A, smoking, larynx, FGA, cell cycle; Basal-like: EGFR ligand, basal differentiation; Immune-like: checkpoint / T-cell, ESTIMATE immune, ESTIMATE stromal). Cell shading encodes signed − log_10_ *q*_BH_; asterisks denote *q <* 0.05*/*0.01*/*0.001.

**Figure 6:**
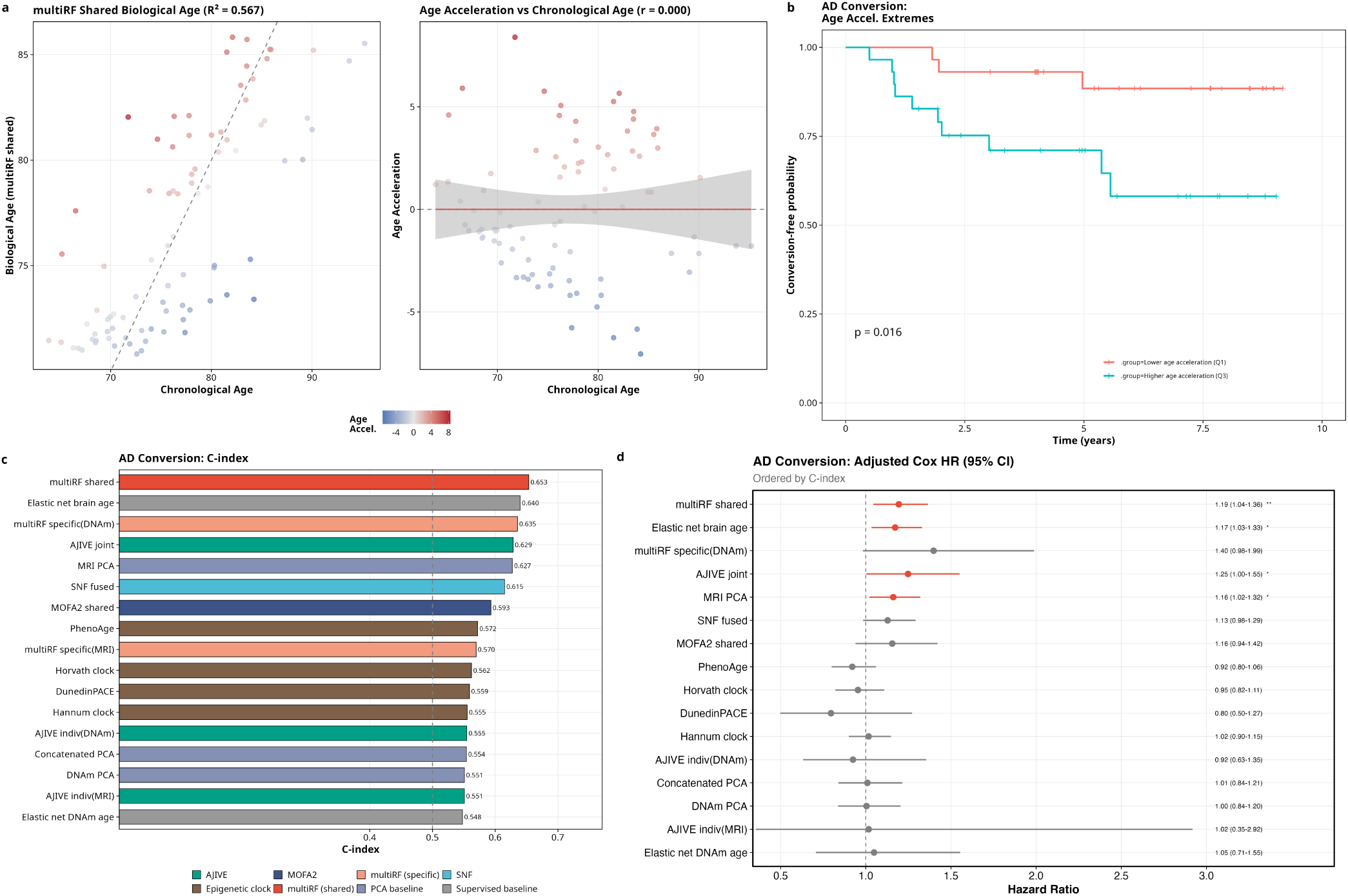
Cross-modal age acceleration and MCI/AD conversion in ADNI (*n* = 87, 24 conversion events). **a** multiRF shared biological age vs. chronological age (left) and the corresponding age-acceleration residual vs. chronological age (right). Age acceleration is defined as the residual from the biological-age fit. **b** Kaplan–Meier curves for AD conversion stratified by multiRF shared age-acceleration tertile (Q1 = decelerated, Q3 = accelerated). **c** *C*-index for AD conversion discrimination across all age-acceleration methods. multiRF shared achieves the highest discrimination among unsupervised methods (*C* = 0.653), higher than SNF fused (*C* = 0.615), established epigenetic clocks, and single-modality baselines. **d** Forest plot of adjusted hazard ratios for AD conversion. HRs are shown per one-year increase in age acceleration for age-in-years estimates and per one-SD increase for DunedinPACE, with the vertical dashed line at HR = 1. The locked primary estimate for multiRF shared is HR = 1.194 per year of age acceleration, *p* = 0.010, and *C* = 0.653.

#### 4.10.2 Subtype Framework Comparison

To assess the biological content of the multiRF shared clusters, we compared them against three established HNSCC subtype systems. First, the Walter/TCGA four expression subtypes [58, 50] were available for 279 patients (those with centroid-based subtype assignments in the original TCGA publication). Agreement was quantified by the Adjusted Rand Index (ARI), Normalized Mutual Information (NMI), Cramér’s *V*, and Fisher’s exact test. Each cluster was assigned its most frequent TCGA subtype label. Second, we mapped each cluster to the five-subtype Keck system [59], which stratifies tumors by consensus gene-expression clustering into two HPV-positive subtypes (HPV-Keratinization and HPV-Immune) and three HPV-negative subtypes (basal, mesenchymal, and classical), with copy-number and mutation profiles annotated for each subtype. Mapping was based on phenotype agreement (HPV status, smoking, anatomic site, mutation profile). Third, for HPV-negative patients only, we compared cluster profiles with CPTAC-like proteogenomic axes described by CPTAC [60]. CIN-like (high chromosomal instability, larynx enrichment, cell-cycle pathway activity), Basal-like (EGFR ligand expression, basal differentiation), and Immune-like (high immune and stromal scores, checkpoint gene expression).

#### 4.10.3 HPV-Negative Analysis

Because HPV-positive and HPV-negative HNSCC are biologically distinct diseases with different driver mutations, prognosis, and treatment response [64], we performed a separate multiRF analysis restricted to HPV-negative patients. The three omics matrices were subset to the HPV-negative samples (*n* = 243), and multiRF was rerun from scratch using the same six-connection structure as the full cohort and *k* = 3 shared clusters to compare with the three CPTAC-like axes. Clustering analysis on the HPV-negative shared similarity matrix was used to assess whether *k* = 3 is supported by the spectral structure of the data. For each of the three resulting clusters, we computed CPTAC-like signature scores. An EGFR-ligand score (mean of scaled *AREG*, *TGFA*, *HBEGF*, *EREG*) [69, 60], a cell-cycle / G1–S pathway score (*CCND1*, *CDK4*, *CDK6*, *RB1*, *CDKN2A*, *E2F1*, *CCNE1*) [70, 60], a checkpoint/T-cell score (*CD8A*, *PDCD1*, *CD274*, *CTLA4*, *LAG3*, *TIGIT*, *GZMB*, *PRF1*, *IFNG*) [71, 72], and a basal/squamous-differentiation score (*KRT5*, *KRT14*, *KRT6A*, *TP63*, *TRIM29*) [47, 58].

#### 4.10.4 Omics-Specific Continuous Analysis

For downstream interpretation of the modality-specific components, we used continuous spectral coordinates derived from each omics-specific similarity matrix. Given a block-specific similarity matrix *S*^(*k*)^, let

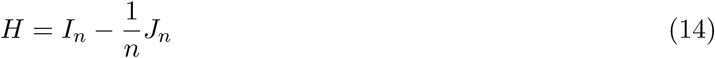

denote the centering matrix, where *I_n_* is the *n* × *n* identity and *J_n_* is the all-ones matrix. We form the double-centered matrix

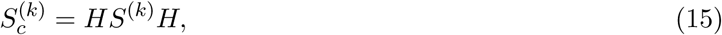

symmetrize it numerically as 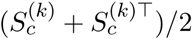, and compute its eigendecomposition

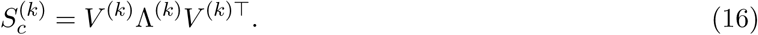

Retaining the positive eigenvalues, we define the spectral coordinate matrix

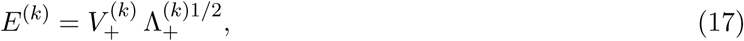

which yields Euclidean coordinates analogous to classical multidimensional scaling. For each omics layer, EV1 denotes the first column of the corresponding matrix E(k). Thus gene EV1, miRNA EV1, and methylation EV1 are the leading continuous coordinates extracted from the gene-, miRNA-, and methylation-specific similarity matrices, respectively. In HNSC, downstream continuous analyses used all three axes, but their biological readouts differed by modality. Gene EV1 was tested against ESTIMATE immune and stromal scores [66], xCell deconvolved cell-type abundance [65], and sensitivity models adjusted for stage, subsite, and purity. miRNA EV1 was interpreted through epithelial and stromal signatures together with enrichment of high-loading miRNAs in curated imprinted and fetal-associated loci. Methylation EV1 was treated as exploratory and related to immune/stromal, keratinization-related, and probe-annotation summaries.

#### 4.10.5 Survival Analysis

Overall survival (OS) and progression-free interval (PFI) were compared across the four shared clusters using the log-rank test, with Kaplan–Meier curves truncated at the clinically meaningful 10-year horizon used by TCGA HNSC analyses. To quantify survival information in the shared and modality-specific signals, we fit nested Cox proportional hazards models. The reduced model included the shared cluster assignment and clinical covariates (age, sex, HPV status, stage, anatomic subsite, and smoking status). The full model added the three leading modality-specific spectral coordinates (gene EV1, miRNA EV1, methylation EV1). Models were fit separately for OS and PFI endpoints. Comparison between the reduced and full models used likelihood-ratio tests and Harrell’s C-index, and coefficient tables report hazard ratios with 95% confidence intervals. This setup tests whether the modality-specific axes contribute independent survival information beyond the shared partition and beyond standard clinical covariates. For the OS model, anatomic subsite was coded as oral, pharyngeal, or laryngeal disease, with hypopharyngeal cases grouped with pharyngeal disease because the complete-case hypopharyngeal stratum was too sparse for a separate Cox coefficient. Tumor purity was evaluated as a sensitivity covariate for the gene-specific immune axis where available.

### 4.11 Real-Data Application: ADNI

#### 4.11.1 Cohort and Models

We applied multiRF to a cohort of 87 cognitively normal (CN), amyloid- and tau-negative (A−T−) subjects from the Alzheimer’s Disease Neuroimaging Initiative (ADNI) [22], integrating two data types (structural MRI and blood DNA methylation, DNAm) measured at the same baseline visit. This early-disease population is the setting in which early biomarkers of neurodegeneration would be most useful.

Blood DNAm was profiled on the Illumina Infinium MethylationEPIC array and preprocessed following the quality-control pipeline described in Zhang et al. [55]. We used a prespecified, biologically motivated panel of 4,332 CpG sites, defined as the union of two external resources. This panel combined (i) 3,503 CpGs identified by an independent prefrontal-cortex epigenome-wide meta-analysis of Braak-stage associations [54], providing AD-prioritized loci, and (ii) 831 brain–blood correlated CpGs from the IMAGE-CpG resource [56], providing cross-tissue evidence that peripheral methylation at these sites may reflect brain-associated epigenetic variation. The two source sets shared two CpGs, yielding a 4,332-CpG union. This targeted CpG selection enriches the DNAm modality for methylation features relevant to AD biology and peripheral-to-brain epigenetic signals, rather than using the full ∼ 850K MethylationEPIC array.

Structural MRI features comprised 85 bilateral cortical and subcortical measures (thickness, volume, and surface area) extracted with FreeSurfer 7.0 from both 1.5 T and 3.0 T acquisitions, normalized by intracranial volume to control for head-size differences. MRI sessions were matched to the DNAm baseline (first visit) time point for each subject. All method comparisons in the main ADNI analysis were restricted to these same 87 matched participants to avoid sample-size confounding between multi-modal and single-modality baselines. As a DNAm-only sensitivity analysis, we also repeated DNAm PCA and the publicly available DNAm clocks in all baseline CN participants with DNAm and follow-up diagnosis (213 subjects, 64 conversions to MCI or dementia). This sensitivity analysis was not used to compare multi-modal methods, because MRI data were not required; it was used only to assess whether DNAm-only baselines changed when the full DNAm cohort was allowed.

#### 4.11.2 Spectral Coordinate Extraction

All methods enter the downstream analyses through a common low-dimensional representation. For mul-tiRF and SNF, we apply the centering, symmetrization, and eigendecomposition pipeline used for HNSC in Section 4.10.4 (Eqs. 14–17) to the shared similarity matrix, yielding shared spectral coordinates *E*^(shared)^. Applying the same pipeline to the multiRF DNAm-specific and MRI-specific residual similarity matrices produces the per-modality coordinates *E*^(DNAm)^ and *E*^(MRI)^. For AJIVE we use the leading principal components of the joint and individual blocks, for MOFA2 we use the leading factor scores, and for single-modality baselines we use the leading principal components of each modality. Throughout, EV*j* denotes the *j*-th column of the corresponding coordinate matrix (equivalently, the *j*-th leading eigenvector of the symmetrized similarity), and EV1–EV5 are used as the common feature space that feeds biological-age regression, discrete clustering, and the per-EV covariate analysis.

#### 4.11.3 Biological Age Estimation

For each method we use the EV coordinates defined in Section 4.11.2 and regress chronological age on the leading five EVs, age ∼ EV_1_ + · · · + EV_5_, and define biological age as the fitted value. Age acceleration is the residual, so that positive values indicate subjects who are biologically older than their chronological age would predict. For established DNAm clocks (Horvath, Hannum, PhenoAge), the clock-predicted age is used directly and age acceleration is the difference from chronological age.

#### 4.11.4 AD Conversion Prediction

The primary endpoint is time from baseline to conversion to mild cognitive impairment (MCI) or AD. We fit Cox proportional hazards models of the form

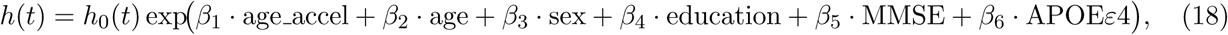

and report the *p*-value for the age-acceleration coefficient, together with the C-index of the full model. In the main method comparison and forest plot, hazard ratios (HRs) are reported per one-year increase for age-in-years acceleration scores. Because DunedinPACE is a pace-of-aging score rather than an age-in-years estimate, its HR is reported per one-SD increase in the pace residual.

#### 4.11.5 Cognitive Association

As a secondary endpoint, we compute the Spearman rank correlation between age acceleration and baseline Mini-Mental State Examination (MMSE) score, under the hypothesis that subjects who appear biologically older than their chronological age should show lower cognitive performance even within this cognitively normal cohort. The rank-based correlation is used rather than Pearson’s to avoid sensitivity to ceiling effects in MMSE, which are common in early-stage samples where most subjects score within one point of 30, and to remain stable under the heavy-tailed distribution of age acceleration produced by the residualization step. A significant negative correlation would indicate that the specific-component axis tracks aging-related cognitive decline before it crosses standard clinical thresholds for mild cognitive impairment.

#### 4.11.6 Discrete Subtype Analysis

Beyond the continuous age-acceleration analyses, we also treat the shared similarity matrix produced by multiRF as the input to a discrete subtype analysis, following the same procedure used for HNSC in Section 4.10.1. Spectral clustering was applied directly to the shared similarity matrix, with the number of clusters selected by the eigengap criterion. The eigengap peaked at *k* = 3, which we took forward for subtype-based survival analysis. For comparison, analogous three-cluster partitions were obtained from SNF, MOFA2, and AJIVE under matched procedures so that all four integrations are evaluated at the same resolution.

For each clustering method we then fit a multivariable Cox proportional hazards model for conversion to MCI or AD, treating cluster membership as the primary exposure and adjusting for the same clinical covariates used in the continuous analysis. Let *z̃_i_* ∈ {1, 2, 3} denote the cluster assignment of subject *i*, with cluster 1 as reference. The cluster-based model is

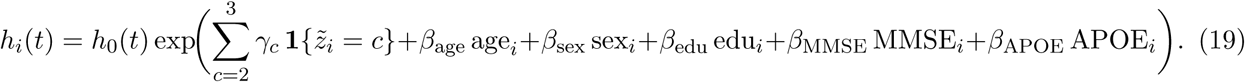

We report adjusted hazard ratios with 95% confidence intervals and Harrell’s *C*-index, and compare the cluster-covariate composition with Kruskal–Wallis tests for continuous variables and chi-squared tests for categorical variables.

#### 4.11.7 Per-EV Covariate Analysis

To understand what each EV represents beyond the main aging axis, we work directly with the shared and modality-specific coordinates {*E*^(shared)^*, E*^(DNAm)^*, E*^(MRI)^} defined in Section 4.11.2. For each of EV1–EV5 from each similarity matrix, we compute associations with clinical covariates (chronological age, sex, education, APOE *ε*4 status, MMSE) and baseline CSF biomarkers (A*β*42, total tau, phospho-tau, restricted to the subset of subjects with lumbar-puncture data). Continuous covariates use Pearson (age, education) or Spearman (MMSE, CSF biomarkers) correlation, binary covariates (sex, APOE4) use Welch’s *t*-test on EV values, and we additionally fit a univariate Cox model of AD conversion on each EV to identify coordinates carrying survival information separable from the main aging axis.

#### 4.11.8 Comparison Methods

We compare 21 methods across five categories.

1. *Single-modality PCA*: MRI PCA, DNAm PCA, and concatenated PCA
2. *Graph-based integration*: SNF (fused and per-modality specific networks).
3. *Multi-modal integration*: multiRF (shared and per-modality specific), AJIVE (joint and individual components), MOFA2 (shared and view-specific factors).
4. *Established DNAm clocks*: Horvath [18], Hannum [15], PhenoAge [26], and DunedinPACE [7].
5. *Supervised baselines*: elastic net brain age (MRI) and elastic net DNAm age (4,332 CpGs and full-array PCs).

For methods that decompose into shared and specific components (multiRF, AJIVE, MOFA2), both representations are evaluated separately, allowing direct comparison of shared versus modality-specific signals for AD prediction. SNF’s “specific” clustering uses the individual pre-fusion affinity matrices, as SNF does not perform an explicit shared–specific decomposition.

## Data Availability

ADNI data are available from the ADNI database (https://adni.loni.usc.edu) upon registration and approval of a data use agreement. TCGA HNSC data are publicly available through the Genomic Data Commons (https://portal.gdc.cancer.gov).

## Code Availability

The multiRF R package is available at https://github.com/novawz/multiRF. Analysis scripts to reproduce all results in this paper are available at https://github.com/TransBioInfoLab/multiRF-cluster.

## Acknowledgments

This research was partially supported by US National Institutes of Health grant R01NS128145.

Data collection and sharing for this project was funded by the Alzheimer’s Disease Neuroimaging Initiative (ADNI) (National Institutes of Health Grant U01 AG024904) and DOD ADNI (Department of Defense award number W81XWH-12-2-0012). ADNI is funded by the National Institute on Aging, the National Institute of Biomedical Imaging and Bioengineering, and through generous contributions from the following: AbbVie, Alzheimer’s Association; Alzheimer’s Drug Discovery Foundation; Araclon Biotech; BioClinica, Inc.; Biogen; Bristol-Myers Squibb Company; CereSpir, Inc.; Cogstate; Eisai Inc.; Elan Pharmaceuticals, Inc.; Eli Lilly and Company; EuroImmun; F. Hoffmann-La Roche Ltd and its affiliated company Genentech, Inc.; Fujirebio; GE Healthcare; IXICO Ltd.; Janssen Alzheimer Immunotherapy Research & Development, LLC.; Johnson & Johnson Pharmaceutical Research & Development LLC.; Lumosity; Lundbeck; Merck & Co., Inc.; Meso Scale Diagnostics, LLC.; NeuroRx Research; Neurotrack Technologies; Novartis Pharmaceuticals Corporation; Pfizer Inc.; Piramal Imaging; Servier; Takeda Pharmaceutical Company; and Transition Therapeutics. The Canadian Institutes of Health Research is providing funds to support ADNI clinical sites in Canada. Private sector contributions are facilitated by the Foundation for the National Institutes of Health (www.fnih.org). The grantee organization is the Northern California Institute for Research and Education, and the study is coordinated by the Alzheimer’s Therapeutic Research Institute at the University of Southern California. ADNI data are disseminated by the Laboratory for Neuro Imaging at the University of Southern California.

## Author Contributions

W.Z. and X.S.C. conceived and designed the study. W.Z. developed the method and R package, performed the analyses, interpreted the results, and drafted the manuscript. X.S.C., L.W. and E.J.F. contributed to interpretation of the results and revision of the manuscript. X.S.C. and L.W. supervised the study. All authors reviewed and approved the manuscript.

