## Supplementary Notes and Figures for "Multivariate Random Forests for Cross-Modal Multi-Omics Integration"

References 17

Supplementary Figures 21

Supplementary Tables 39

### 1 Response Subsampling ($q_{\text{try}}$)

At each tree node the multivariate split criterion is evaluated on a random subset of $q_{\text{try}}$ response coordinates rather than all $q_y$ responses. This is directly analogous to the classical predictor subsampling parameter `mtry` in univariate random forests [9], where using a random subset of predictors per split in regression forests is known to improve generalization by decorrelating trees. We hypothesize that response subsampling plays a similar regularization role on the response side. By forcing each split to rely on a different random subset of response coordinates, the forest avoids overfitting to noise dimensions in high-dimensional response blocks and produces cleaner weight matrices. This subsection uses an empirical sweep to test this hypothesis and to identify a practical default for $q_{\text{try}}/q_y$. The full InterSIM benchmark is shown in Supplementary Fig. 1; this subsection focuses on the response-subsampling sweep in Supplementary Fig. 3.

We generate InterSIM [12] datasets with four shared clusters (proportions $0.1, 0.2, 0.3, 0.4$), fix two directed connections, methyl $\to$ gene ($q_y = 131$) and gene $\to$ protein ($q_y = 160$), and sweep $q_{\text{try}}/q_y$ over 11 values from 0.01 to 1 (where 1 corresponds to using all response columns, i.e. no subsampling). Six scenarios are considered. The first four cross signal strength ($\delta \in \{0.5, 2.0\}$) with sample size ($n \in \{500, 100\}$) and contain no extraneous features. The remaining two scenarios use $n = 500$ and $\delta \in \{0.5, 2.0\}$ but append $\lceil 0.5 \cdot p_k \rceil$ i.i.d. $\mathcal{N}(0, 1)$ columns to every block, inflating the response dimensionality by 50% without adding any cluster-discriminative signal. For instance, the gene block grows from 131 to 197 columns and the protein block from 160 to 240 columns. Each scenario is replicated 10 times with $B = 300$ trees per forest. Three families of metrics are recorded for each replicate. Per-forest clustering quality is measured by the adjusted Rand index (ARI) obtained from spectral clustering on the second-order similarity $WW^\top$. Weight-matrix fidelity is quantified by the Pearson correlation between the vectorised weight matrix at a given $q_{\text{try}}$ and the reference weight matrix obtained with $q_{\text{try}} = q_y$. A fused ARI is also computed after uniform-weight fusion of the two per-connection weight matrices. Wall-clock runtime for forest fitting is also recorded. Supplementary Fig. 3 presents the full results across all six scenarios.

In the four noise-free scenarios, per-forest ARI and weight-matrix correlation both reach a plateau once $q_{\text{try}}/q_y$ exceeds roughly 0.1 to 0.2. Below this range, particularly at $q_{\text{try}}/q_y = 0.01$ (corresponding to only 1 or 2 response columns per split), the criterion evaluates too few columns to reliably detect the multivariate signal, and clustering accuracy drops. Above the plateau, including more response columns per split provides no additional benefit because the signal-carrying columns already appear in each random draw with high probability. The fused ARI confirms that per-forest gains carry over to the integrated result, with the fused ARI near 1.0 for $q_{\text{try}}/q_y \geq 0.1$ in all four noise-free scenarios. Importantly, subsampling at moderate fractions matches the full-response baseline in clustering quality while being much faster, showing that the full response set is unnecessary even when it contains no noise.

The two scenarios with 50% appended noise features reveal the more critical benefit of response subsampling, namely that it acts as an implicit regularizer. When all response columns are used ($q_{\text{try}}/q_y = 1$), every split evaluates the full set of columns including the noise dimensions, which dilute the composite split criterion (Eq. (2)) and produce weight matrices that are more sensitive to high-dimensional noise. Under weak signal ($\delta = 0.5$) with noise, the fused ARI at $q_{\text{try}}/q_y = 1$ falls to the 0.5 to 0.7 range, a substantial degradation compared with near-perfect recovery in the noise-free counterpart. Reducing $q_{\text{try}}/q_y$ to the 0.1 to 0.3 range recovers much of this lost performance because each split then draws only a random subset of response columns, so noise dimensions appear in $\mathcal{J}$ less often and their contribution to the criterion is diluted across many independent draws. In other words, response subsampling does not merely match the full-response model in the presence of noise; it meaningfully outperforms it by preventing noise columns from dominating the split criterion. The per-forest ARI panels show that the noise scenarios have wider interquartile ranges than their noise-free counterparts, reflecting greater variability across replicates, but the median performance at $q_{\text{try}} = \lceil q_y/3 \rceil$ remains close to the noise-free level. Computational time for forest fitting grows roughly linearly with $q_{\text{try}}$, as expected from the per-node cost of evaluating the composite criterion over $q_{\text{try}}$ response columns. At $q_{\text{try}} = \lceil q_y/3 \rceil$ the runtime is roughly $2\times$ to $3\times$ smaller than using all responses, and at $q_{\text{try}}/q_y = 0.1$ the speedup reaches $3\times$ to $5\times$.

These results show that response subsampling is not simply a computational shortcut but a regularization mechanism that actively improves weight-matrix quality when the response block contains uninformative features. The default $q_{\text{try}} = \lceil q_y/3 \rceil$ provides a good balance across all conditions tested. It sits on the performance plateau for clean data, delivers the best or near-best clustering accuracy under noise contamination, and reduces runtime by a factor of two to three. This mirrors the well-established practice of setting $\texttt{mtry} = \lfloor p/3 \rfloor$ for regression forests on the predictor side, where a moderate fraction outperforms both extremes (too few predictors lose signal, all predictors overfit). Just as $\lfloor p/3 \rfloor$ gives a stable default for predictor subsampling without requiring per-dataset tuning, $\lceil q_y/3 \rceil$ serves the same role for response subsampling in multivariate forests.

### 2 Implementation Details

#### 2.1 Adjusted Forest Weight

We provide algebraic detail for the weight adjustment and top-$v$ truncation described in Section 4.3 of the main text. The raw forest prediction for sample $i$ and feature $j$ is $\hat{X}_{ij} = \sum_{\ell=1}^{n} W_m(i,\ell)\, X_{\ell j}$. Since $W_m$ is row-stochastic, this can be written as

$$\hat{X}_{ij} = W_m(i,i)\, X_{ij} + \big(1 - W_m(i,i)\big)\, \bar{X}_{-i,j}, \tag{S1}$$

where $\bar{X}_{-i,j} = \sum_{\ell \neq i} W_m(i,\ell)\, X_{\ell j} / (1 - W_m(i,i))$ is the weighted average excluding sample $i$ (the renormalized weights sum to one). Thus the forest prediction is a convex combination of the observed value $X_{ij}$ and the leave-one-out average $\bar{X}_{-i,j}$. The corresponding residual is

$$R_{ij} = X_{ij} - \hat{X}_{ij} = \big(1 - W_m(i,i)\big)\big(X_{ij} - \bar{X}_{-i,j}\big). \tag{S2}$$

The factor $(1 - W_m(i,i))$ shrinks the residual toward zero, so large self-weights make the reconstruction overly dependent on the observation itself.

The weight adjustment (Eq. (3) in the main text) therefore sets $\widetilde{W}_m(i,i) = 0$ and renormalises the off-diagonal entries by dividing by $1 - W_m(i,i)$. The adjusted prediction becomes the leave-one-out average $\hat{X}_{ij}^{\text{adj}} = \bar{X}_{-i,j}$, and the adjusted residual is

$$\widetilde{R}_{ij} = X_{ij} - \bar{X}_{-i,j}, \tag{S3}$$

which is free of the shrinkage factor. The original and adjusted residuals are related by

$$\widetilde{R}_i^{(k)} = \frac{R_i^{(k)}}{1 - W_m(i,i)}, \qquad i = 1, \dots, n. \tag{S4}$$

The adjustment rescales each row but does not change the identity $X^{(k)} = \widetilde{W}_m X^{(k)} + \widetilde{R}^{(k)}$. The adjusted matrix $\widetilde{W}_m$ retains row-stochasticity and nonnegativity while ensuring that the reconstruction at each sample depends only on other samples.

After top-$v$ truncation, the identity $X^{(k)} = \widetilde{W}_m^{(v)} X^{(k)} + R_v^{(k)}$ continues to hold, with

$$R_v^{(k)} = \widetilde{R}^{(k)} + \big(\widetilde{W}_m - \widetilde{W}_m^{(v)}\big)\, X^{(k)}. \tag{S5}$$

The additional term is the contribution of weak neighbors removed by truncation, so top-$v$ truncation transfers low-weight borrowing from the reconstruction to the residual without altering the basic decomposition identity. The truncated matrix $\widetilde{W}_m^{(v)}$ remains row-stochastic and nonnegative after renormalisation.

#### 2.2 Unsupervised Random Forest via Column-Wise Pseudo-Responses

The unsupervised random forest used here follows the column-resampled splitting rule of Ishwaran [21] and is implemented as the default unsupervised mode of `randomForestSRC` [20, 19]. Unlike the original Shi–Horvath construction [38], which trains a binary classifier to discriminate the real samples from a synthetic column-shuffled copy of the data, the column-resampled rule treats the predictor matrix itself as a multivariate pseudo-response and uses regression-tree splitting throughout. This choice avoids an extra binary classification stage that is incompatible with the multivariate-regression forest weights used in the cross-modal stage (Eq. (3)), and removes the dependence on a single synthetic null draw whose joint structure is destroyed column-by-column and can leak spurious co-occurrences when features are correlated.

Given a residual matrix $R^{(k)} \in \mathbb{R}^{n \times p_k}$ (in our case the block-$k$ residual after cross-modal reconstruction), each tree $b = 1, \dots, B$ in the unsupervised forest is grown recursively as follows. At each node:

1. Sample mtry $= \lfloor \sqrt{p_k} \rfloor$ candidate split variables uniformly without replacement from the $p_k$ columns of $R^{(k)}$.

2. For each candidate split variable $j$, sample ytry pseudo-response columns uniformly without replacement from the remaining $p_k - 1$ columns. We use ytry $= 15$, the `randomForestSRC` default.

3. Standardize the selected pseudo-response columns within the node (subtract the node mean, divide by the node SD) so that the per-column contributions to the split score are on a common scale.

4. Among all admissible split values of variable $j$, choose the value that minimizes the sum of standardized within-child residual sums of squares aggregated across the ytry pseudo-response columns.

5. The pair $(j^*, s^*)$ achieving the lowest aggregated score over all mtry candidate variables is used to split the node.

Splitting continues until each terminal node contains fewer than $2 \times$nodesize samples (we use nodesize $= 3$).

The block-specific weight matrix $W^{(k)}_{\text{spec}}(i, j) = \frac{1}{B} \sum_{b=1}^{B} \mathbf{1}\{L_b(i) = L_b(j)\}/|L_b(i)|$ records, for each tree $b$ and each pair of samples $(i, j)$, the inverse-leaf-size co-occurrence in tree $b$. Because each tree selects its own random subset of pseudo-responses at every node, the forest average over $B$ trees integrates over many views of the pseudo-response geometry, and the resulting $W^{(k)}_{\text{spec}}$ reflects the internal neighborhood structure of $R^{(k)}$ without privileging any single column or linear combination of columns. This is the same forest-weight construction (Eq. (3)) used in the cross-modal stage; the only differences are that there is no external response and that the pseudo-response is resampled per node from the predictor matrix itself.

#### 2.3 Algorithmic Details

**Require:** Multi-omics matrices $\{X^{(k)} \in \mathbb{R}^{n \times p_k}\}_{k=1}^K$; connection set $\mathcal{M} = \{(a \leftarrow b)\}$; number of shared clusters $G$; number of trees $B$

**Ensure:** Shared partition $\hat{z} \in \{1, \ldots, G\}^n$; per-block specific partitions $\{\hat{u}^{(k)} \in \{1, \ldots, G_k\}^n\}_{k=1}^K$

**1. Directed cross-omics forest weight matrices**

**for** each connection $m = (a \leftarrow b) \in \mathcal{M}$ **do**
  Fit multivariate RF with predictor $X^{(b)}$, response $X^{(a)}$, mtry $= \lfloor\sqrt{p_b}\rfloor$, $q_{\text{try}} = \lceil p_a/3 \rceil$, $B$ trees
  $W_m(i,j) \leftarrow \frac{1}{B}\sum_{t=1}^{B} \frac{\mathbf{1}[\ell_t(i)=\ell_t(j)]}{|\ell_t(i)|}$ ▷ *Eq.* (3)
  Diagonal adjustment: $\widetilde{W}_m(i,j) \leftarrow W_m(i,j)/(1 - W_m(i,i))$ for $i \neq j$, 0 for $i = j$ ▷ *Eq.* (4)
**end for**

**2. Connection scoring (quality weights, no filtering)**

**for** each $m \in \mathcal{M}$ **do**
  $Q_m \leftarrow \text{modularity}(\max(\widetilde{W}_m, \widetilde{W}_m^\top))$ ▷ *Eq.* (5)
**end for**
All connections retained: $\mathcal{M}^* \equiv \mathcal{M}$ ▷ *Section 4.4*

**3. Two-stage fusion**

**for** each block $k = 1, \ldots, K$ **do**
  $\mathcal{M}^{(k)} \leftarrow \{m = (k \leftarrow b) \in \mathcal{M}\}$ ▷ *connections with block k as response*
  $\alpha_{km} \leftarrow Q_m / \sum_{m' \in \mathcal{M}^{(k)}} Q_{m'}$ for $m \in \mathcal{M}^{(k)}$ ▷ *Eq.* (6)
  $W^{(k)} \leftarrow \sum_{m \in \mathcal{M}^{(k)}} \alpha_{km} \widetilde{W}_m$; row-renormalise ▷ *Eq.* (7)
**end for**
$W_{\mathcal{M}^*} \leftarrow \frac{1}{K}\sum_{k=1}^K W^{(k)}$ ▷ *Eq.* (8), *uniform global average*

**4. Shared–specific decomposition**

**for** each block $k = 1, \ldots, K$ **do**
  $\hat{X}^{(k)} \leftarrow W^{(k)} X^{(k)}$; $R^{(k)} \leftarrow X^{(k)} - \hat{X}^{(k)}$ ▷ *per-response reconstruction, Eq.* (10)–(11)
  Fit unsupervised RF on $R^{(k)}$ (Algorithm 2) $\rightarrow W_{\text{spec}}^{(k)}$
**end for**

**5. Clustering**

Cluster samples using $W_{\mathcal{M}^*} \rightarrow$ shared partition $\hat{z}$
Cluster samples using each $W_{\text{spec}}^{(k)} \rightarrow$ specific partitions $\hat{u}^{(k)}$

**return** $\hat{z}$, $\{\hat{u}^{(k)}\}_{k=1}^K$

Algorithm 1: multiRF: Multivariate Random Forest Integration and Clustering

**Require:** Residual matrix $R^{(k)} \in \mathbb{R}^{n \times p_k}$; number of trees $B$; candidate-variable subsample size mtry $= \lfloor \sqrt{p_k} \rfloor$; pseudo-response subsample size ytry $= 15$

**Ensure:** Block-specific weight matrix $W_{\text{spec}}^{(k)} \in \mathbb{R}^{n \times n}$

Initialise $W_{\text{spec}}^{(k)} \leftarrow \mathbf{0}_{n \times n}$

**for** $t = 1, \ldots, B$ **do**

Grow a regression tree using $R^{(k)}$ as both predictor and pseudo-response. At each node:

sample mtry candidate split variables from the $p_k$ columns of $R^{(k)}$;

for each candidate $j$, sample ytry pseudo-response columns from the remaining $p_k - 1$ columns of $R^{(k)}$;

standardize the chosen pseudo-response columns within the node;

pick split $s_j^* = \arg\min_s \sum_{c \in \text{ytry}} \left[\text{RSS}_L^{(c)} + \text{RSS}_R^{(c)}\right]$;

split on the $(j^*, s_{j^*}^*)$ pair with the lowest aggregated score over the mtry candidates.

Record terminal-node map $\ell_t(\cdot)$

**for** each pair $(i, j)$ with $\ell_t(i) = \ell_t(j)$ **do**

$W_{\text{spec}}^{(k)}(i, j) \mathrel{+}= 1 / |\ell_t(i)|$

**end for**

**end for**

$W_{\text{spec}}^{(k)} \leftarrow W_{\text{spec}}^{(k)} / B$ ▷ *normalize by number of trees*

**return** $W_{\text{spec}}^{(k)}$

Algorithm 2: Unsupervised Random Forest via Column-Wise Pseudo-Responses

**Require:** Predictor matrix $X^{(b)} \in \mathbb{R}^{n\times p_b}$ (or response $X^{(a)}$); forest of $B$ trees with terminal-node maps $\{\ell_t\}_{t=1}^{B}$; embedding dimension $d_{\text{embed}} = \min(10,\, p,\, n-1)$; sibling weight $\gamma \in [0,1]$
**Ensure:** Enhanced proximity matrix $P^{\text{enh}} \in \mathbb{R}^{n\times n}$

**Step 1: Global PCA embedding**
Compute rank-$d_{\text{embed}}$ PCA of $X$: $E = X\,V_{d_{\text{embed}}} \in \mathbb{R}^{n\times d_{\text{embed}}}$ ▷ *$V_{d_{\text{embed}}}$: top loadings*

**Step 2: Standard and sibling proximity**
Initialise $P^{\text{std}} \leftarrow \mathbf{0}_{n\times n},\ P^{\text{sib}} \leftarrow \mathbf{0}_{n\times n}$
**for** $t = 1,\ldots,B$ **do**
  **for** each terminal node $\ell$ of tree $t$ **do**
    **for** each pair $(i,j)$ with $\ell_t(i) = \ell_t(j) = \ell$ **do**
      $P^{\text{std}}(i,j) \ \mathrel{+}= \ 1/B$ ▷ *standard co-occurrence*
    **end for**
  **end for**

  **for** each internal node with children $\ell_L, \ell_R$ **do**
    $\bar{e}_L \leftarrow \frac{1}{|\ell_L|}\sum_{i\in\ell_L} E_i;\quad \bar{e}_R \leftarrow \frac{1}{|\ell_R|}\sum_{i\in\ell_R} E_i$ ▷ *leaf centroids in PCA space*
    $\rho \leftarrow \text{SpearmanCorr}(\bar{e}_L,\, \bar{e}_R)$
    **if** $\rho > 0$ **then**
      **for** each cross-sibling pair $(i,j)$ with $i\in\ell_L,\ j\in\ell_R$ **do**
        $P^{\text{sib}}(i,j) \ \mathrel{+}= \ \rho\,/\,B$
      **end for**
    **end if**
  **end for**
**end for**

**Step 3: Combine**
$P^{\text{enh}} \leftarrow P^{\text{std}} + \gamma \cdot P^{\text{sib}}$
Symmetrise: $P^{\text{enh}} \leftarrow (P^{\text{enh}} + {P^{\text{enh}}}^{\top})/2$

**return** $P^{\text{enh}}$

Algorithm 3: Enhanced Proximity with PCA-based Sibling Similarity

### 3 Clustering Variant Comparison

The main text presents the second-order similarity approach (Section 4.7) as the default clustering strategy. The MULTIRF package additionally implements two proximity-based alternative strategies, which we describe here and compare empirically.

#### 3.1 Proximity-Based Clustering

Proximity-based variants bypass the weight-matrix reconstruction and aggregate terminal-node co-occurrence across trees and connections into a proximity matrix, an empirical estimate of the probability that two samples share a partition cell [9, 67]. The standard random forest proximity between samples $i$ and $j$ in connection $m$ is

$$P_m(i,j) = \frac{1}{B}\sum_{t=1}^{B} \mathbf{1}[\ell_t(i) = \ell_t(j)], \tag{S6}$$

where $\ell_t(i)$ is the terminal node of sample $i$ in tree $t$. The aggregate proximity across all selected connections is $P = \sum_{m\in\mathcal{M}^*} P_m/|\mathcal{M}^*|$. Spectral clustering is applied to $P$.

#### 3.2 Enhanced Proximity: PCA Embedding and Correlation Details

For tree $t$ let $\ell_t(i)$ denote the leaf of sample $i$, and write $\ell \sim \ell'$ when two leaves share a common parent. The per-tree enhanced proximity is

$$P_t^{\text{enh}}(i,j) = \underbrace{\mathbf{1}[\ell_t(i) = \ell_t(j)]}_{\text{same-leaf}} + \underbrace{\gamma \cdot \max\big(\rho_t(i,j),\, 0\big) \cdot \mathbf{1}[\ell_t(i) \sim \ell_t(j)]}_{\text{sibling-leaf}}, \tag{S7}$$

where $\gamma \in [0,1]$ is a user-specified scaling factor and $\rho_t(i,j)$ is the Spearman rank correlation between the centroids of the two sibling leaves in a common low-dimensional embedding. The forest-level enhanced proximity is $P^{\text{enh}}(i,j) = B^{-1}\sum_{t=1}^{B} P_t^{\text{enh}}(i,j)$, aggregated across connections and fed to spectral clustering exactly as in the standard variant.

A global PCA embedding is computed once from the predictor (or response) matrix with $d_{\text{embed}} = \min(10,\, p_k,\, n-1)$ components, yielding $e_i \in \mathbb{R}^{d_{\text{embed}}}$ and leaf centroid $\bar{e}_\ell = |\ell|^{-1}\sum_{i\in\ell} e_i$. A single global embedding keeps every tree on the same geometric scale; per-leaf or per-split PCA would give incomparable coordinate systems and unstable estimates at typical leaf sizes. Centroid similarity uses Spearman rather than Pearson correlation,

$$\rho_t(i,j) = \rho_{\text{Sp}}\big(\bar{e}_{\ell_t(i)},\, \bar{e}_{\ell_t(j)}\big), \qquad \ell_t(i) \sim \ell_t(j), \tag{S8}$$

because the statistic only needs to detect whether two sibling leaves occupy comparable directions in the embedding, and rank correlation is less sensitive to small leaves and outlying principal coordinates than Pearson. The rectifier $\max(\rho,\, 0)$ suppresses negatively correlated siblings, where a well-separated split is already doing its job, so $\gamma$ controls the maximum strength of cross-sibling borrowing while $\rho$ gates whether borrowing is warranted at each split.

Despite this adaptive smoothing, the proximity matrix is still built from binary tree-membership events and cannot recover the continuous borrowing pattern that the row-stochastic MULTIRF weight matrix provides through the whole set of terminal-node memberships. Enhanced proximity therefore narrows the gap to standard proximity but still falls short of the similarity-based estimator on both clustering accuracy and shared-versus-specific separation, most visibly in scenarios where the two structures partially overlap.

### 4 Simulation Studies

Scenario parameter grids are provided in Supplementary Table 2, and full benchmark results (mean ± SD across 30 replicates) are in Supplementary Table 3.

#### 4.1 Simulation Evaluation Metrics

The following metrics are used in both the InterSIM and NL-JIVE benchmarks. Each method returns a shared partition $\hat{z} : \{1, \ldots, n\} \to \{1, \ldots, K_Z\}$. Methods with an explicit shared and specific decomposition also return block- or view-specific partitions. In InterSIM, the shared label has $K_Z$ classes and each of the three omics layers (indexed by $d$) has an independent two-class specific label $U^{(d)}$ ($K_U = 2$). In NL-JIVE, the shared label has $K_Z = 3$ classes and each of the three views (indexed by $k$) has an independent $K_U = 2$-class specific label $U^{(k)}$. We report recovery of the shared structure, recovery of the block/view-specific structure, and leakage between the two.

**Adjusted Rand Index.** The main metric is the Adjusted Rand Index [68]. For two partitions with contingency table counts $n_{ij}$, row totals $a_i$, column totals $b_j$, and total sample size $n$, the ARI is

$$\mathrm{ARI} = \frac{\sum_{ij} \binom{n_{ij}}{2} - \frac{\sum_i \binom{a_i}{2} \sum_j \binom{b_j}{2}}{\binom{n}{2}}}{\frac{1}{2}\left[\sum_i \binom{a_i}{2} + \sum_j \binom{b_j}{2}\right] - \frac{\sum_i \binom{a_i}{2} \sum_j \binom{b_j}{2}}{\binom{n}{2}}}. \tag{S9}$$

ARI corrects the Rand index for chance agreement, with values near 1 indicating close agreement and values near 0 indicating chance-level agreement.

**Shared-Structure Recovery.** We compare the shared partition $\hat{z}$ against the ground-truth shared label $Z$ using $\mathrm{ARI}_Z$. High values indicate successful recovery of the cross-omics consensus.

**Specific-Structure Recovery.** For methods that produce specific partitions, we compare each $\hat{u}^{(d)}$ (InterSIM) or $\hat{u}^{(k)}$ (NL-JIVE) against the corresponding ground-truth specific label and average across blocks or views. The resulting metric is denoted $\mathrm{ARI}_U$. Methods that do not output specific partitions are marked not applicable for this metric.

**Cross-Evaluation (Leakage).** To assess whether shared and specific signals are cleanly separated, we compute two cross-evaluation quantities and report their mean as a single leakage score. $\mathrm{ARI}_{Z \to U}$ is the ARI between the method's shared partition $\hat{z}$ and each true specific label ($U^{(d)}$ or $U^{(k)}$), averaged over blocks or views. $\mathrm{ARI}_{U \to Z}$ is the ARI between the method's specific partition ($\hat{u}^{(d)}$ or $\hat{u}^{(k)}$) and the true shared label $Z$, averaged over blocks or views. The reported leakage measure is

$$\mathrm{ARI}_{\mathrm{leak}} = \tfrac{1}{2}\big(\mathrm{ARI}_{Z \to U} + \mathrm{ARI}_{U \to Z}\big).$$

Both components should be near zero in a well-separated decomposition: low $\mathrm{ARI}_{Z \to U}$ confirms that the shared partition does not track block- or view-specific structure, and low $\mathrm{ARI}_{U \to Z}$ confirms that the specific partition does not leak shared signal. Their mean therefore provides a single direction-symmetric summary, with values near zero indicating clean separation and values approaching $\mathrm{ARI}_Z$ or $\mathrm{ARI}_U$ indicating substantial leakage.

#### 4.2 InterSIM Simulation: Full Details

The InterSIM benchmark uses a full factorial design with $K_Z \in \{4, 8\}$ shared clusters, $n \in \{500, 1000\}$ samples, $\delta \in \{0.5, 2.0\}$ shared signal strength, and $\rho_{\mathrm{noise}} \in \{0, 0.2\}$ appended-noise fraction. This gives 16 scenarios. Each scenario is replicated 30 times. Full design parameters and shared cluster proportions for each $K_Z$ are listed in Supplementary Table 2. Each modality also has an independent two-class specific label ($K_U = 2$).

##### 4.2.1 Data Generating Process

We use the InterSIM R package [12] to generate correlated multi-modal profiles (DNA methylation, gene expression, protein) from TCGA-derived covariance structures. The simulation proceeds in two stages.

InterSIM [12] generates $n$ samples across $D = 3$ omics layers, methylation ($p_1 = 367$), gene expression ($p_2 = 131$), and protein ($p_3 = 160$), using TCGA ovarian cancer covariance structures. Each sample receives a shared cluster label $Z_i \sim \text{Categorical}(\boldsymbol{\pi})$, and InterSIM introduces differential signal of strength $\delta$ on a subset $\mathcal{S}^{(d)} \subset \{1, \ldots, p_d\}$ of size $|\mathcal{S}^{(d)}| = \lfloor 0.2 \cdot p_d \rfloor$ features per block. The remaining features retain inter-omics correlations but carry no cluster-discriminative signal.

For each block $d$, an independent specific label $U_i^{(d)} \sim \text{Uniform}\{1, 2\}$ is drawn (independently of $Z$), and a random feature subset $\mathcal{T}^{(d)}$ of size $\lfloor 0.2 \cdot p_d \rfloor$ is selected. For each $j \in \mathcal{T}^{(d)}$, a perturbation $\tilde{\delta}_j^{(d)} \sim \mathcal{N}(0, \delta_{\text{spec}}^{(d)})$ is added (in logit space for methylation, in $z$-score space for expression and protein) to samples with $U_i^{(d)} = 2$, then back-transformed to the original scale. Optionally, pure-noise features drawn from $\mathcal{N}(0, 1)$ are appended at fraction $\rho_{\text{noise}}$, and each block retains at most 400 features. In the noisy settings, the number of appended columns is $\lceil 0.2 \cdot p_d \rceil$ for each block. The benchmark therefore tests three things at once, namely recovery of shared clusters under realistic cross-omics covariance, recovery of modality-specific clusters after removing the shared signal, and stability under irrelevant features.

##### 4.2.2 InterSIM Summary Results

Full benchmark results are reported in Supplementary Table 3 and Supplementary Fig. 1. The InterSIM study shows that shared clustering is not the main point of separation between methods, because several approaches recover the shared labels almost perfectly once signal is moderate to strong. Across the full grid, SNF attains the highest mean $\text{ARI}_Z$ (0.999), with multiRF close behind at 0.989. What matters more is that multiRF remains stable in the harder settings. Under weak signal without appended noise ($\delta = 0.5$, $\rho_{\text{noise}} = 0$), its mean $\text{ARI}_Z$ is still 0.985, and with appended noise it remains 0.971. The corresponding values are lower for several comparison methods, including MOFA2 (0.870 and 0.855), RGCCA (0.589 and 0.596), and AJIVE (0.206 and 0.151).

The sharper contrast appears after the shared component is removed, which is the harder part of the benchmark. Here the clearest and most consistent recovery of the modality-specific labels is seen for multiRF. Its mean $\text{ARI}_U$ is 0.785, well above enhanced proximity with soft weighting (0.344), standard proximity (0.290), and AJIVE (0.200), while methods such as SNF, RGCCA, O2PLS, and sPLS are near zero on average. In practical terms, many methods can recover the main shared grouping, but only multiRF continues to separate the residual modality-specific structure once that main signal has been taken out.

The leakage diagnostics support the same interpretation. For multiRF, both $\text{ARI}_{Z \to U}$ and $\text{ARI}_{U \to Z}$ remain essentially zero on average ($-0.002$ and $-0.002$), which indicates that the shared and specific partitions are not simply tracking each other. The proximity-based variants show the same qualitative pattern, whereas several baselines show appreciable carry-over from one layer to the other, especially RGCCA (mean $\text{ARI}_{U \to Z} = 0.272$), SNF (0.232), mixKernel (0.225), and sPLS (0.168). This suggests that their apparent structure is less cleanly separated into shared and specific components. Within the proximity family, the enhanced variant is consistently better than standard proximity, but the gap to the similarity-based multiRF fit remains large in both shared and specific recovery.

The runtime cost is moderate rather than minimal. multiRF requires 20.7 s per dataset on average, which is close to MOFA2 (21.6 s), lower than SNF (51.1 s) and intNMF (86.1 s), and still compatible with repeated simulation evaluation. The proximity variants are not uniformly faster once the full clustering workflow is included, so the second-order similarity formulation remains the default throughout the paper.

##### 4.3 Nonlinear JIVE Simulation: Full Details

The nonlinear JIVE benchmark uses a factorial design with joint nonlinearity $f_J \in \{\mathrm{id}, \mathrm{mixed}\}$, $n \in \{500, 1000\}$, and $p_k \in \{(200, 200, 100), (500, 500, 200)\}$. This gives 8 scenarios, each replicated 30 times. The shared structure has $K_Z = 3$ classes and latent rank $r_J = 2$. Each view has an independent two-class specific structure with rank $r_A = 1$. Joint and individual signal strengths are both set to 2, and the additive noise SD is 0.2.

###### 4.3.1 Data Generating Process

We adopt a multi-view latent factor model inspired by Joint and Individual Variation Explained (JIVE) [29], augmented with element-wise nonlinear transformations. For $K$ views, the observed data for view $k$ is

$$X^{(k)} = \delta_J \cdot f_J\big(\widetilde{\Theta}_J \widetilde{B}_{J,k}^{\top}\big) + \delta_A \cdot f_A\big(\widetilde{\Theta}_{A,k} \widetilde{B}_{A,k}^{\top}\big) + E^{(k)}, \tag{S10}$$

where $\widetilde{(\cdot)}$ denotes column normalization, $f_J, f_A$ are element-wise nonlinearities, $\delta_J, \delta_A$ control the joint and individual signal strengths, and $E^{(k)} \sim \mathcal{N}(0, \sigma_\varepsilon^2)$ i.i.d.

The shared latent scores $\Theta_J \in \mathbb{R}^{n \times r_J}$ encode a $K_Z$-cluster structure in which each sample is assigned a label $Z_i \in \{1, \ldots, K_Z\}$ uniformly at random and $\Theta_{J,i} \sim \mathcal{N}(\delta\, \mu_{Z_i},\, I_{r_J})$, where $\{\mu_c\}$ are regular simplex vertices. Loading matrices $B_{J,k} \in \mathbb{R}^{p_k \times r_J}$ have i.i.d. $\mathcal{N}(0, 1)$ entries. For each view $k$, an independent specific label $U_i^{(k)} \in \{1, \ldots, K_U\}$ generates individual scores analogously, with loadings $B_{A,k}$ orthogonalized against $B_{J,k}$ via Gram–Schmidt.

The observed matrices are built by first projecting the latent scores through the loading matrices, scaling each feature to unit variance, then applying the chosen element-wise nonlinearity. The identity regime leaves every shared feature unchanged after projection and scaling. In the mixed regime, each feature is independently assigned either the identity map or the squaring map with probability 0.5. This preserves the latent cluster geometry but breaks the global linear structure assumed by factor models. Throughout, the individual component remains linear, so the mismatch is isolated to the shared signal In the benchmark reported here, $f_J \in \{\mathrm{id}, \mathrm{mixed}\}$, with $f_A = \mathrm{id}$ throughout. The identity case is the linear oracle, because the shared component remains exactly linear after the latent projection step. The mixed case is the hard setting, because some shared features remain linear while others are squared, so the same shared clusters are still present but no longer represented by a single coherent linear factor model.

###### 4.3.2 Design Rationale

The two simulations test different failure modes. Simulation 1 evaluates methods under realistic omics covariance structures where the shared signal is confounded by modality-specific substructure. Simulation 2 isolates model mismatch due to nonlinearity. When $f_J = \mathrm{id}$, linear factor methods are close to their ideal setting. When $f_J$ is mixed, the joint signal is still present but no longer well described by a single low-rank linear factorization. This is where tree-based neighborhood estimation should help.

###### 4.3.3 NL-JIVE Summary Results

Full benchmark results are reported in Supplementary Table 3 and Supplementary Fig. 2. The NL-JIVE benchmark is more discriminating because it separates the easy linear case from the misspecified nonlinear case. When the joint signal is linear, several methods perform similarly well on the shared partition, which is the expected result under a correctly specified latent factor model. Mean $\mathrm{ARI}_Z$ is 0.876 for SNF, 0.876 for O2PLS, 0.870 for MOFA2, 0.869 for RGCCA, and 0.869 for multiRF. In other words, there is little to separate these methods when the data-generating mechanism is close to the assumptions built into the linear baselines.

The picture changes once the joint component is made nonlinear. SNF remains strong at $\text{ARI}_Z = 0.872$, and multiRF decreases only slightly from 0.869 to 0.855. By contrast, the linear factor methods deteriorate sharply, with O2PLS falling to 0.208, RGCCA to 0.248, MOFA2 to 0.421, and intNMF to 0.019. The main message of this experiment is therefore not that multiRF dominates the easy case, but that it retains most of its performance when the shared structure is no longer well described by a linear low-rank model. The same comparison also shows that enhanced proximity improves on standard proximity but does not close the gap to the similarity-based formulation. For shared clustering, its mean $\text{ARI}_Z$ is 0.532 in the identity setting and 0.491 in the mixed setting, compared with 0.500 and 0.466 for standard proximity.

The same pattern is visible in block-specific recovery. multiRF attains mean $\text{ARI}_U = 0.744$ overall, with very similar values in the identity (0.732) and mixed (0.755) settings. SNF and sPLS have higher raw $\text{ARI}_U$ values, but they are not separating shared and specific structure in the same way as multiRF. Among methods that do attempt an explicit decomposition, multiRF is the most stable across the nonlinearity shift. The enhanced proximity variant again improves on standard proximity, reaching mean $\text{ARI}_U$ values of 0.169 in the identity setting and 0.223 in the mixed setting, versus 0.108 and 0.125, but both remain far below the main multiRF estimator.

The heterogeneous NL-JIVE setting tests a different stress case: the view-specific signal strengths are no longer equal across views ($\delta_A = 4, 1, 0.5$). In this setting, multiRF again retained both shared and specific recovery (mean $\text{ARI}_Z = 0.958$, mean $\text{ARI}_U = 0.837$). SNF remained strong for the shared partition (mean $\text{ARI}_Z = 0.961$), but its raw specific recovery dropped to mean $\text{ARI}_U = 0.325$ and its cross-evaluation leakage increased, consistent with the absence of an explicit residual decomposition. MOFA2 retained moderate shared and specific recovery (mean $\text{ARI}_Z = 0.734$, mean $\text{ARI}_U = 0.653$), whereas AJIVE and RGCCA were more sensitive to the heterogeneous view-specific signal strengths.

Again, the leakage metrics support the substantive interpretation. For multiRF, mean $\text{ARI}_{Z\to U}$ is 0.000 and mean $\text{ARI}_{U\to Z}$ is −0.001, both effectively zero. Several baselines show markedly larger leakage, especially mixKernel ($\text{ARI}_{U\to Z} = 0.437$), RGCCA (0.241), and AJIVE (0.077). As in InterSIM, this indicates that multiRF is not only recovering clusters, but is also keeping the shared and specific layers distinct. This stability comes at a moderate computational cost rather than an extreme one. Mean wall-clock time for multiRF is 63.1 s per dataset, much lower than MOFA2 (153.8 s), and somewhat higher than SNF (43.1 s). Given the gain in stability under nonlinearity, this runtime remains reasonable for the benchmark sizes considered here.

### 5 HNSC Supplementary Results

#### 5.1 Concordance with Established HNSCC Subtype Frameworks

The per-system alignment statistics underlying the main-text cluster-to-subtype mapping are tabulated in Supplementary Table 4, and the full canonical-marker panel across the four shared clusters, together with the accompanying statistical tests, is given in Supplementary Fig. 4 and Supplementary Table 5. Two quantitative details are worth singling out from those materials. First, 96% of labeled TCGA Atypical tumors land in the multiRF Atypical cluster. Second, the Mesenchymal cluster, which is the most heterogeneous in expression-subtype labels because the mesenchymal program overlaps with stromal and immune infiltration, is nevertheless distinguished by elevated *NOTCH1* mutation frequency and the highest ESTIMATE immune score. Within the Keck five-subtype taxonomy, the HPV-driven multiRF cluster splits across both the HPV-Immune and HPV-Keratinization Keck groups rather than mapping to a single Keck label, which reflects the known internal variation of HPV-positive HNSCC (Supplementary Fig. 5 and Supplementary Table 6).

##### 5.2 Gene-Specific Immune Signal

Beyond the ESTIMATE correlations and discrete immune-low/-intermediate/-high clustering reported in the main text, the cell-type deconvolution underlying the gene-specific axis shows that the signal is adaptive and myeloid in character rather than generic, with correlations strongest for dendritic cells, B cells, macrophages, and $CD8^+$ T cells, while NK cells show essentially no association (Supplementary Fig. 7). This distribution argues against the reading that the residual is absorbing noise left over from the shared decomposition, and stratified Cox estimates by gene EV1 tertile within each shared cluster (Supplementary Table 7) quantify the survival contribution of this axis beyond the discrete partition. As a light single-cell-informed sensitivity analysis, we also scored bulk signatures related to HNSCC programs highlighted by Puram et al. [61] (Supplementary Fig. 8). These included immune/T-cell, interferon-$\gamma$, antigen-presentation, EMT/stromal, keratinization or basal differentiation, hypoxia, cell-cycle, and oxidative-stress programs. Gene EV1 was most strongly associated with immune/TME scores (Spearman $\rho = 0.58$ for T-cell, 0.49 for interferon-$\gamma$, 0.42 for antigen presentation, and 0.72 for ESTIMATE immune score in the full cohort) and with a more moderate EMT/stromal signal ($\rho = 0.37$). In contrast, keratinization, basal differentiation, and cell-cycle scores showed little association with gene EV1 ($|\rho| \leq 0.05$). The same pattern was preserved in HPV-negative tumors, supporting the interpretation that the gene-specific residual is primarily an immune/TME axis rather than a broad malignant epithelial-state axis. Because gene EV1 is strongly correlated with immune and stromal infiltration, we also fit Cox sensitivity models that added tumor purity, ESTIMATE immune score, or ESTIMATE stromal score to the full HNSC clinical model. These models were repeated for gene EV1, miRNA EV1, and methylation EV1 and are reported in Supplementary Table 14. The miRNA-specific association remained stable across these adjustments, whereas the gene-specific association was attenuated after explicit adjustment for microenvironmental content.

##### 5.3 HPV-Negative Analysis

Full-cohort spectral $k$-selection diagnostics for the shared and component-specific similarity matrices are summarized in Supplementary Fig. 11. The HPV-negative analysis reported in the main text (Figure 5) serves a narrower interpretive purpose. It shows that when the HPV-driven subgroup is excluded, the remaining HPV-negative tumors do not collapse into a single continuum but resolve the chromosomal-instability, basal, and immune-rich CPTAC-like proteogenomic axes, which supports the reading that the full-cohort four-cluster solution is not simply a HPV-positive versus HPV-negative split with three incidental strata attached.

##### 5.4 miRNA-Specific Biology

Beyond the imprinted-locus accounting and the keratinization association reported in the main text, the miRNA-specific axis sits in a different biological register from the gene-specific immune axis, reflecting a post-transcriptional program tied to epithelial state and viral context that is not explained by the shared subtype partition (Supplementary Fig. 9; program overlap in Supplementary Table 8); putative target-program assignments for the top miRNA-specific drivers are reported in Supplementary Tables 9 and 10, and the miRNA-specific partition Cox summary backing its secondary survival association is given in Supplementary Table 11.

##### 5.5 Methylation-Specific Component

The methylation-specific structure is weaker than either the gene- or the miRNA-specific axis and should be interpreted cautiously, since its clusters only partially recapitulate the TCGA methylation subtypes and do not support a sharp standalone taxonomy, but the methylation-specific eigenvector is associated most strongly with immune score, EMT/stromal variation, and to a lesser extent keratinization, which suggests an additional composition-sensitive epigenetic axis (Supplementary Fig. 12); because the reduced methylation feature set used for clustering does not retain annotated *CDKN2A* promoter probes, we treat this component as hypothesis-generating rather than definitive.

##### 5.6 Prognostic Value and Stability

Stability of the four shared clusters was assessed by 100 bootstrap resamples drawn with replacement at the sample level, preserving the joint multi-omics structure within each subject. For each resample, multiRF was refit end-to-end using identical hyperparameters to the point estimate, and the resulting four-cluster solution was matched back to the original assignment by the Hungarian algorithm on the confusion matrix, which resolves the label-switching ambiguity inherent to unsupervised partitions. We report the bootstrap distribution of the adjusted Rand index (ARI) between each resampled partition and the reference partition, together with per-cluster Jaccard indices, which quantify how reliably the membership of each individual cluster is reproduced across resamples and therefore distinguish globally stable partitions from partitions that are stable on average but contain one or two poorly resolved subgroups.

The main-text claims about prognosis and stability are backed by supplementary material on two parallel tracks. Full method-by-method benchmarks on the HNSC feature panel (multiRF versus SNF, MOFA2, iClusterPlus, and AJIVE) are reported in Supplementary Fig. 13 and Supplementary Table 12; cluster-by-cluster nested Cox model increments and the parallel progression-free interval analysis are reported in Supplementary Fig. 10 and Supplementary Tables 13 and 15; and bootstrap resampling of the shared clustering yielded mean ARI near 0.93 with the Atypical/HPV$^+$ cluster showing the highest per-cluster Jaccard index (Supplementary Fig. 6), which matters because HNSC is a noisy, heterogeneous disease in which purely expression-based subtype calls can vary across preprocessing pipelines and patient subsets.

#### 6 ADNI Supplementary Results

##### 6.1 Direct Cox Models: Incremental Value Over Clinical Covariates

Supplementary Table 18 reports an auxiliary direct-coordinate Cox benchmark, in which each method's learned low-dimensional coordinates are added directly to a covariate-only baseline of age, sex, education, MMSE, and APOE $\varepsilon4$; the parallel downstream leave-one-out cross-validated comparison after fixed representation learning appears in Supplementary Fig. 14. This direct-coordinate analysis tests whether the representation itself contains conversion-related information and is complementary to the main scalar age-acceleration benchmark in main-text Table 5. multiRF shared achieved the highest full-model $C$-index ($C_{\text{full}} = 0.704$, likelihood-ratio $p = 0.012$), followed by MRI PCA ($C_{\text{full}} = 0.688$, $p = 0.035$), multiRF specific(DNAm) ($C_{\text{full}} = 0.683$, $p = 0.180$), and SNF fused ($C_{\text{full}} = 0.683$, $p = 0.046$). AJIVE joint reached $C_{\text{full}} = 0.648$ ($p = 0.018$), and remaining methods did not improve on the covariate-only baseline of $C_{\text{cov}} = 0.623$.

##### 6.2 Ablation Study: Sensitivity to CpG Selection

We tested three CpG selection strategies on the DNAm layer with multiRF rerun 10 times per strategy under different random seeds. These were (i) AD-prioritized (3,503 CpGs from prior EWAS), (ii) brain-blood cross-tissue CpGs (831 CpGs with established brain-blood correlation), and (iii) union of both sets (4,332 CpGs, after two overlapping CpGs were counted once); full mean ± SD across age $R^2$, adjusted HR per one-year increase in age acceleration, and C-index is given in Supplementary Table 19, with parallel seed-stability plots for the survival C-index in Supplementary Fig. 15 and for age $R^2$ in Supplementary Fig. 16. This analysis examines whether the shared ADNI signal is directionally stable when the DNAm feature space is restricted to either component of the biologically motivated union panel. The union panel remains the primary feature set for the preclinical ADNI analysis because it combines AD-prioritized CpGs with CpGs supported by peripheral-to-brain methylation correspondence. All three strategies produced comparable results for the shared signal (HR 1.167–1.198, C-index 0.628–0.650). The brain-blood set achieved higher age $R^2$ (0.596) but slightly lower HR because restricting to CpGs with known brain-blood correlation improves age prediction at the cost of excluding some disease-relevant CpGs. The brain-blood shared signal was significant in 9 of 10 random seeds, compared with 10 of 10 for the union panel. The DNAm-specific signal showed more variability across CpG sets (HR 1.306–1.464, C-index 0.592–0.622), consistent with its role in capturing residual variance that depends on which CpGs enter the model.

##### 6.3 All-DNAm Sensitivity Analysis

The main ADNI comparison used the same 87 matched DNAm–MRI participants for every method to avoid sample-size confounding. Because this restriction excludes participants who had DNAm but no matched MRI, we repeated the DNAm-only baselines in all available baseline CN participants with DNAm and follow-up diagnosis ($n = 213$, 64 conversions to MCI or dementia). This analysis cannot evaluate multimodal methods, but it tests whether the DNAm-only clocks or DNAm PCA show stronger conversion associations when the full DNAm cohort is used. DNAm PCA on the same 4,332-CpG panel remained non-significant after covariate adjustment (HR = 1.066 per year of age acceleration, HR = 1.177 per SD, $p = 0.224$, $C = 0.673$; Supplementary Table 24). The established DNAm clocks and DunedinPACE also did not show significant adjusted associations ($p = 0.347$–$0.872$; Supplementary Table 24). Thus, the weaker DNAm-only results in the matched analysis are unlikely to be explained solely by excluding DNAm-only participants without MRI, although this sensitivity does not replace validation in an independent multimodal cohort.

##### 6.4 Quadrant Analysis: Agreement of Modality-Specific Age Acceleration

We stratified participants into four groups according to whether DNAm-specific and MRI-specific age acceleration lay above or below the cohort median. Kaplan–Meier curves for the two extreme quadrants showed that both-decelerated participants had markedly better conversion-free survival than both-accelerated participants (log-rank $p = 0.0057$; $n = 87$, 24 events; Supplementary Fig. 17). In a Cox regression with both-accelerated as the reference group and adjusting for age, sex, education, MMSE, and APOE $\varepsilon 4$ (Supplementary Table 20), DNAm-high accelerators showed an elevated conversion hazard (HR = 2.32, 95% CI 0.85–6.33, $p = 0.100$) and MRI-high accelerators showed a reduced but non-significant hazard (HR = 0.63, 95% CI 0.22–1.82, $p = 0.391$). The both-decelerated group contained no conversion events, so its adjusted Cox estimate is numerically unstable (point estimate $\approx 0$ with an infinite upper confidence bound) and the log-rank contrast of the two extremes is a more informative summary of that cell. The pattern indicates that DNAm-high acceleration flags a higher-risk pattern even though the small number of events in the off-diagonal cells limits precision.

##### 6.5 Covariate Associations of EVs

We examined associations between the shared and modality-specific EVs and baseline covariates (age, education, sex, APOE $\varepsilon 4$, MMSE) together with CSF biomarkers ($A\beta_{42}$, total tau, phospho-tau) available on the subset of participants with lumbar-puncture data (Supplementary Fig. 18; full per-EV statistics in Supplementary Table 21). On the shared decomposition, EV1 carried the main aging axis by construction (Pearson $r = 0.706$, $p = 2.2 \times 10^{-14}$), EV2 showed the strongest per-EV conversion association (age- and sex-adjusted HR = 0.48, $p < 0.05$), and EV5 correlated negatively with CSF $A\beta_{42}$ (Spearman $\rho = -0.479$, $p = 0.029$, $n = 21$). EV2 also showed sex and age covariate associations (sex Welch $p = 2.1 \times 10^{-4}$; age $r = -0.237$, $p = 0.027$), so it should be interpreted as a conversion-associated axis with residual demographic structure rather than as a pure disease-risk component. On the DNAm-specific decomposition, EV5 likewise tracked CSF $A\beta_{42}$ ($\rho = -0.514$, $p = 0.018$, $n = 21$) but no EV reached significance for age, which is the expected consequence of residualizing the shared aging axis prior to the modality-specific decomposition. On the MRI-specific decomposition, EV1 showed a borderline positive correlation with CSF phospho-tau ($\rho = 0.325$, $p = 0.047$, $n = 38$) and EV2 a comparable negative trend ($\rho = -0.328$, $p = 0.045$, $n = 38$), suggesting a small MRI-specific tauopathy signal that is not detectable in the unrestricted cohort. None of the EVs reached significance for MMSE (all $p > 0.25$), consistent with cognition being roughly preserved across this A−T− subset.

#### 6.6 Discrete Subtype Analysis

Beyond the continuous age-acceleration analyses, we performed an exploratory discrete clustering analysis on the multiRF shared similarity matrix, with the eigengap criterion selecting $k = 3$. Kaplan–Meier curves and per-cluster Cox coefficients are shown in Supplementary Fig. 19. The three resulting subtypes differed in age and sex distribution but not in baseline MMSE, education, APOE $\varepsilon 4$, or CSF biomarkers (Supplementary Table 16). In a multivariable Cox model adjusting for age, sex, education, MMSE, and APOE $\varepsilon 4$ (Supplementary Table 17; $n = 87$, 24 events, $C = 0.662$), the Cluster 3 versus Cluster 1 contrast was associated with conversion (HR = 4.54, 95% CI 1.09–18.84, $p = 0.037$), while Cluster 2 was numerically elevated but imprecise (HR = 2.85, 95% CI 0.72–11.31, $p = 0.136$). Because the event count is small and the clinical covariate coefficients are themselves imprecise in this subset, these cluster hazard ratios should not be interpreted as evidence that subtype membership outperforms or replaces standard clinical variables. Matched three-cluster partitions from SNF, MOFA2, and AJIVE produced qualitatively similar but method-dependent contrasts (SNF Cluster 1-vs-Cluster 3 HR = 4.95, $p = 0.015$; MOFA2 $p = 0.603$; AJIVE $p = 0.866$; Supplementary Table 22). We therefore treat the discrete subtype analysis as a secondary visualization of the shared aging structure rather than as a clinical prediction result.

#### 6.7 Robustness to Embedding Dimension

The primary analyses use the first five coordinates (EV1–EV5) for the biological-age regression. To verify that the conclusions do not hinge on this choice, we swept $k$ over $\{3, 4, \ldots, 10\}$ and refit the biological-age pipeline at each value (Supplementary Fig. 20; Supplementary Table 23). multiRF shared retained its adjusted hazard across the sweep (HR 1.18–1.21, significant for $k = 3$ to $k = 10$), and multiRF specific(DNAm) strengthened with increasing $k$ (HR 1.21, $p = 0.168$ at $k = 3$; HR 1.66, $p = 0.010$ at $k = 10$). SNF fused also improved modestly with $k$ (HR 1.10 at $k = 3$, HR 1.14, $p = 0.063$ at $k = 10$), but its cross-validated C-index remained below multiRF at the same $k$, consistent with SNF recovering signal only when given many components. The five-EV setting used in the main text is therefore a conservative choice that does not inflate the multiRF advantage.

[17] Trevor Hastie, Robert Tibshirani, and Jerome H. Friedman. *The Elements of Statistical Learning: Data Mining, Inference, and Prediction*. Springer, New York, 2nd edition, 2009.

[18] Steve Horvath. DNA methylation age of human tissues and cell types. *Genome Biology*, 14(10):R115, 2013.

[19] Hemant Ishwaran and Udaya B. Kogalur. *Fast Unified Random Forests for Survival, Regression, and Classification (RF-SRC)*, 2025. R package version 3.3.3.

[31] Bastian Pfeifer, Caterina Sirocchi, Marcus D. Bloice, Markus Kreuzthaler, and Martin Urschler. Federated unsupervised random forest for privacy-preserving patient stratification. *Bioinformatics*, 40(Supplement_2):ii198–ii207, 2024.

[46] Wei Zhang, Hanchen Huang, Lily Wang, Brian D. Lehmann, and X. Steven Chen. An integrative multiomics random forest framework for robust biomarker discovery. *GigaScience*, 15:giaf148, 2026.

#### Supplementary Figures

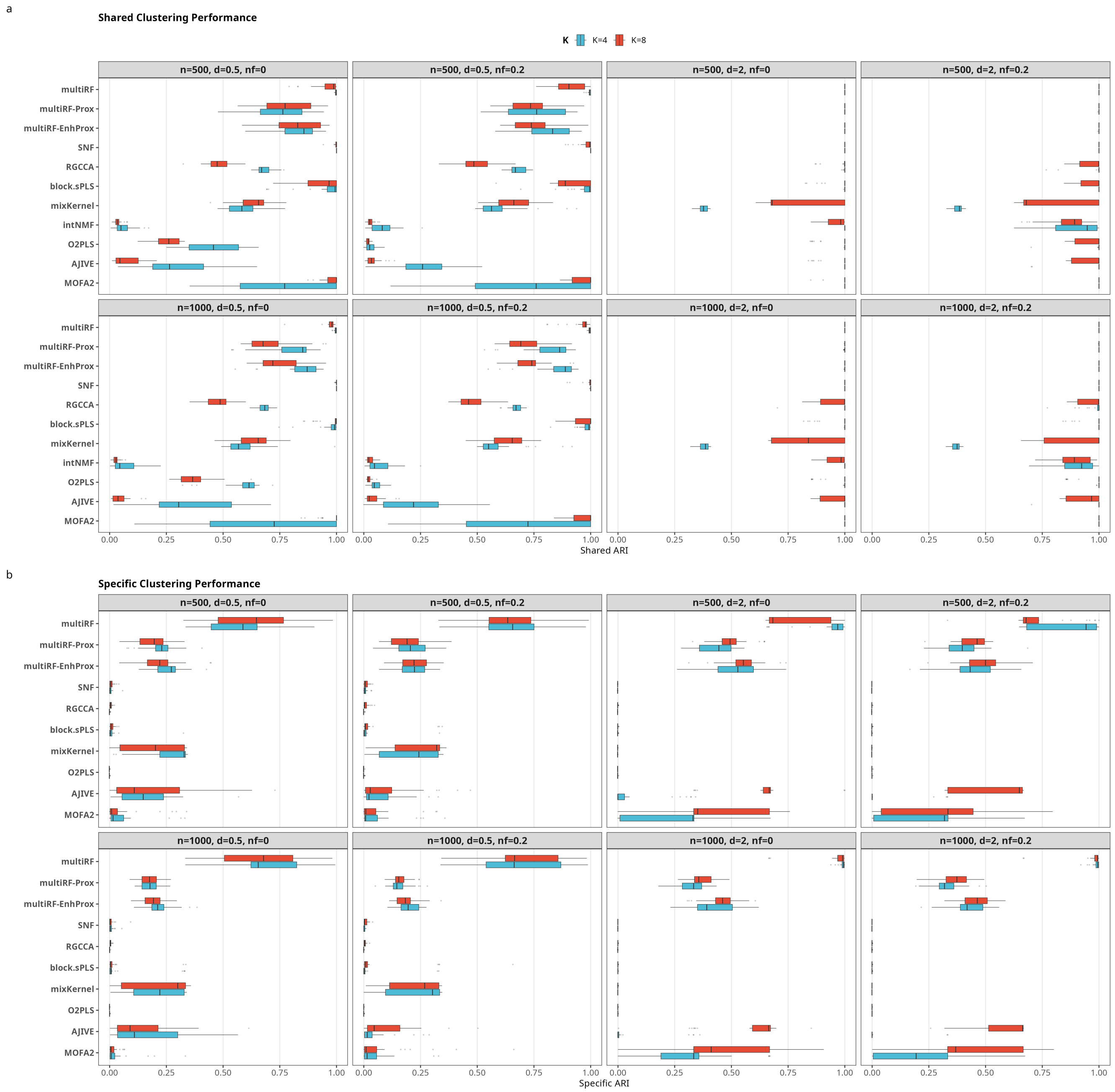

Supplementary Figure 1: InterSIM benchmark: full clustering comparison across all methods and scenarios. Boxplots summarize 30 replicates per condition, faceted by scenario parameters ($K$, $n$, $\delta$, noise fraction). **a** Shared clustering performance ($\mathrm{ARI}_Z$). **b** Specific clustering performance ($\mathrm{ARI}_U$). multiRF achieves near-perfect shared recovery across all conditions (mean $\mathrm{ARI}_Z = 0.989$) and is the only method to reliably detect modality-specific clusters (mean $\mathrm{ARI}_U = 0.785$).

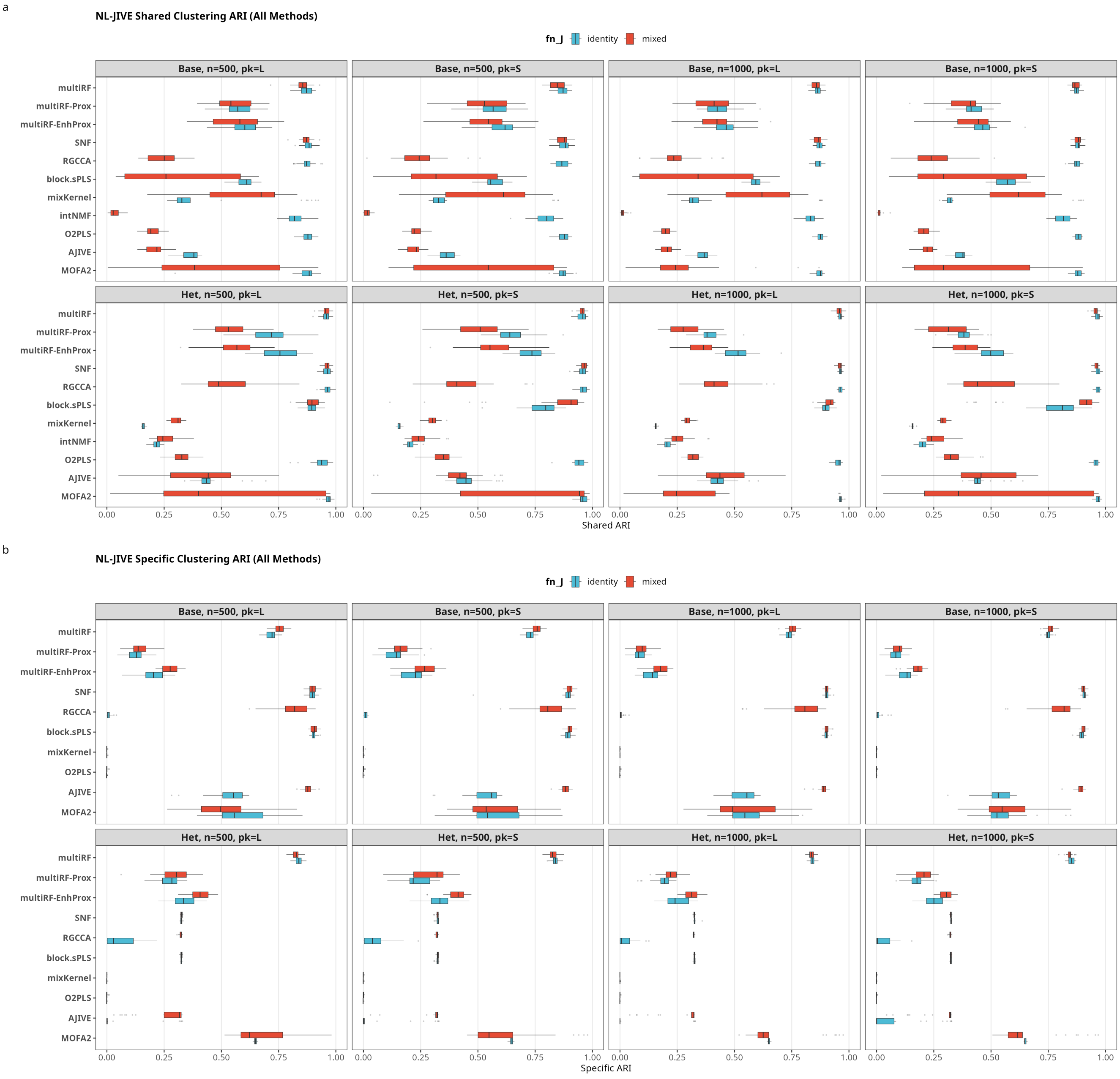

* Base: baseline homogeneous signal (dJ = dA).
* Het: heterogeneous individual signal with block-specific dA_k = 4/1/0.5.

Supplementary Figure 2: NL-JIVE benchmark: full clustering comparison across all methods and nonlinearity regimes. Boxplots summarize 30 replicates per condition, faceted by $f_J$ (identity, mixed), sample size $n$, and feature dimension $p_k$. **a** Shared clustering performance ($\mathrm{ARI}_Z$). **b** Specific clustering performance ($\mathrm{ARI}_U$). Under the mixed nonlinearity, linear factor methods (RGCCA, O2PLS, intNMF) collapse while multiRF and SNF remain stable.

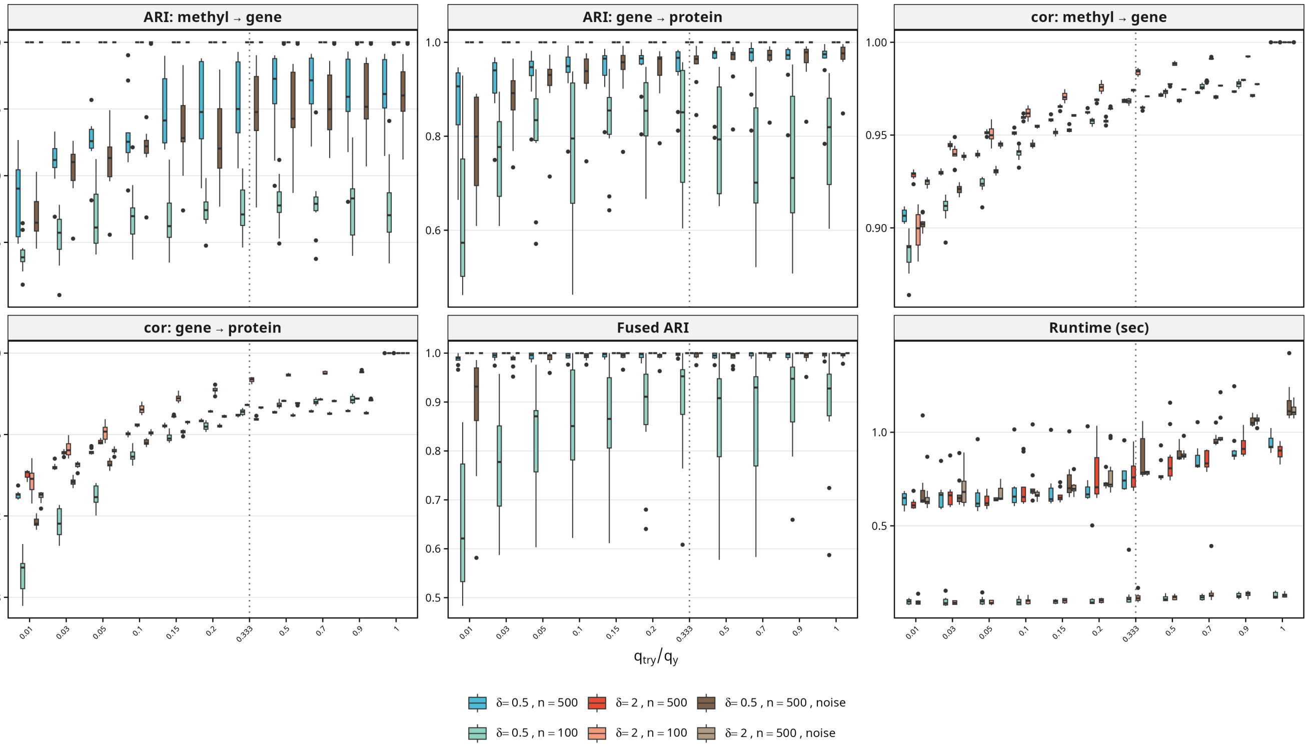

Supplementary Figure 3: Sensitivity of multiRF to the response subsampling fraction $q_{\text{try}}/q_y$. Boxplots summarize 10 replicates per scenario. The six panels report (top row, left to right) per-forest clustering ARI for the methyl → gene and gene → protein connections, and Pearson correlation of each per-forest weight matrix with the full-response reference; (bottom row) correlation for gene → protein, fused ARI after uniform-weight fusion of both connections, and wall-clock runtime for forest fitting. The dotted vertical line marks the default $q_{\text{try}} = \lceil q_y/3 \rceil$. Six scenarios are shown. $\delta \in \{0.5, 2.0\}$ (signal strength) crossed with $n \in \{500, 100\}$ (sample size), plus two noise conditions ($n = 500$) in which 50% i.i.d. $\mathcal{N}(0, 1)$ columns are appended to every block.

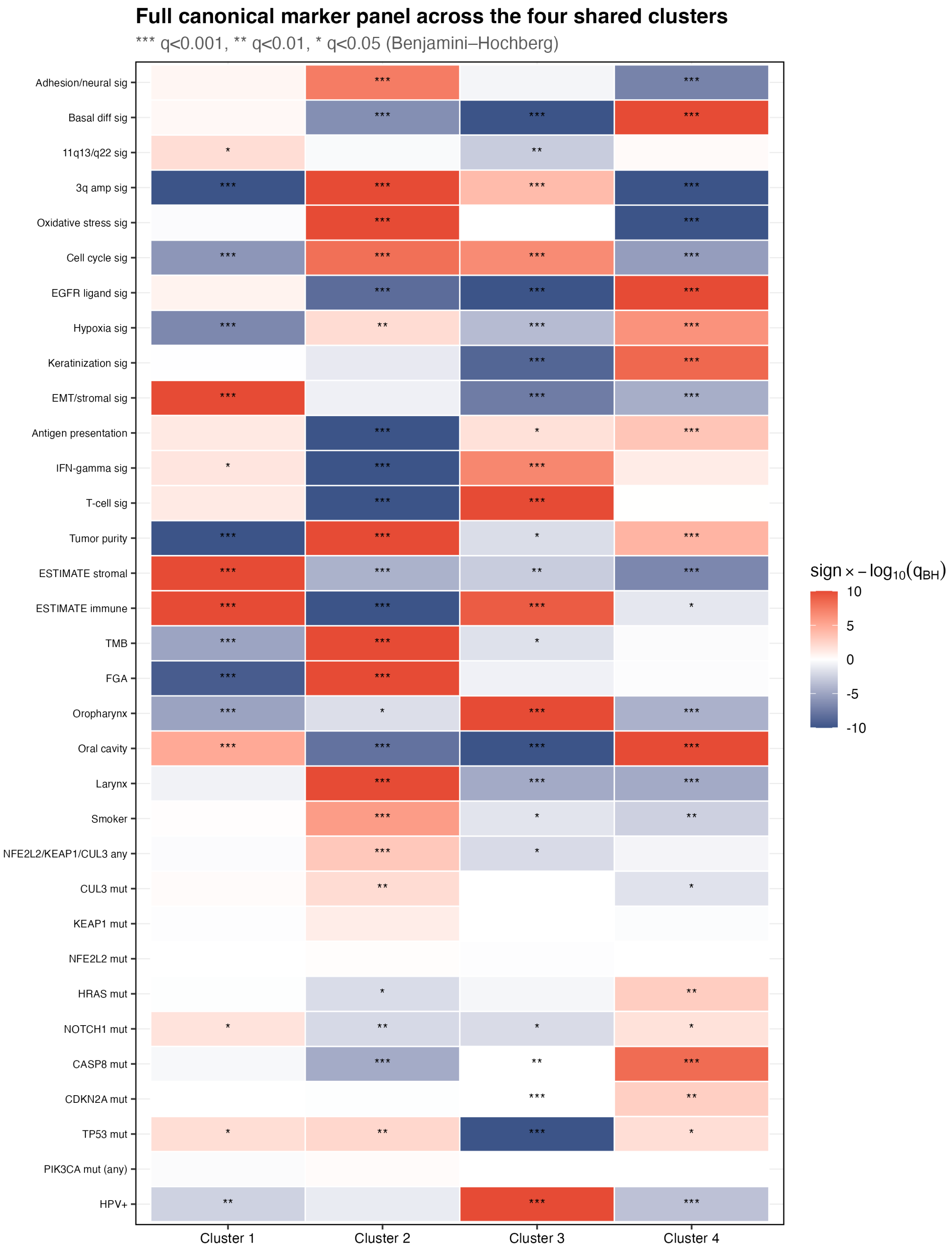

Supplementary Figure 4: Full canonical-marker panel across the four shared clusters. Heatmap cells encode signed $-\log_{10} q_{\mathrm{BH}}$ from one-vs-rest Fisher (binary) or Wilcoxon (continuous) tests; asterisks denote $q < 0.05/0.01/0.001$. Markers include HPV status; *TP53*, *CDKN2A*, *NOTCH1*, *PIK3CA*, *NFE2L2* / *KEAP1* / *CUL3*, *HRAS*, *CASP8* mutations; anatomic subsite; smoking; FGA; TMB; ESTIMATE immune, stromal, and tumor purity; xCell T-cell, IFN-$\gamma$, antigen-presentation; and canonical gene-expression signatures (keratinization, EMT/stromal, cell cycle, hypoxia, EGFR ligand).

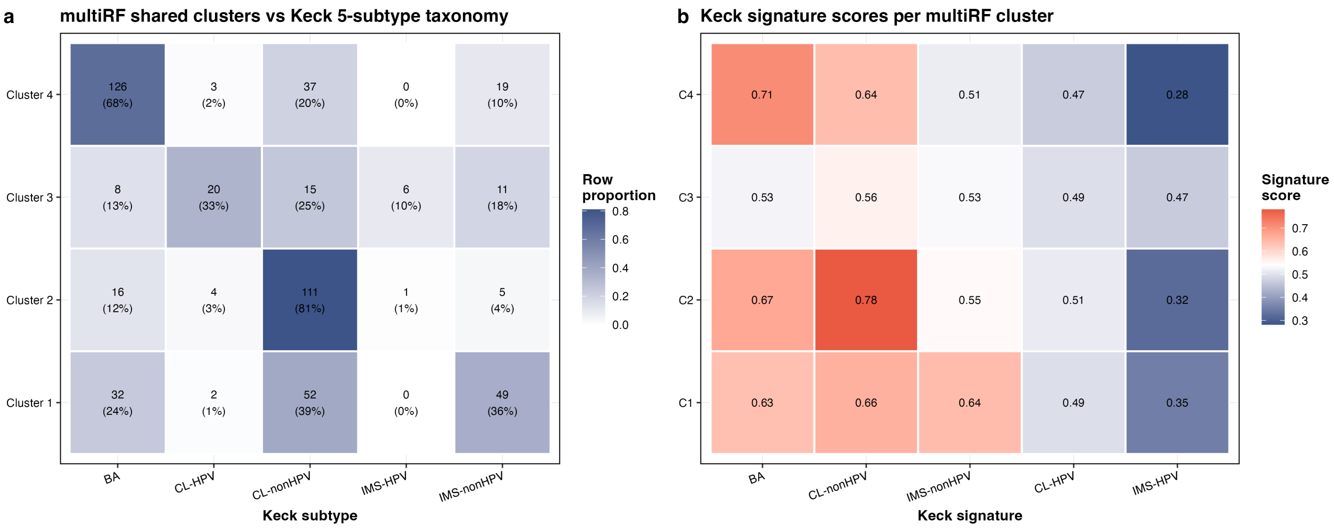

Supplementary Figure 5: Keck 5-subtype crosswalk. **a** Contingency heatmap of multiRF shared clusters against the Keck 5-subtype taxonomy (BA, CL-HPV, CL-nonHPV, IMS-HPV, IMS-nonHPV); cell annotations show counts and row proportions. **b** Keck signature scores (z-scored) per multiRF cluster, indicating the main Keck assignment for each cluster.

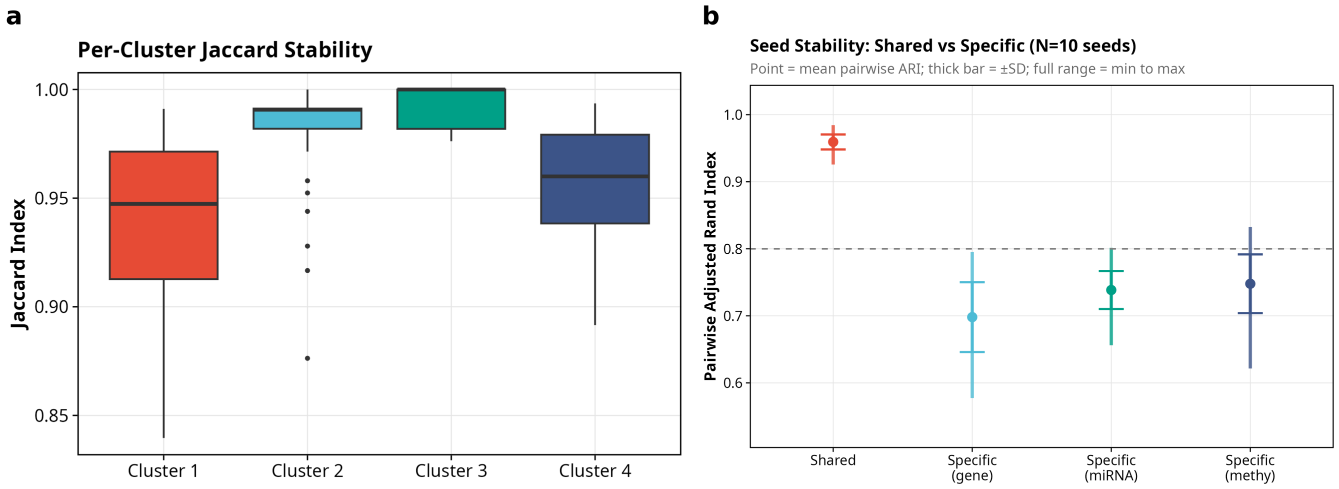

Supplementary Figure 6: HNSC multiRF stability. **a** Per-cluster bootstrap Jaccard index across 100 resamples of the shared multiRF similarity matrix. Clusters 2 and 3 attain near-perfect stability (median Jaccard near 0.99), and Clusters 1 and 4 retain median Jaccard $\geq 0.95$. **b** Pairwise adjusted Rand index across 10 independent random seeds for the shared clustering and for each omic-specific clustering. Points denote the mean pairwise ARI, thick bars $\pm 1$ SD, and thin whiskers the min–max range. The shared solution is highly reproducible (mean ARI near 0.96), whereas the gene-, miRNA- and methylation-specific solutions show markedly larger seed variability (mean ARI in the 0.70–0.75 range), consistent with the interpretation that the specific axes capture a weaker, more exploratory signal than the shared consensus. The dashed reference line marks ARI = 0.80.

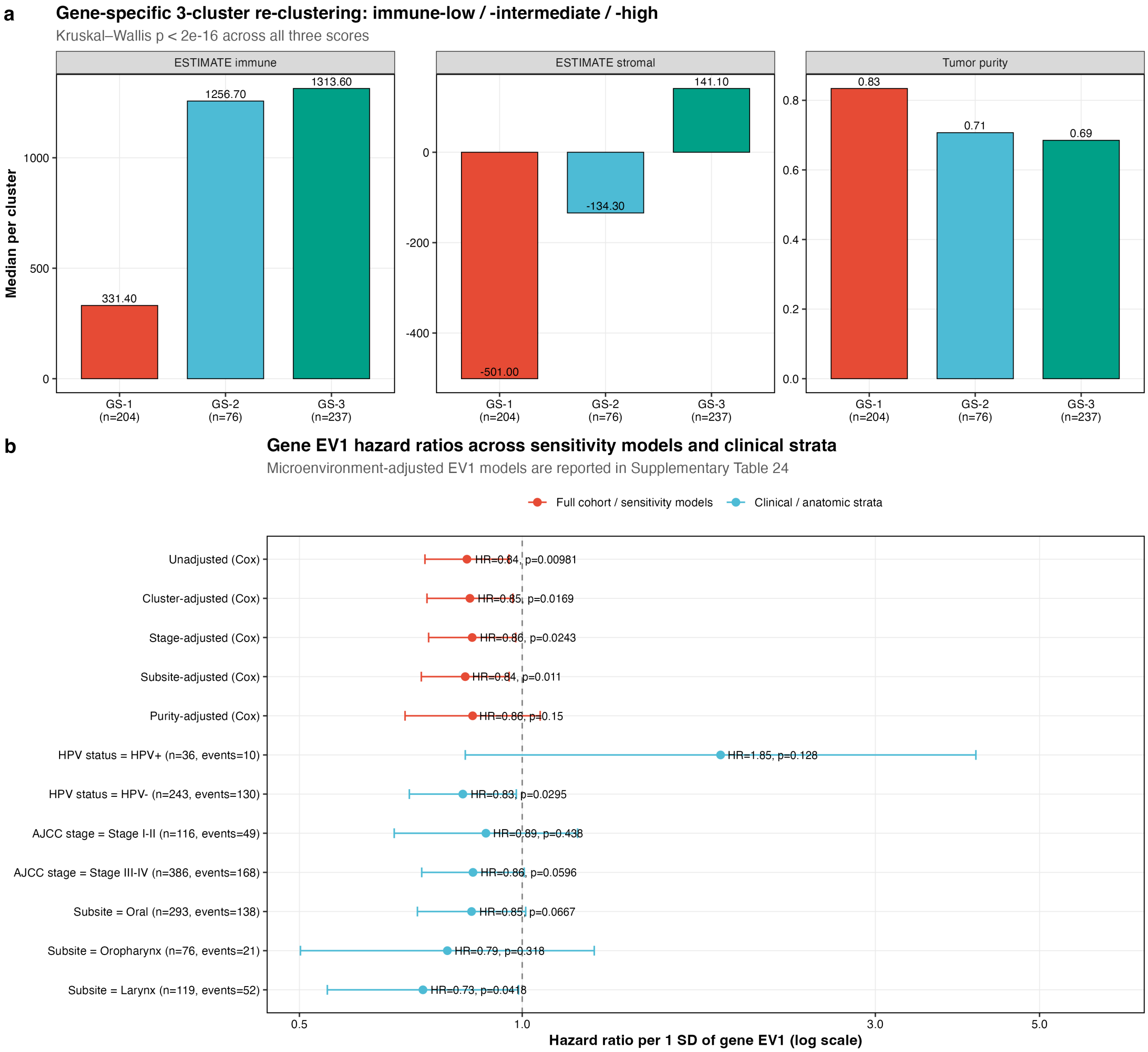

Supplementary Figure 7: Gene-specific axis: 3-cluster re-clustering and stratified hazard ratios. **a** Gene-specific spectral re-clustering ($k = 3$) produces immune-low (GS-1), -intermediate (GS-2), and -high (GS-3) groups with ESTIMATE immune medians of 330, 1,260, and 1,310, respectively (Kruskal–Wallis $p < 2 \times 10^{-16}$ for immune, stromal, and tumor-purity axes). **b** Gene EV1 hazard ratios for overall survival across sensitivity models (unadjusted, cluster-adjusted, HPV-adjusted, tumor-purity-adjusted, stage- and subsite-adjusted) and across clinical strata (HPV status, pathologic stage, anatomic subsite). Expanded microenvironment-adjusted Cox models for all three modality-specific EV1s are reported in Supplementary Table 14.

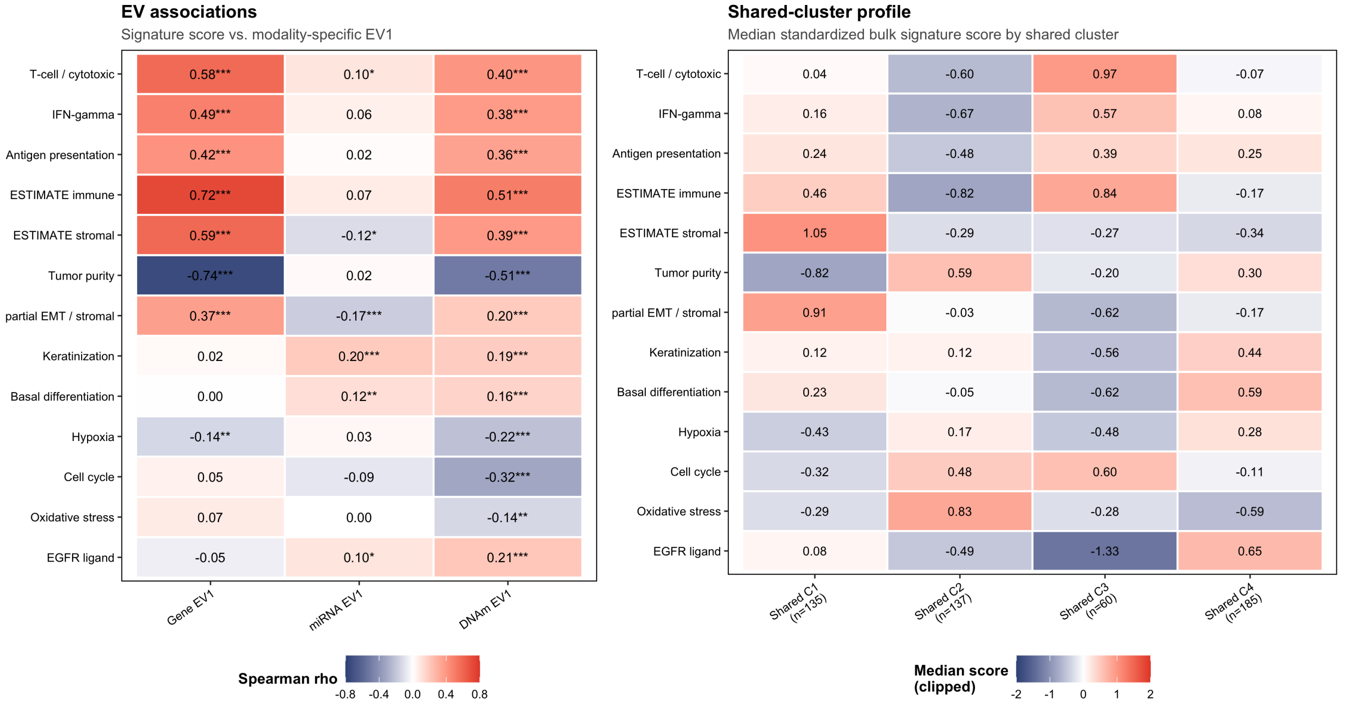

Supplementary Figure 8: Puram-related bulk signature analysis in TCGA HNSC. **a** Spearman correlations between targeted bulk signatures related to single-cell HNSCC programs described by Puram et al. [61] and the leading modality-specific axes from MULTIRF. Asterisks denote BH-FDR thresholds across the full-cohort signature–EV tests ($q < 0.05/0.01/0.001$). **b** Median standardized bulk signature score by MULTIRF shared cluster. The analysis is intended as bulk-signature contextualization of the shared and specific axes; it is not a single-cell or spatial state decomposition.

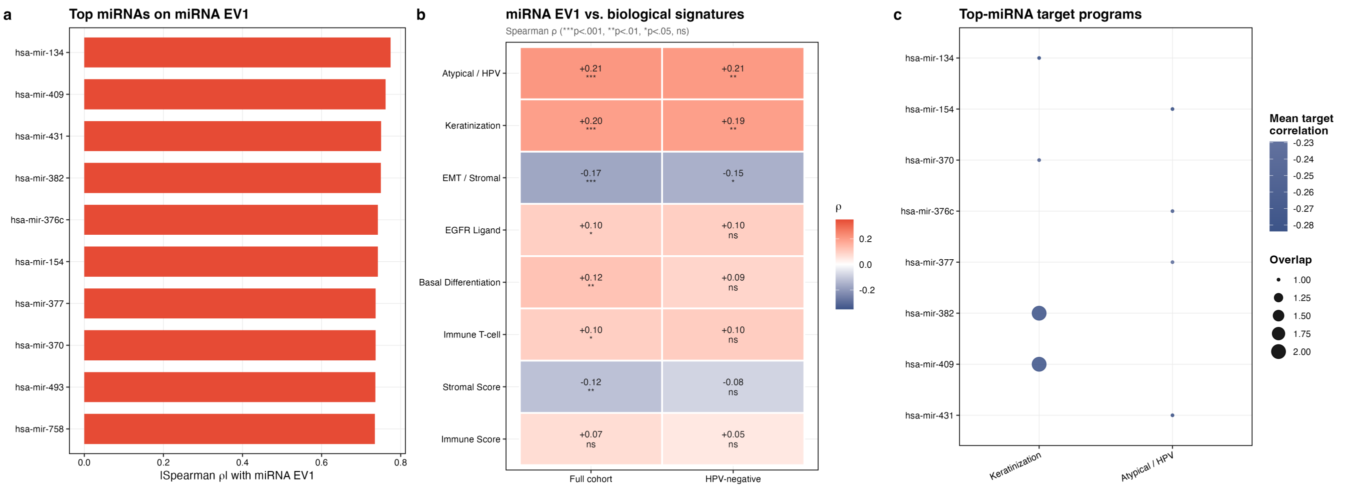

Supplementary Figure 9: miRNA-specific biology in TCGA HNSC. **a** Top miRNAs defining the miRNA-specific leading axis, ranked by absolute Spearman correlation with miRNA EV1. **b** Association of miRNA EV1 with targeted biological signatures in the full cohort and in HPV-negative tumors. The main pattern links the miRNA-specific component to keratinization, atypical / HPV-related features, and weaker basal / EGFR-ligand biology rather than to the immune residual. **c** Putative target-program summary for the highest-weight miRNAs, based on negatively correlated target genes. Together these panels support the interpretation that miRNA-specific variation reflects post-transcriptional epithelial-state variation distinct from the shared subtype structure.

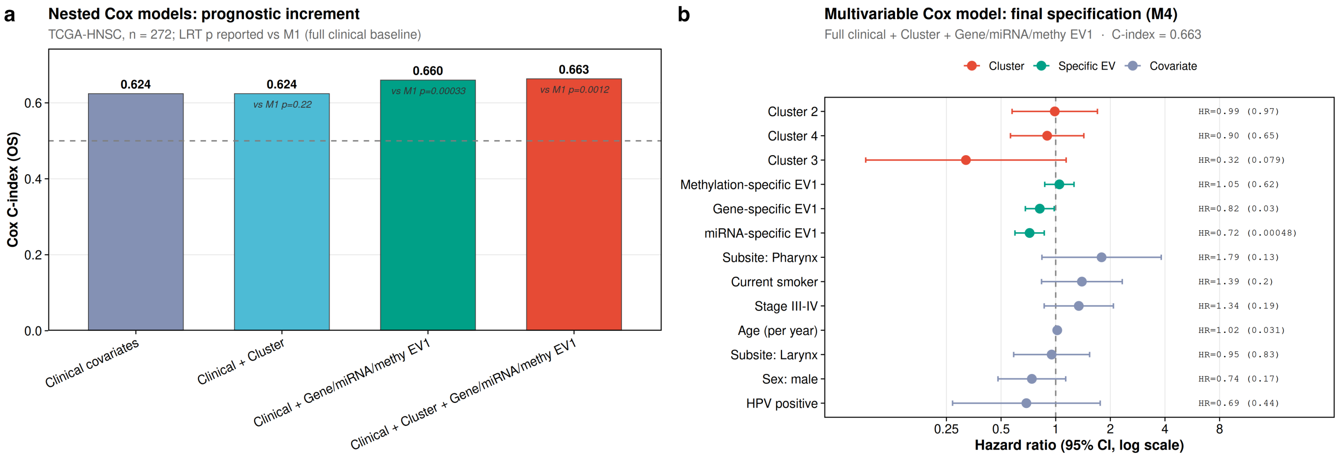

Supplementary Figure 10: Nested Cox survival modeling for HNSC overall survival. **a** Harrell's $C$-index across four nested Cox models fit on the complete-case TCGA HNSC survival subset ($n = 272$): M1, clinical covariates alone (age, sex, HPV status, stage, anatomic subsite, and smoking status); M2, M1 plus the four-cluster shared assignment; M3, M1 plus the gene-, miRNA- and methylation-specific leading axes (EV1 per modality) without cluster; M4, the full model combining clinical covariates, cluster assignment, and all three modality-specific axes. Incremental $C$-index gains across the M1→M2→M4 sequence quantify the survival information added by the shared clustering and by each modality-specific axis beyond clinical variables. **b** Forest plot of the final (M4) Cox coefficients reported as hazard ratios with 95% confidence intervals. Cluster dummies are referenced against Cluster 1 (Mesenchymal / immune-like); modality axes are standardized so that hazard ratios correspond to a one-SD increase. Numerical estimates also appear in Supplementary Table 13.

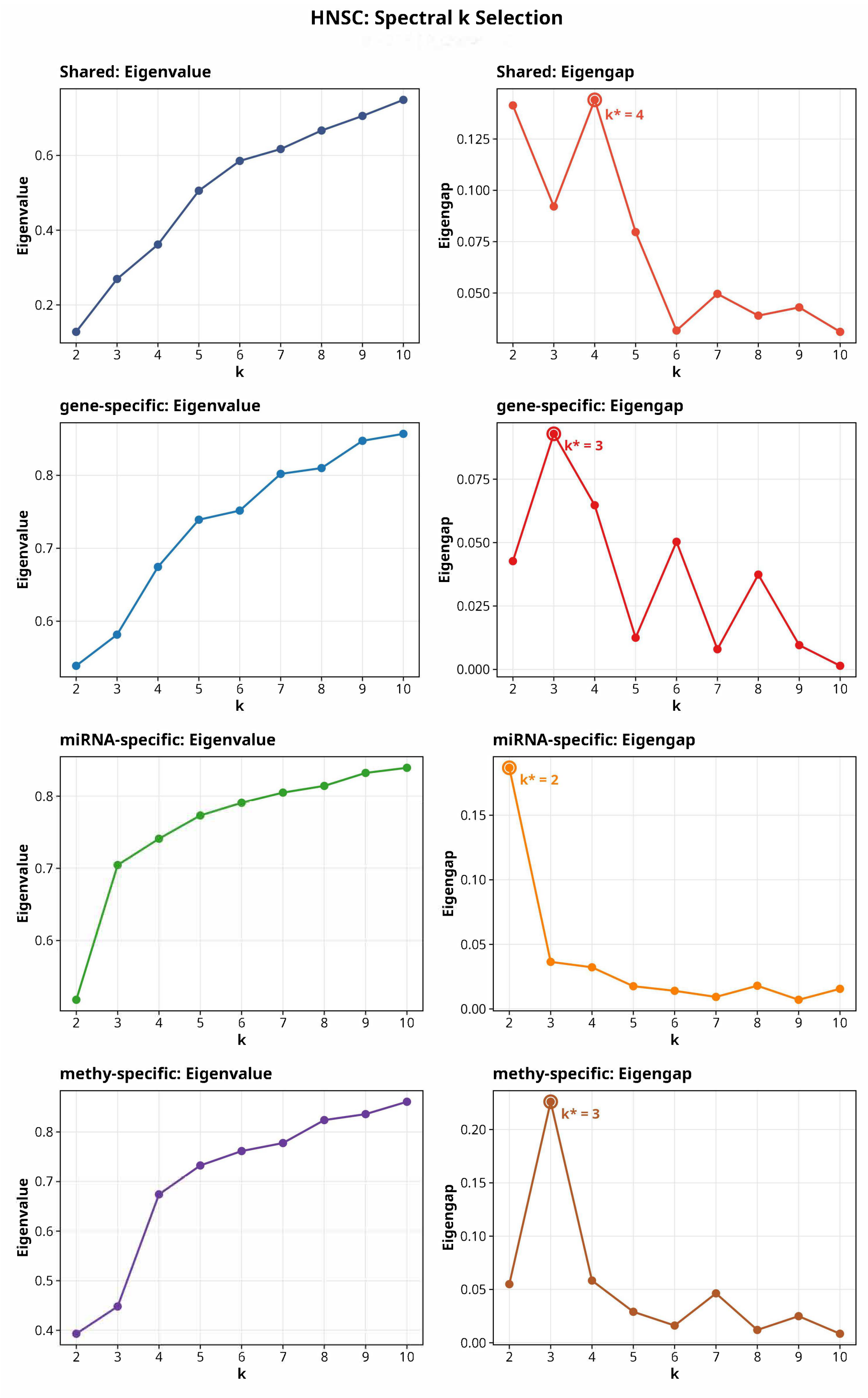

Supplementary Figure 11: HNSC spectral $k$ selection across the shared and component-specific multiRF similarity matrices. For each row the left panel shows the leading eigenvalues of the normalized similarity matrix and the right panel shows the eigengap $\Delta\lambda_k = \lambda_{k+1} - \lambda_k$, with the eigengap-maximizing $k^*$ circled and annotated. The shared similarity matrix supports $k_{\text{shared}} = 4$, and the gene-, miRNA- and methylation-specific matrices each contribute their own $k^*$ used by the specific-axis analyses.

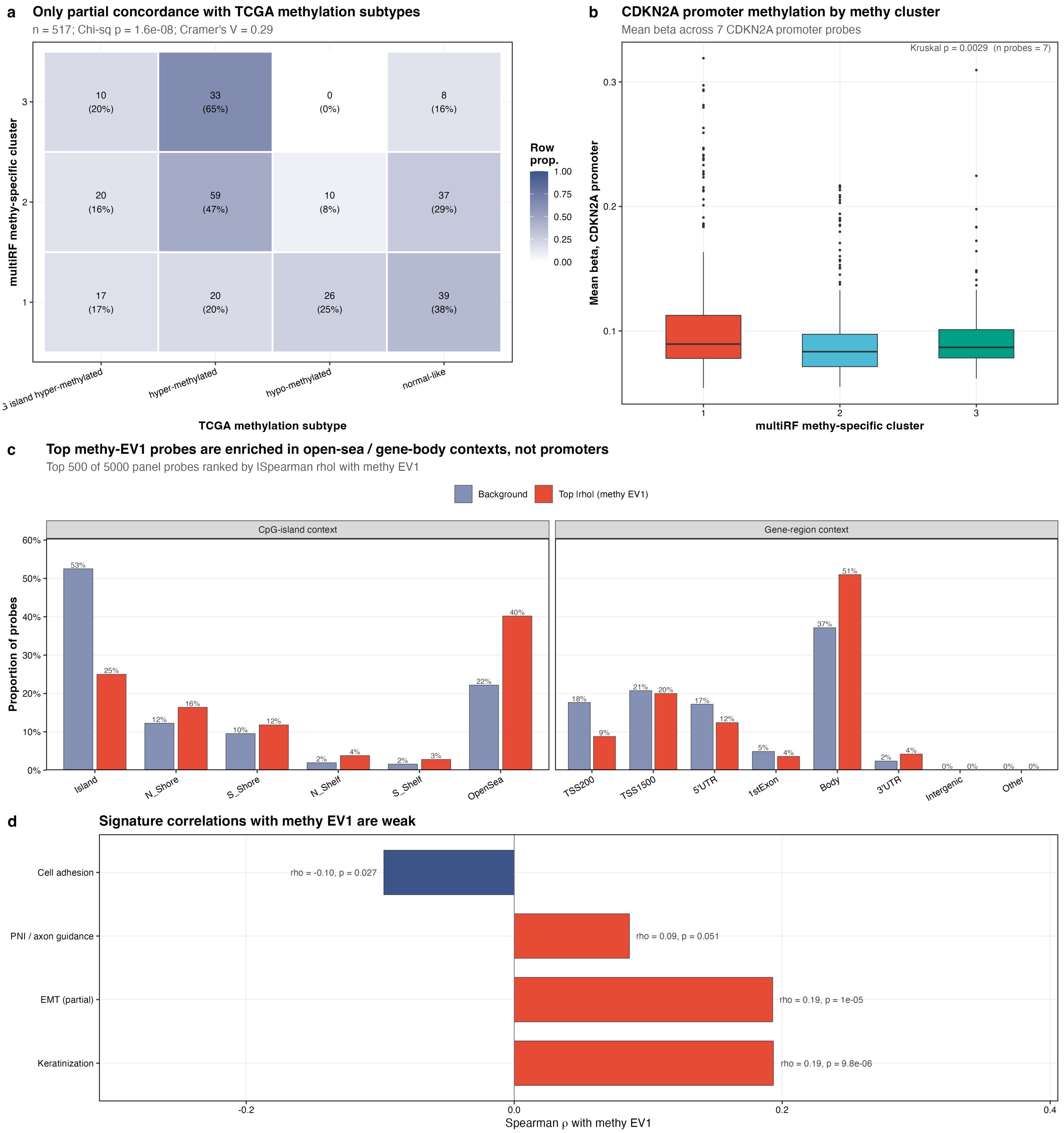

Supplementary Figure 12: Methylation-specific structure in TCGA HNSC. **a** Contingency heatmap comparing the three methylation-specific clusters with the TCGA methylation subtype labels. **b** CDKN2A promoter methylation by methy-specific cluster. **c** Genomic-region distribution of top methy-EV1 probes vs background panel. **d** Signature correlations with methy EV1.

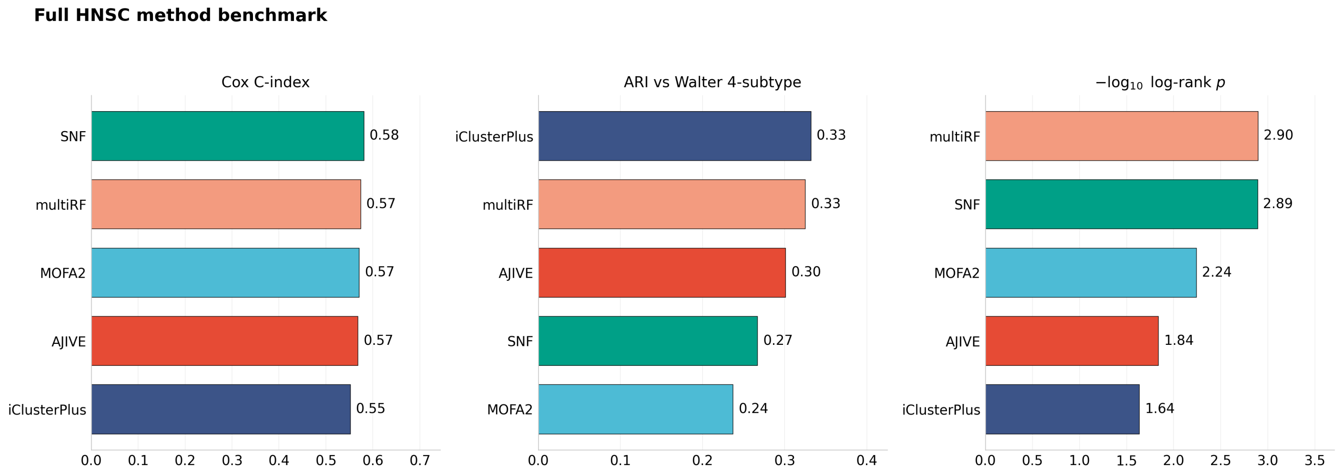

Supplementary Figure 13: HNSC method benchmark detail. Cox $C$-index, adjusted Rand index against the Walter 4-subtype reference, and $-\log_{10}$ log-rank $p$ for the five evaluated methods (multiRF, SNF, MOFA2, AJIVE, iClusterPlus). Values correspond to those in the main-text HNSC method-comparison table and Supplementary Table 12.

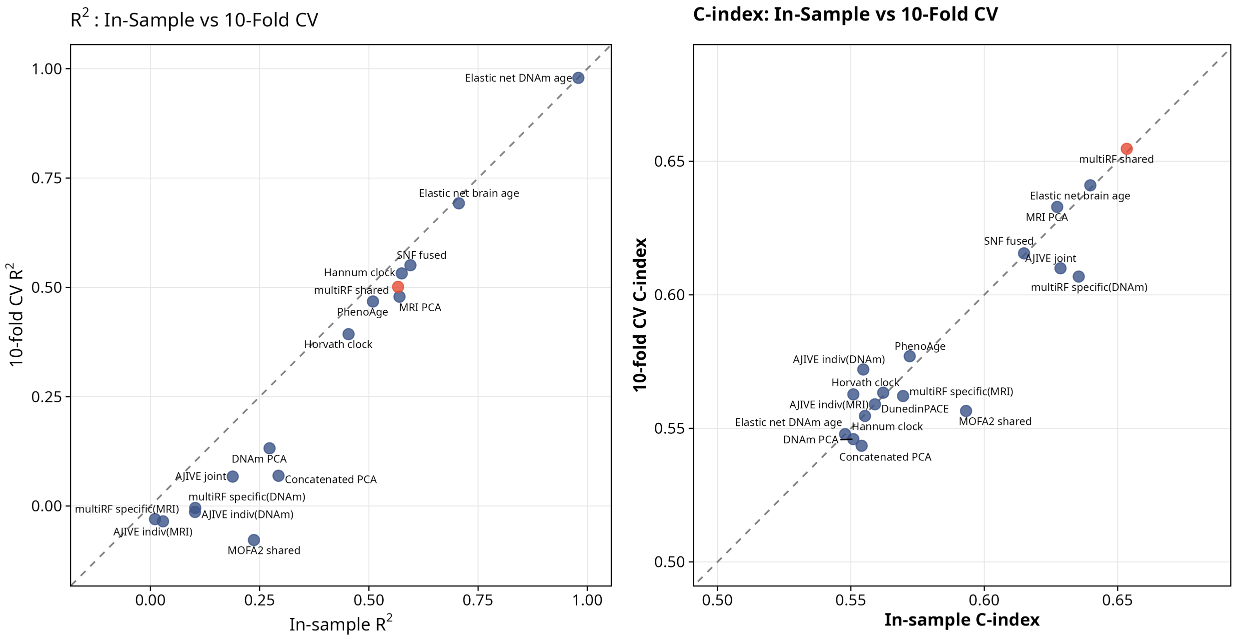

Supplementary Figure 14: Downstream LOO-CV performance after fixed representation learning. **a** CV $R^2$ vs. in-sample $R^2$ for all methods; points near the diagonal indicate stable generalization. **b** CV $C$-index vs. in-sample $C$-index; multiRF shared retains its advantage under downstream LOO-CV.

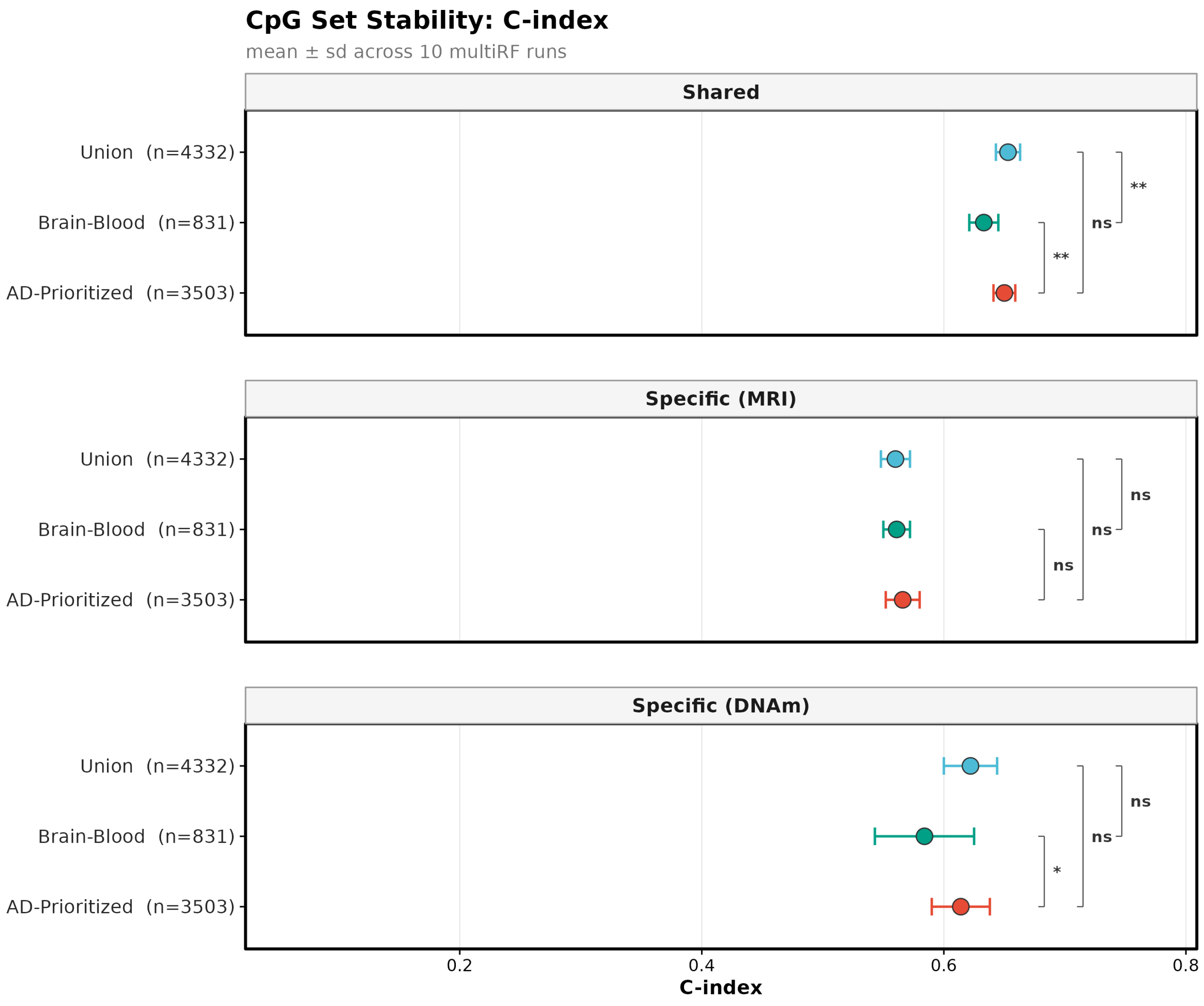

Supplementary Figure 15: CpG set stability: $C$-index across 10 random seeds for three CpG selection strategies (AD-prioritized, brain-blood, union) and three signal types (shared, specific DNAm, specific MRI). Error bars show mean ± SD.

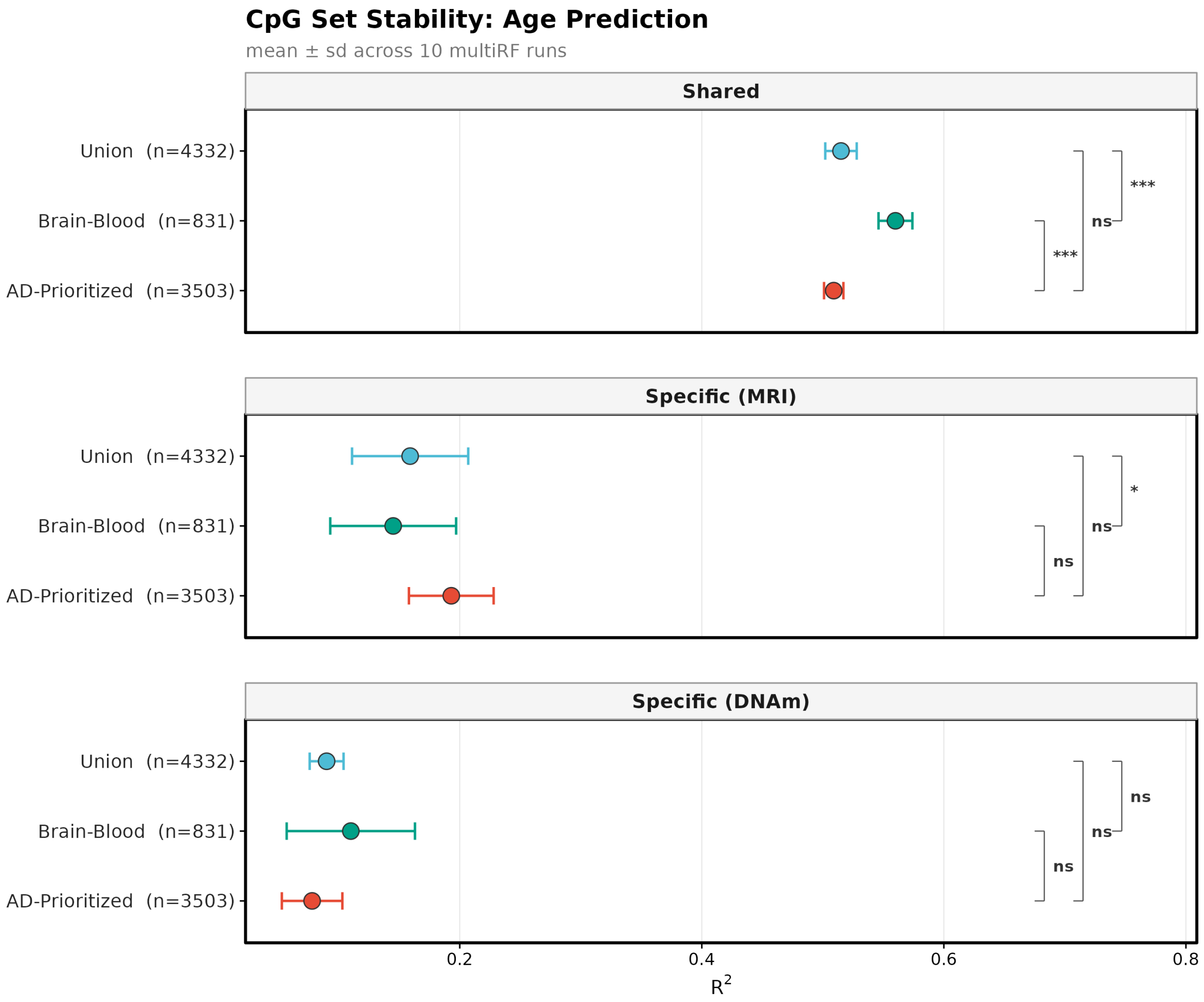

Supplementary Figure 16: CpG set stability: age prediction $R^2$ across 10 random seeds for three CpG selection strategies and three signal types.

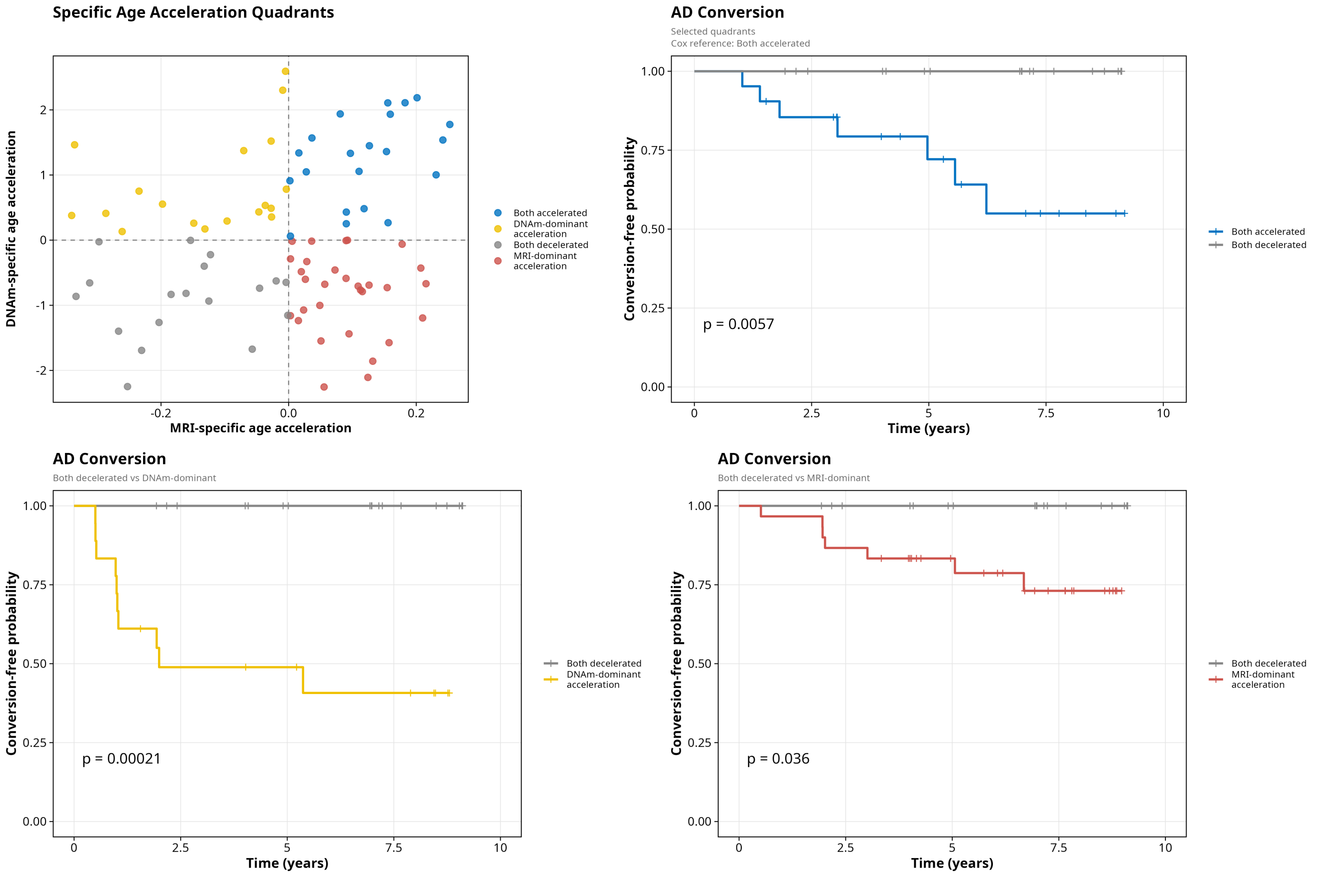

Supplementary Figure 17: Quadrant analysis of modality-specific age acceleration. Scatter plot of DNAm-specific vs. MRI-specific age acceleration colored by quadrant, with Kaplan–Meier curves for extreme quadrants (both accelerated vs. both decelerated; log-rank $p = 0.0057$).

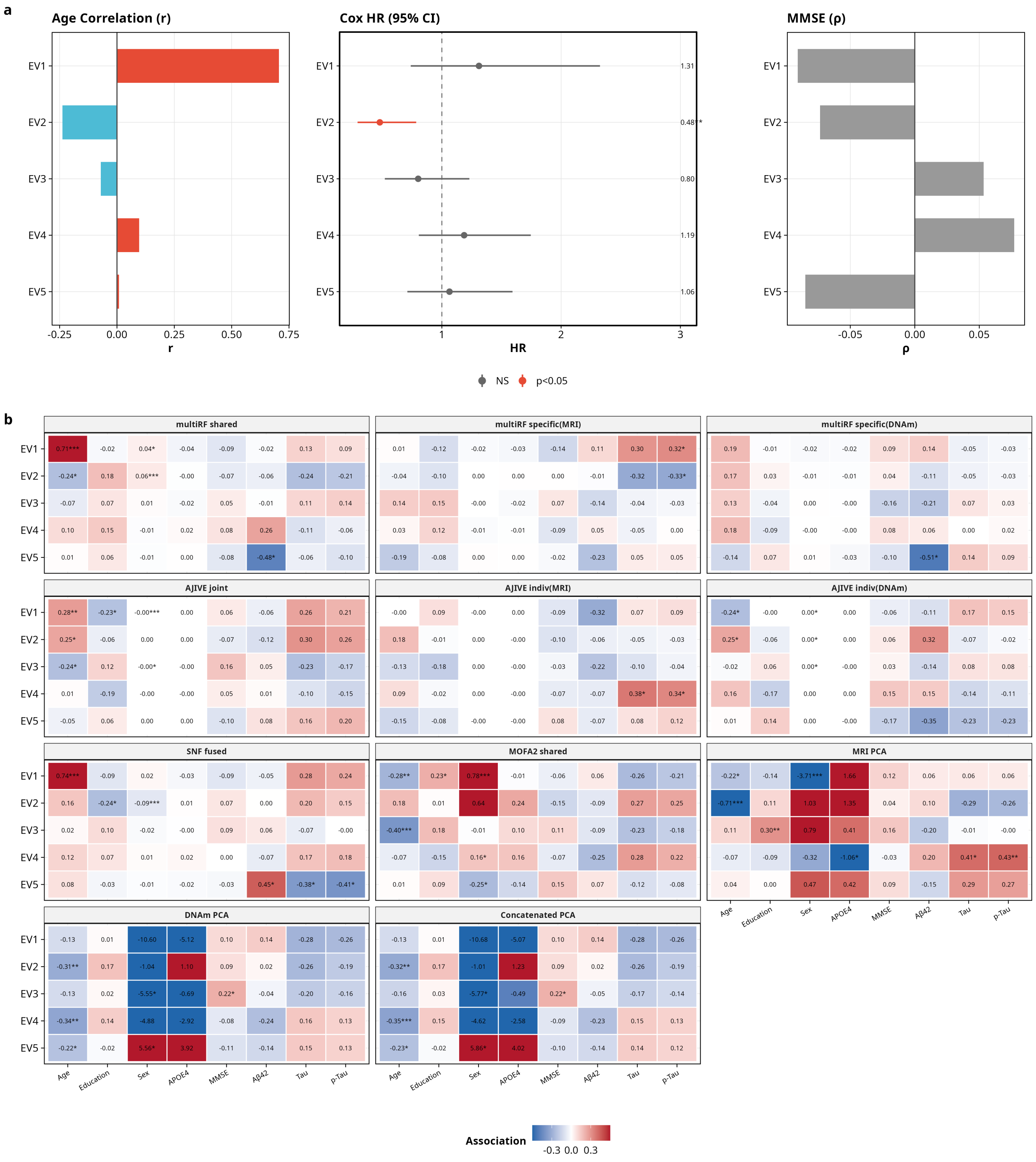

Supplementary Figure 18: Exploratory per-eigenvector analysis of ADNI multimodal representations. **a** For multiRF shared, per-EV summary panels: (left) Pearson correlation with chronological age, (center) age- and sex-adjusted Cox hazard ratio for MCI/AD conversion with 95% confidence intervals, and (right) Spearman correlation with baseline MMSE. EV1 carries the main chronological-age signal (Pearson $r = 0.706$, $p = 2.2 \times 10^{-14}$). EV2 reaches nominal significance in the per-EV Cox model (HR $= 0.48$, $p < 0.05$, age- and sex-adjusted) and also shows sex and age covariate associations (sex Welch $p = 2.1 \times 10^{-4}$; age $r = -0.237$, $p = 0.027$); the negative direction of the Cox HR indicates that, within this small cohort, larger EV2 scores are associated with lower conversion hazard. EV5 aligns with baseline CSF A$\beta_{42}$ (Spearman $\rho = -0.479$, $p = 0.029$, $n = 21$). None of the per-EV MMSE correlations reach significance. **b** Cross-method per-EV heatmap of associations with clinical covariates and CSF biomarkers. Values are Pearson $r$, Spearman $\rho$, or mean difference depending on variable type; asterisks denote nominal significance. These patterns are exploratory because the matched ADNI cohort contains only 87 participants and 24 conversion events.

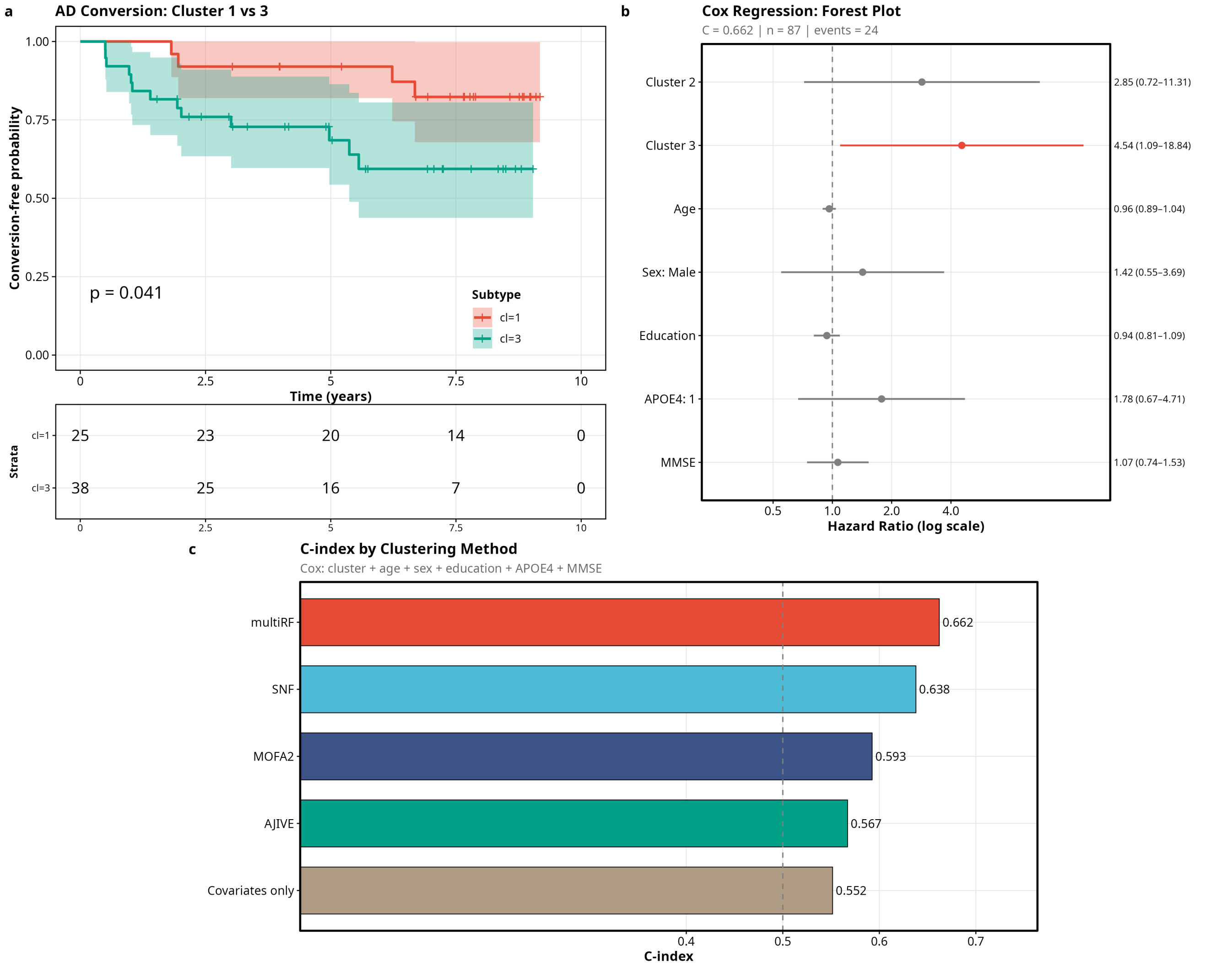

Supplementary Figure 19: Exploratory discrete subtype analysis from multiRF shared clustering in ADNI. **a** Kaplan–Meier curves for conversion to MCI or AD comparing Cluster 1 and Cluster 3. **b** Adjusted Cox coefficients for cluster membership with age, sex, education, APOE4, and MMSE. **c** $C$-index comparison across clustering methods. The cluster hazard-ratio estimates have wide confidence intervals and are interpreted as hypothesis-generating rather than as a clinical risk classifier.

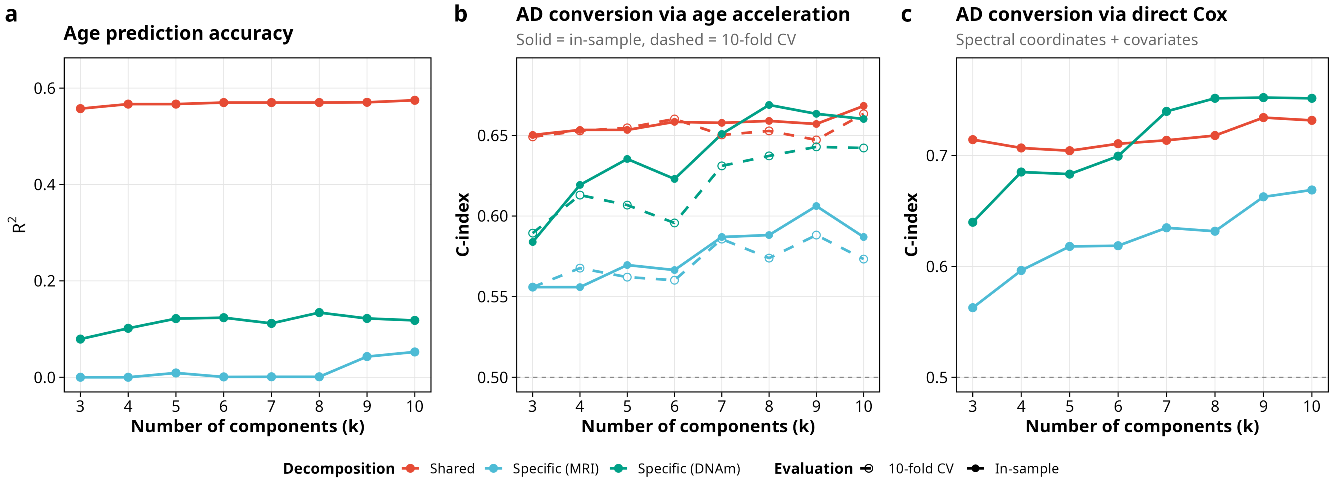

Supplementary Figure 20: Embedding-dimension sweep for the continuous age-acceleration pipeline. **(a)** Age-prediction $R^2$ as a function of the number of EVs $k \in \{3, 4, \ldots, 10\}$ for MULTIRF shared, specific(DNAm), and specific(MRI) biological ages. **(b)** In-sample and 10-fold cross-validated $C$-index for the age-acceleration Cox model (cluster-free, pipeline-A: decomposition → age prediction → age acceleration → Cox) at each $k$. **(c)** $C$-index for the direct-Cox pipeline (pipeline-B: EVs $E^{(\cdot)}$ entered directly as covariates alongside clinical adjusters). Shared decomposition retains stable discrimination across the entire range; DNAm-specific biological-age acceleration strengthens monotonically with $k$. Corresponding numerical values are in Supplementary Table 23.

### Supplementary Tables

Supplementary Table 1: Summary of methods. Summary of all methods evaluated in the experiments. Three multiRF clustering variants are compared: multiRF (the default similarity-based pipeline), multiRF-Prox (proximity-based), and multiRF-EnhProx (enhanced proximity with soft weighting). All baseline methods use their recommended default settings.

Supplementary Table 2: Simulation design. Design parameters for the InterSIM and NL-JIVE simulation studies. The InterSIM sheet lists all 16 factorial scenarios with per-block feature dimensions; $\delta$ controls the separation between cluster centroids, $K_Z$ is the number of shared clusters, and $K_U$ is the number of view-specific clusters. The NL-JIVE sheet lists 16 scenarios (8 Baseline + 8 Hetero); $\delta_J$ is the joint (shared) signal strength common across all views, and $\delta_A$ per view gives the individual (view-specific) signal strengths. Baseline uses homogeneous view-specific signals ($\delta_A = 2$ for all views), while Hetero uses heterogeneous signals ($\delta_A = 4, 1, 0.5$). Each scenario is replicated 30 times.

Supplementary Table 3: Simulation benchmark results. Clustering performance (mean ± SD across 30 replicates) for the InterSIM and NL-JIVE simulations. Shared ARI measures recovery of the shared cluster structure; Specific ARI measures recovery of the view-specific structure. For InterSIM, the number of shared clusters is $K_Z \in \{4, 8\}$ and the number of view-specific clusters is $K_U = 2$. For NL-JIVE, the shared structure has $K_Z = 3$ clusters and each view has $K_U = 2$ view-specific clusters. Two NL-JIVE signal settings are evaluated: Baseline ($\delta_J = 2$, $\delta_A = 2/2/2$) and Hetero ($\delta_J = 2.5$, $\delta_A = 4/1/0.5$). Runtime is the median wall-clock time in seconds.

Supplementary Table 4: Shared-cluster alignment with established HNSCC subtype systems. The table reports the contingency of multiRF shared clusters against the TCGA / Walter 4-subtype reference and the pairwise Fisher / Cramér's $V$ agreement statistics.

Supplementary Table 5: Full canonical-marker test results for the four HNSC shared clusters. For each cluster × marker pair we report the test used (Fisher / Wilcoxon), counts, effect size (odds ratio or median difference), test statistic, raw $p$-value, BH-adjusted $q$-value, and signed $-\log_{10}(q)$ used to color Supplementary Fig. 4.

Supplementary Table 6: Keck 5-subtype crosswalk. Contingency of multiRF shared clusters against the Keck 5-subtype taxonomy (BA, CL-HPV, CL-nonHPV, IMS-HPV, IMS-nonHPV) with per-cluster counts and row proportions, and Keck signature scores (BA, CL-nonHPV, IMS-nonHPV, CL-HPV, IMS-HPV) per multiRF cluster.

Supplementary Table 7: Gene EV1 stratified hazard ratios. Cox proportional-hazards estimates (HR, 95% CI, $p$-value) for gene EV1 as a continuous predictor under a sequence of sensitivity models (unadjusted, cluster-adjusted, HPV-adjusted, tumor-purity-adjusted, stage- and subsite-adjusted) and across clinical strata (HPV status, pathologic stage, anatomic subsite), summarizing Supplementary Fig. 7b.

Supplementary Table 8: miRNA-specific leading-axis program overlap. Hypergeometric enrichment of the top miRNA EV1 contributors against four reference programs (DLK1–DIO3 imprinted cluster, C19MC trophoblast cluster, HPV-responsive, miR-200), reporting program size, overlap, fold enrichment, hypergeometric $p$-value, BH-adjusted $q$-value, and the overlapping miRNAs.

Supplementary Table 9: Putative target-program enrichment for top miRNA-specific contributors. For each highest-weight miRNA and each target program (keratinization, atypical/HPV, immune T-cell, EGFR ligand, EMT/stromal), the table reports the overlap size and mean target correlation used to summarize miRNA $\rightarrow$ program assignments in Supplementary Fig. 9c.

Supplementary Table 10: GO Biological Process enrichment of predicted targets of miRNA-specific leading miRNAs. The table reports the top enriched terms with gene-ratio, background ratio, raw $p$-value, BH-adjusted $p$-value, $q$-value, and count.

Supplementary Table 11: miRNA-specific partition Cox summary. Likelihood-ratio comparison of nested models with and without the miRNA-specific cluster assignment added to the baseline (shared cluster, age, sex, HPV status, stage, anatomic subsite, and smoking status): log-likelihood, AIC, $C$-index, and LRT $p$-value. The miRNA-specific partition improved the C-index from $C = 0.624$ to $C = 0.652$ (LRT $p = 4.9 \times 10^{-3}$) in the full clinical complete-case subset.

Supplementary Table 12: HNSC shared-clustering method benchmark (full). Cox $C$-index, log-rank $p$-value, adjusted Rand index against the Walter 4-subtype reference, normalized mutual information, and Cramér's $V$ for each of the five evaluated methods (multiRF, SNF, MOFA2, iClusterPlus, AJIVE). This table provides the numerical values summarized graphically in Supplementary Fig. 13.

Supplementary Table 13: Nested Cox models for overall survival. Incremental $C$-index and likelihood-ratio $p$-values for nested models (full clinical baseline $\rightarrow$ + shared cluster $\rightarrow$ + gene EV1 $\rightarrow$ + miRNA EV1 and methylation EV1). The full clinical baseline includes age, sex, HPV status, stage, anatomic subsite, and smoking status.

Supplementary Table 14: HNSC EV1 Cox sensitivity models with tumor purity, ESTIMATE immune score, and ESTIMATE stromal score. Sheet 14a reports the Table 4 full clinical Cox model after adding one microenvironmental covariate at a time; for each model, the table extracts the adjusted hazard ratio for gene EV1, miRNA EV1, and methylation EV1. Sheet 14b reports simpler EV-plus-composition models, in which each EV is modeled with one microenvironmental covariate at a time. Together, the two sheets distinguish whether attenuation is seen in the full clinical model or already in a direct EV-composition sensitivity model.

Supplementary Table 15: Progression-free interval (PFI) Cox models for the multiRF shared clusters. Cluster hazard ratios and sensitivity models mirroring the OS analysis in Table 4.

Supplementary Table 16: Covariate associations with multiRF shared clusters. Kruskal–Wallis tests for continuous variables and chi-squared tests for categorical variables. CSF biomarkers are reported on the subset of participants with available lumbar puncture data. Significance codes: *** $p < 0.001$, ** $p < 0.01$, * $p < 0.05$.

Supplementary Table 17: Cox regression coefficients for the multiRF shared cluster model. The model includes cluster membership (reference: Cluster 1), age, sex, education, APOE $\varepsilon 4$, and MMSE as covariates; $n = 87$, 24 conversion events, model $C$-index = 0.662.

Supplementary Table 18: ADNI direct-coordinate Cox models, incremental value over clinical covariates. For each learned representation, the baseline "covariates-only" model (age, sex, education, MMSE, APOE $\varepsilon 4$) is compared with a full model that adds the signal as a continuous set of coordinate predictors. The table also reports the coordinates-only model for the same representation. $C_{\text{coords}}$ and $C_{\text{full}}$ denote the Cox C-indices of the coordinates-only and covariates-plus-signal models, AIC gives each model's information criterion, and LRT $p$ is the likelihood-ratio test for the added signal over the covariates-only baseline ($C_{\text{cov}} = 0.623$ on this cohort). Methods are sorted by $C_{\text{full}}$.

Supplementary Table 19: Ablation study, sensitivity of multiRF to CpG selection strategy. Three CpG sets were evaluated: an AD-prioritized set (3,503 CpGs from prior EWAS), a brain–blood cross-tissue set (831 CpGs), and their union (4,332 CpGs after counting two overlapping CpGs once). For each set, multiRF was run 10 times with different random seeds. Reported values are mean ± SD across runs for age $R^2$, adjusted hazard ratio per one-year increase in age acceleration for MCI/AD conversion, and Cox C-index. DNAm-specific rows are reported; MRI-specific residual rows were excluded because numerical HR estimates were unstable at this sample size (for reference, raw HR means ranged over 4–16 orders of magnitude across the three CpG sets).

Supplementary Table 20: Quadrant analysis of modality-specific age acceleration. Participants were stratified into four groups by the sign of DNAm-specific and MRI-specific age acceleration relative to the cohort median, and adjusted Cox models were fit with the both-accelerated group as reference, adjusting for age, sex, education, MMSE, and APOE $\varepsilon 4$ ($n = 87$, 24 events). The both-decelerated cell contained no conversion events, so its Cox point estimate and confidence interval are numerically unstable and are reported as "unstable"; the log-rank contrast of the two extremes ($p = 0.0057$) in Supplementary Fig. 17 is the stable summary for that cell.

Supplementary Table 21: Per-EV covariate and CSF-biomarker associations for multiRF signals. Pearson correlation ($r$) for continuous covariates (age, MMSE) and Welch $t$-test $p$-value for binary covariates (sex, APOE $\varepsilon 4$); Spearman correlation ($\rho$) for CSF biomarkers (A$\beta_{42}$, phospho-tau). Demographic correlations are computed on $n = 87$ and CSF associations on the lumbar-puncture subset (A$\beta_{42}$ $n = 21$; p-tau $n = 38$). Asterisks denote $p < 0.05$. EVs are indexed by descending positive eigenvalue of the double-centered similarity matrix.

* $p < 0.05$. CSF significant cells: shared EV5 vs. A$\beta_{42}$ ($p = 0.029$); DNAm-specific EV5 vs. A$\beta_{42}$ ($p = 0.018$); MRI-specific EV1 vs. p-tau ($p = 0.047$); MRI-specific EV2 vs. p-tau ($p = 0.045$).

Supplementary Table 22: Cross-method cluster comparison on ADNI. Adjusted Cox hazard ratios for Cluster $c$ versus Cluster 1 (reference) in $k = 3$ spectral partitions from four integration methods (multiRF shared, SNF fused, MOFA2 shared, AJIVE joint) and the two multiRF modality-specific signals. Models are adjusted for age, sex, education, MMSE, and APOE $\varepsilon 4$ ($n = 87$, 24 events). Model $C$-index is reported once per method as it is invariant across contrasts within the same fit.

Supplementary Table 23: Embedding-dimension sweep for the continuous biological-age pipeline. For each method we refit the biological age regression and adjusted Cox model at $k \in \{3, 5, 10\}$ EVs and report the age $R^2$, adjusted hazard ratio per one-year increase in age acceleration, in-sample $C$-index ($C_{\text{in}}$), and downstream leave-one-out cross-validated $C$-index after fixed representation learning ($C_{\text{cv}}$). MRI-specific hazard ratios are numerically unstable at small $k$ and are omitted.

Supplementary Table 24: ADNI all-DNAm sensitivity analysis: DNAm-only biological-age signals across the expanded cognitively normal cohort. Cohort comprises all baseline ADNI CN participants with available DNAm and follow-up diagnosis ($n = 213$, 64 conversions to MCI or dementia), regardless of CSF amyloid/tau status. The main-text analysis was restricted to the A−T− preclinical subset ($n = 87$); this sensitivity test asks whether the DNAm-only signals would replicate in the broader CN cohort. For each method we report the number of input CpGs, the chronological-age $R^2$ of the biological-age estimate, the adjusted Cox hazard ratio per one-year and per one-SD increase in age acceleration (clinical clocks: difference between predicted and chronological age; DunedinPACE: pace of aging), the $p$-value for the age-acceleration coefficient, and the model concordance index. All Cox models adjust for age, sex, education, baseline MMSE, and APOE $\varepsilon 4$ carrier status.
